# Multi-objective Evaluation and Optimization of Stochastic Gradient Boosting Machines for Genomic Prediction and Selection in Wheat (*Triticum aestivum*) Breeding

**DOI:** 10.1101/2025.07.24.665873

**Authors:** Henry Munroe, Bright Osatohanmwen, Reza Sharifi

**Author notes:** Corresponding author: Reza Sharifi.

## Abstract

Machine learning (ML) models with stochastic and non-deterministic characteristics are increasingly used for genomic prediction in plant breeding, but evaluation often neglects important aspects like prediction stability and ranking performance. This study addresses this gap by evaluating how two hyperparameters of a Gradient Boosting Machine (GBM), learning rate (v) and boosting rounds (ntrees), impact stability and multi-objective predictive performance for cross-season, cross-environment prediction in a MAGIC wheat population. Using a grid search of 36 parameter combinations, we evaluated four agronomic traits with five metrics: Pearson’s r, Area Under the Curve (AUC), Normalized Discounted Cumulative Gain (NDCG), and the Intraclass Correlation Coefficient (ICC) and Fleiss’ κ for stability. Our findings show that a low learning rate combined with a high number of boosting rounds substantially improves prediction stability (ICC > 0.98) and selection stability (Fleiss’ κ > 0.80), while reducing train-test performance gaps. This combination produced concurrent improvements for predictive accuracy (r) and ranking efficiency (NDCG), though optimal settings were trait-dependent. Conversely, classification accuracy (AUC) was poor and performed relatively better with higher learning rates, revealing a conflict in optimization hyperparameters. Despite moderate Pearson’s r and poor AUC in this challenging cross-season, cross-environment prediction scenario, NDCG remained high (> 0.85), indicating strong ability to rank top-performing entries. Ultimately, prioritizing stability when tuning GBMs effectively yields reproducible cross-environment predictions with improved accuracy and top-end ranking performance.

## Introduction

As machine learning (ML) becomes more prevalent in plant breeding, breeders face increasing demands from large genomic and phenotypic datasets, computational time constraints and pressing market and environmental challenges (Pieruschka and Schurr 2019; van Dijk et al. 2020; Calvin et al. 2023; Yoosefzadeh Najafabadi et al. 2023; Crossa et al. 2025). The effectiveness of ML models depends on how they are constructed, optimized, and evaluated (Domingos 2012; Feurer and Hutter 2019). Model evaluation often relies on a limited range of metrics emphasizing overall predictive accuracy, via Pearson’s r (Meuwissen et al. 2001; Goddard and Hayes 2007). Although overall predictive accuracy is fundamental to assessing model performance, relying solely on accuracy metrics to evaluate ranking efficiency, selection efficiency, and stability, is inherently limited; this creates an evaluation gap and underscores the need for targeted metrics (Blondel et al. 2015; Dekkers et al. 2021; Lange et al. 2025).

A multi-metric evaluation is therefore justified to address this gap. Ranking ability can be assessed with Normalized Discounted Cumulative Gain (NDCG), which rewards the correct ordering of top-performing candidates (Järvelin and Kekäläinen 2002; Blondel et al. 2015). For binary “select/reject” decisions, the Area Under the Curve (AUC) measures a model’s ability to distinguish between classes (Huang and Ling 2005; Robin et al. 2011). Yet, neither metric directly addresses the issue of non-determinism, or stochasticity, inherent in many algorithms.

In stochastic Gradient Boosting Machines (GBM), random subsampling introduces variability between runs (Friedman 2002). This requires explicit stability assessment. The Intraclass Correlation Coefficient (ICC) can quantify the consistency of continuous predictions across iterations (Bartko 1966; McGraw and Wong 1996; Weir 2005; Koo and Li 2016; Lange et al. 2025). A high ICC value indicates that variability in predictions is mainly due to inherent differences among targets rather than random fluctuations, demonstrating stable model performance (Weir 2005). For categorical selections, Fleiss kappa (κ) measures the stability of agreement among “candidate” classifications across runs (Fleiss 1971; Lange et al. 2025).

GBM performance is influenced by hyperparameters that balance the bias-variance trade-off. The learning rate and the number of boosting rounds must be carefully tuned to capture data complexity without overfitting (Friedman 2001; Ridgeway and Developers 2003). This creates a multi-objective optimization problem: maximizing predictive, selection and ranking accuracy while ensuring high stability.

This study performs a parametric optimization of a stochastic GBM to address these challenges. We systematically evaluate 36 combinations of learning rate and boosting rounds using genomic and phenotypic data from a Multi-parent Advanced Generation Inter-Cross (MAGIC) wheat population. Cross-season predictions are assessed on four traits-Grain Yield (GY), Grain Protein Content (GPC), Thousand Grain Weight (TGW), and Height to Ear tip (HET). We aim to: (1) evaluate performance using Pearson’s r, NDCG, and AUC; (2) quantify prediction and selection stability with ICC and Fleiss κ; (3) determine the effects of hyperparameters on all metrics; and (4) provide insights into model generalizability. Findings will help breeders select ML models that are not only accurate, but also stable enough to support confident, repeatable selection decisions.

## Methods and Materials

A 500-line wheat (*Triticum aestivum*) Multi-parent Advanced Generation Inter-Cross (MAGIC) population was analyzed (Scott et al. 2021). Four agronomic traits were assessed from their study: GY, TGW, GPC, HET in a cross-season prediction scenario. Field experiments were conducted at Girton, UK (52.24144 °N, 0.08794 °E; clay-loam soil; 2016-17 season, training) and Whittlesford, UK (52.09909 °N, 0.13311 °E; loam soil; 2017-18 season, testing) (Scott et al. 2021).

### SNP genotyping and quality control

Genotypes comprised 25,508 single-nucleotide polymorphisms (SNPs) from the Axiom 35k wheat breeders’ SNP genotyping array (Winfield et al. 2016). Loci with > 5 % missing calls were excluded. Remaining missing values were imputed by Random Forest using the missRanger R package (v2.5.0, see File S1 - Package_Functions) (Mayer 2024).

### Train-test splits

To evaluate performance in a cross-environment prediction scenario, models were trained using data from the 2016/17 season and evaluated against observed phenotypes from the 2017/18 season. Five random 80%/20% partitions of the 500 lines were generated using set.seed(123).

### Gradient-boosting

Stochastic gradient-boosting machine was fitted with gbm v2.2.2 in R 4.4.3. A 6 × 6 tune grid, set manually with; learning rate = {0.0025, 0.042, 0.0815, 0.121, 0.1605, 0.20} and boosting rounds = {100, 1080, 2060, 3040, 4020, 5000}, while all other arguments remained at default (including bag.fraction = 0.5) (File S1 - Package_Functions). (DOI: 10.5281/zenodo.16785772 - Data Availability).

The ensemble update at boost round *m* followed (Friedman 2002; Ridgeway and Developers 2003):

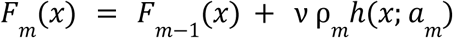

Where:

*F_m_*(*x*) is the updated ensemble prediction
*F_m_*_-1_(*x*) is the prediction from the previous iteration
*ρ_m_* is the optimal step length determined via line search
*h*(*x*:*a_m_*) is a regression tree fit to the pseudo residuals
*v* is the learning rate (shrinkage, 0 < v ≤ 1)

### Prediction pipeline

For each of the five train-test splits, each of the ten stochastic model fits was independently used to generate predictions on its corresponding 20% hold-out set. This yielded 50 distinct prediction vectors (5 splits × 10 iterations per split) for each hyperparameter combination, which were then compared to the observed phenotypes. Reproducibility ensured via set.seed(123 + split_number × 1000 + prediction_iteration) (DOI: 10.5281/zenodo.16785772 - Data Availability). See Fig. 1 for the schematic diagram of the pipeline.

**Figure 1.**
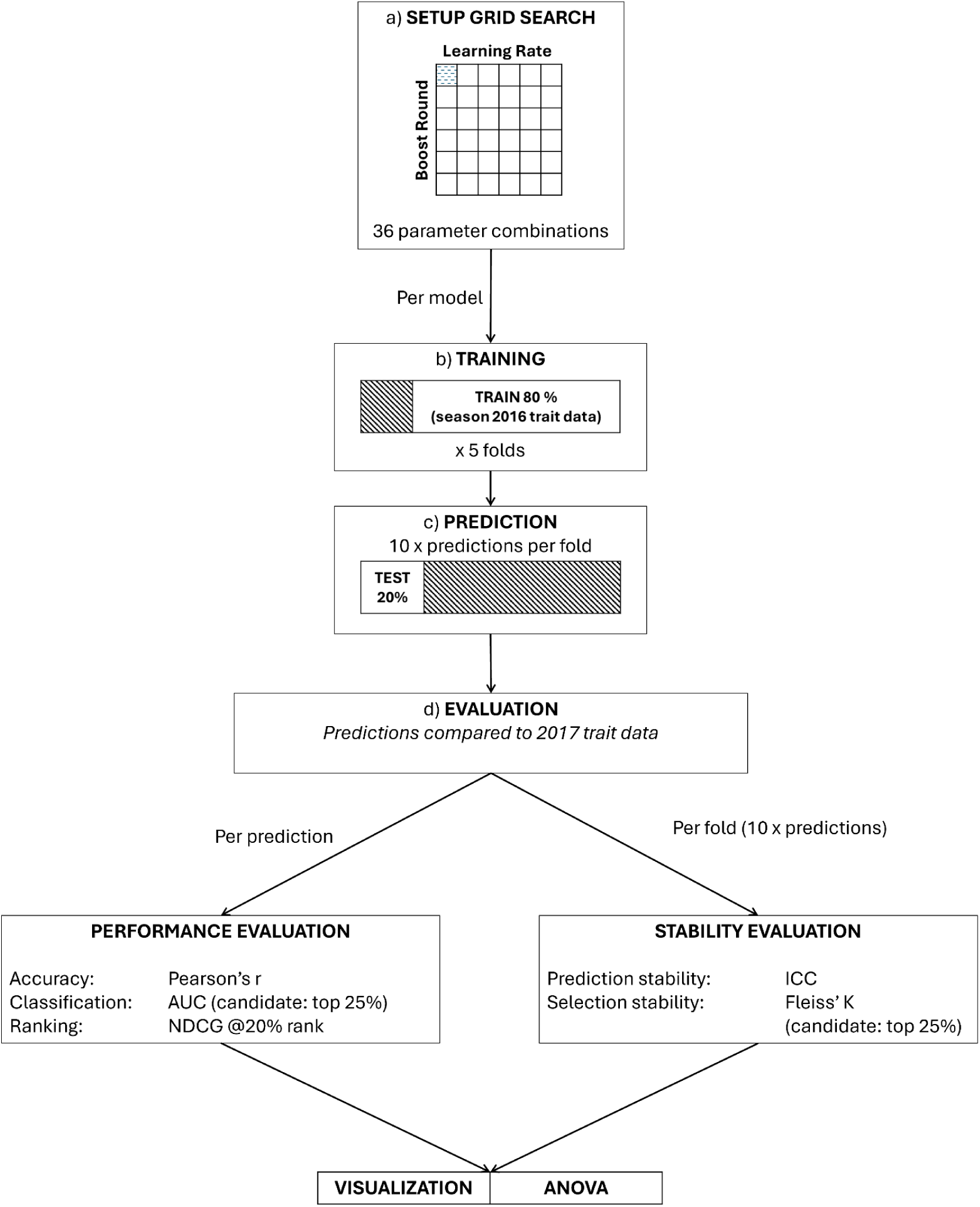
shows the pipeline (per trait): (a) grid search setup, where 36 tuning parameter combinations (6 learning rates x 6 boosting rounds) are specified; (b) training, where each combination is trained on 80 % of entries using the 2016 trait data with five-fold cross-validation; (c) prediction, in which ten iterations of predictions are produced per fold on the hold out 20 % test set; and (d) evaluation, where predictions are compared against 2017 trait values and divided into performance and stability evaluation performance metrics (Pearson’s r, AUC for the top 25 % candidates, and NDCG at the top 20 % rank) are calculated for each prediction, and stability (Intraclass correlation coefficient and Fleiss’ κ) is assessed per fold. All metrics are then plotted and analyzed via type 3 ANOVA.

### Performance and stability metrics

Performance metrics, Accuracy - Pearson’s r, ranking efficiency - Normalized Discounted Cumulative Gain for the top 20% of predicted lines (NDCG@20%), and classification ability - by labeling lines with observed phenotypes above the 75th percentile as Candidates and then computing the ROC AUC (File S1 - Formulae). Stability across the ten stochastic repeats was quantified by the single-measure, one-way intraclass correlation coefficient (ICC) for predictions and by Fleiss K for overall agreement of binary selection (File S1 - Formulae, DOI: 10.5281/zenodo.16785772 - Data Availability). Across the hyperparameter grid, each trait yielded n = 1,800 performance estimates (r, NDCG@20 %, AUC; 5 splits × 6 learning-rates × 6 boost-rounds 10 iterations) and n = 180 stability estimates (ICC, Fleiss K; 5 splits × 6 learning-rates × 6 boost-rounds) (File S4).

### Data visualization

Figures were generated with ggplot2 3.4 and viridis 0.6. Error bars are the standard error (± SE) which are found in the table of summary statistics (mean, SD, SE, min, max, IQR) for every metric and hyperparameter combination in File S2. Min-max normalization of performance and stability metrics were applied and plotted as supplementary material (Fig. S2).

### Statistical analysis

All statistics were run in R 4.4.3 to quantify how GBM hyperparameters influence predictive performance and stability within and across traits. Q-Q and residual-versus-fitted plots were made for observing normality of residuals (Fig. S4). Effect sizes were calculated as partial *η*^2^ (*η*^2^_p_) for all Type III ANOVA Satterthwaite-approximated F-tests and as Kendall’s W for Friedman tests, using the effectsize R package (v1.01).

### Modelling framework

Trait-wise predictive performance was analyzed with random-intercept linear mixed models (LMM; lme4 v1.1-36), using learning rate and boosting rounds as fixed effects and random intercepts to account for Subset and Subset:Iteration nesting. Trait-wise stability was assessed non-parametrically via Friedman tests. To examine hyperparameter effects across traits, we fitted three-way LMMs (Trait × Learning Rate × Boost Rounds) with random intercepts for Trait and Subset (and, for performance only, Subset:Iteration). Cross-trait (aggregated) performance and stability were evaluated with two-way LMMs (Learning Rate × Boost Rounds), where performance models again included Subset:Iteration, but stability models retained only Trait and Subset nesting. Fixed effects were tested by, Type III, Satterthwaite-approximated F-tests (lmerTest v3.1-3), Tukey-Kramer (Tukey-adjusted pairwise) tests were made via emmeans v1.11.0. All p-values reported are raw unless otherwise specified, with post-hoc contrasts presented as Tukey-adjusted p-values. (DOI: 10.5281/zenodo.16785772 - Data Availability, File S3).

## Results

Analysis of raw agronomic data of the 500 entries across two seasons found that some traits were more stable year-to-year than others. TGW and HET showed strong Pearson’s r between years (r = 0.83 and r = 0.93, respectively). In contrast, GY showed a weak cross-year correlation (r = 0.38), whereas GPC was moderate (r = 0.61). The mean values for GPC also varied largely between the first year (13.09% SE = 0.03) and the second year (9.47% SE = 0.03), unlike other traits (Fig. 2).

**Figure 2.**
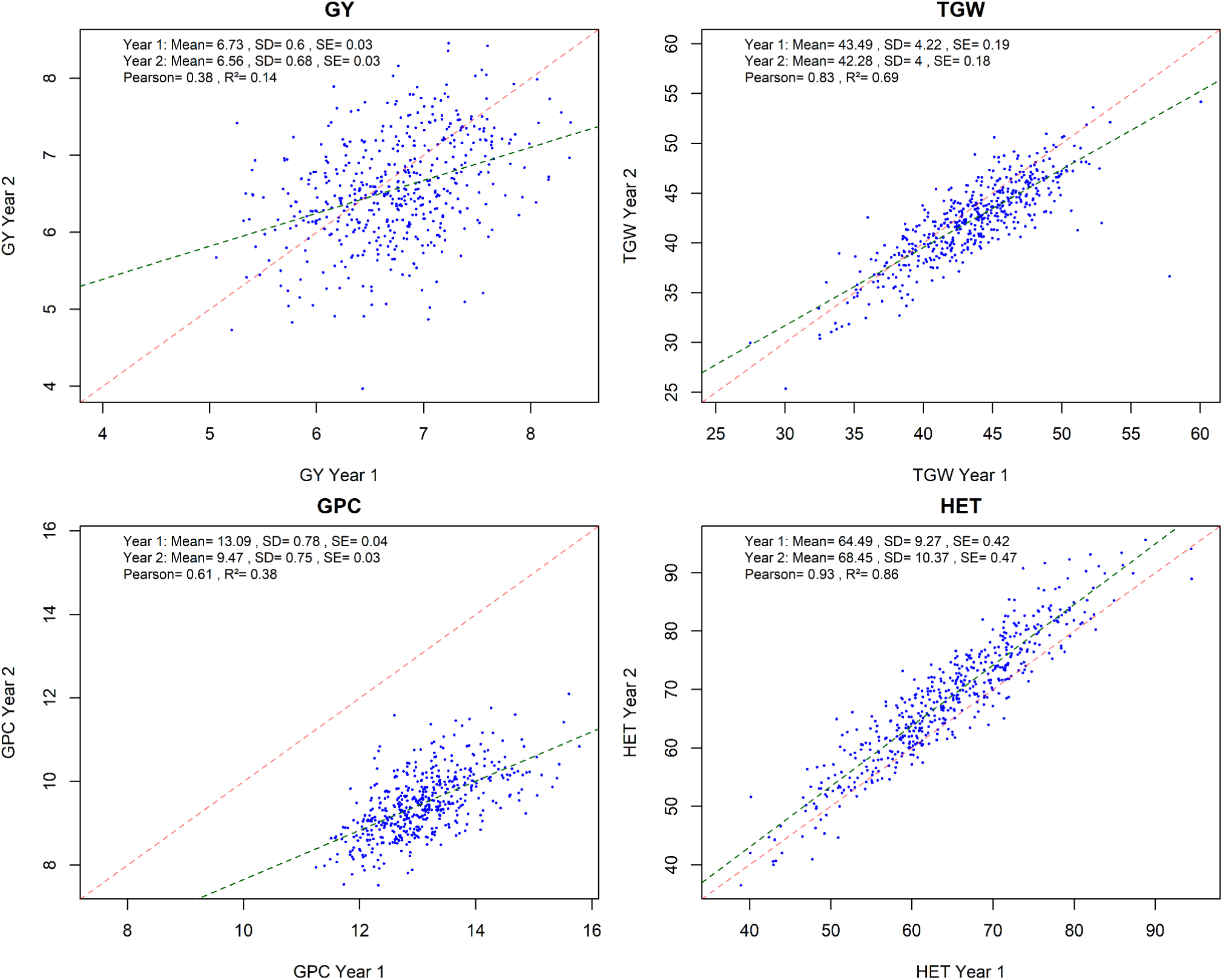
This figure displays four scatter plots of observed measurements, shows the relationship between Year 1 (x-axis) and Year 2 (y-axis) for four agronomic traits from the wheat dataset. Traits: GY (grain yield) - top left, TGW (thousand grain weight) - top right, GPC (grain protein content) - bottom left and HET (height to ear tip) - bottom right. Each blue point represents an individual wheat line’s measurements in both years. The red diagonal line is the 45° (theoretical) perfect agreement between the two years. The green dashed line is the best-fit regression line. The annotations in each panel summarize the mean, standard deviation (SD), and standard error (SE) for both years, Pearson’s r and R².

To determine the optimal hyperparameters for genomic prediction, we evaluated the effects of learning rate and boosting rounds on overall model performance and stability by analyzing metrics aggregated across all four traits. Elapsed time was not affected by learning rate but increased linearly with boosting rounds: models with 100 rounds finished in under 30 minutes per core, while 5,000 boost rounds required ≈ 10 hours per core (Fig. S3).

The aggregated performance of the 36 hyperparameter combinations across all traits is presented in Figure 3. Models combining low learning rates (v = 0.0025 to 0.0815) with moderate-to-high boosting rounds (> 2000) achieved the highest ICC, Fleiss’ κ, Pearson’s r and NDCG@20% (Fig. 3a-c, 3e). Predictive accuracy (Pearson’s r) and ranking performance (NDCG@20 %) showed moderate improvements, while substantial improvements in stability metrics (ICC and Fleiss’ κ) were observed (Fig. 3a-c, 3e). A Type III, three-way ANOVA, with fixed-effect significance determined by Satterthwaite-approximated F-tests, showed that the interaction of trait × learning rate × boosting rounds was a significant source of variation for all three performance metrics: Pearson’s r (F_(75, 6860)_ = 5.06, *η*^2^_p_ = 0.052, *p* < 0.001), AUC (F_(75, 6860)_ = 1.95, *η*^2^_p_ = 0.020, *p* < 0.001), and NDCG@20% (F_(75, 6860)_ = 2.60, *η*^2^_p_ = 0.028, *p* < 0.001). This indicates that optimal hyperparameter settings were trait-dependent. The two-way interaction between learning rate × boosting rounds was also significant across all three metrics: Pearson’s r (F_(25, 6860)_ = 22.18, *η*^2^_p_ = 0.075, *p* < 0.001), AUC (F_(25, 6860)_ = 8.26, *η*^2^_p_ = 0.029, *p* < 0.001), NDCG@20% (F_(25, 6860)_ = 1.71, *η*^2^_p_ = 0.006, *p* < 0.001) (File S3 - ANOVA 3-way).

**Figure 3.**
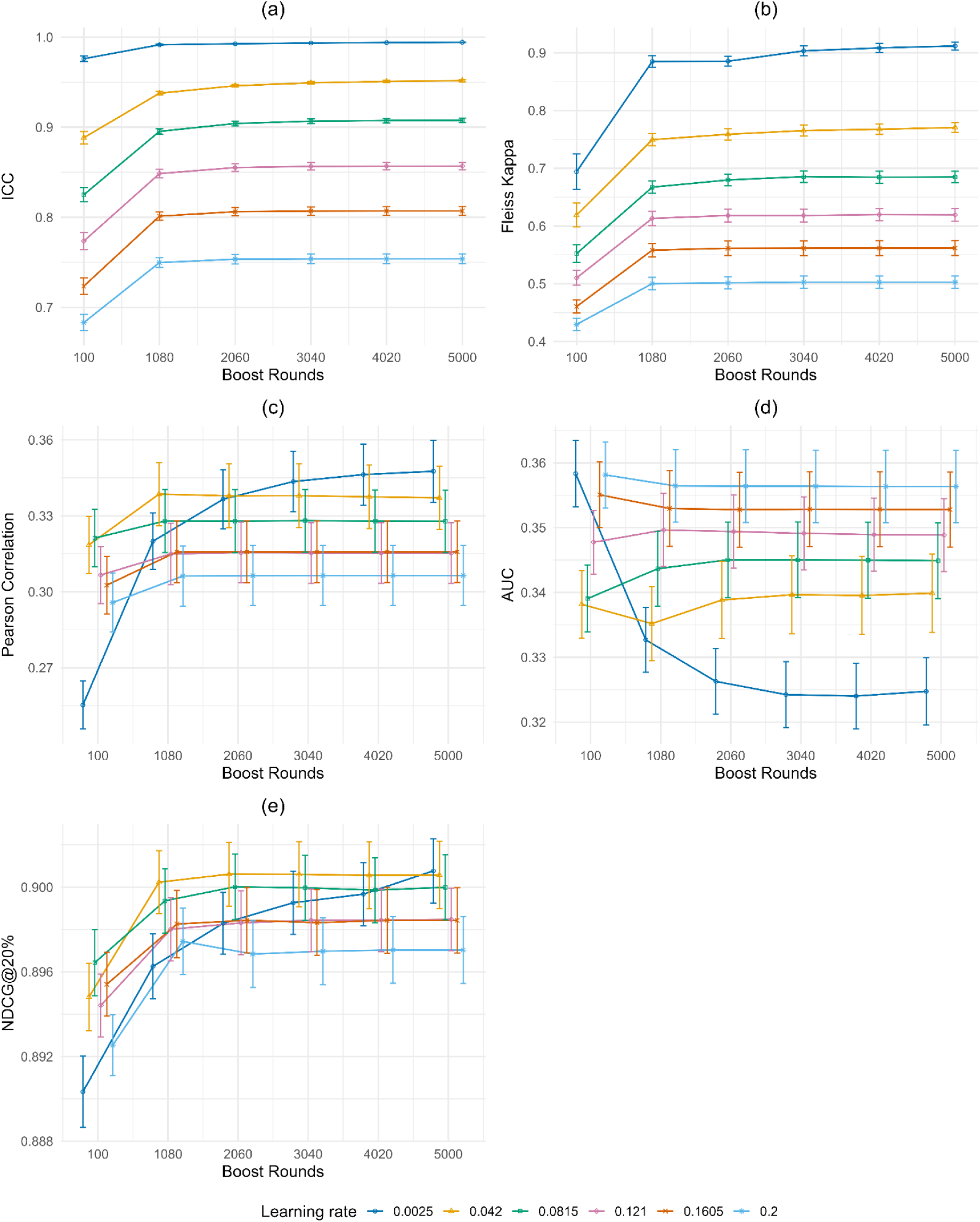
Displays trait aggregated values for five performance metrics: Intra-class Correlation Coefficient (ICC, top left), Fleiss’ κ (top right), Pearson’s r (center left), Area Under the Curve (AUC, center right), and Normalized Discounted Cumulative Gain at the top 20% ranking (NDCG@20%, bottom left). Each plot shows aggregated results across all four traits, with the x-axis representing boosting rounds and the y-axis indicating the respective evaluation metric. Colored lines connect the mean metric values for each learning rate, and error bars reflect ±1 standard error. This figure provides a view of model performance across all metrics. Each color corresponds to a distinct Learning Rate; Dark Blue (0.0025), Orange (0.042), Green (0.0815), Purple (0.121), Red-Orange (0.1605), and Light Blue (0.2).

Pairwise comparisons of the estimated marginal means (Tukey-adjusted) for the aggregated results identified two distinct performance trends. For Pearson’s r and NDCG@20%, models with lower learning rates (v = 0.0025 to 0.0815) combined with a moderate-to-high number of boosting rounds (ntrees > 2000) significantly outperformed models with higher learning rates (v > 0.1) (*p* < 0.001) (Fig. 3c, 3e). Within the low learning-rate group, at ≥ 2000 boosting rounds, the estimated marginal means for Pearson’s r and NDCG did not differ between v = 0.0025 and v = 0.042 (e.g Pearson’s r: Δ = −0.00134, SE = 0.004, t(6965) = −0.323, *p* > 0.99) (File S3 - Pairwise_overallmodel_Adj_Pears). In contrast, classification accuracy (AUC) found models with low boosting rounds (ntrees = 100) and higher learning rates (v > 0.01) consistently and significantly outperformed those with lower learning rates (v < 0.01) (*p* < 0.001) (Fig. 3d, File S3 - Pairwise_overallmodel_Adj_AUC).

The type III three-way ANOVA showed that hyperparameters, learning rate and boosting rounds had significant main effects on all performance metrics (r, AUC, NDCG@20%) although the magnitude of their effects varied by metric (File S3 - ANOVA 3-way). For Pearson’s r, both the learning rate and boosting rounds showed moderate, statistically significant effects: learning rate (F_(5, 6860)_ = 97.28, *η*^2^_p_ = 0.066, *p* < 0.001) and boosting rounds (F_(5, 6860)_ = 78.00, *η*^2^_p_ = 0.054, *p* < 0.001). For classification performance (AUC) the learning rate was the main driver (F_(5, 6860)_ = 128.73, *η*^2^_p_ = 0.086, *p* < 0.001) with boosting rounds provided significant but low effect (F_(5, 6860)_ = 5.55, *η*^2^_p_ = 0.004, *p* < 0.001). Ranking performance (NDCG@20%) showed boosting rounds as the main driver (F_(5, 6860)_ = 38.74, *η*^2^_p_ = 0.027, *p* < 0.001) compared to learning rate (F_(5, 6860)_ = 14.23, *η*^2^_p_ = 0.010, *p* < 0.001) (File S3 - ANOVA 3-way).

The effect of hyperparameter tuning for model stability showed substantial improvements (Figure 3a, 3b). A Type III, three-way ANOVA showed that the trait × learning rate × boosting rounds interaction was not significant for either stability metric (ICC: F_(75, 560)_ = 0.59, *η*^2^_p_ = 0.073, *p* = 0.99, Fleiss’ κ: F_(75, 560)_ = 1.21, *η*^2^_p_ = 0.140, *p* = 0.11), indicating consistent stability tuning effects across traits. Learning rate made the greatest contribution to stability (ICC: F_(5, 560)_ = 7304.27, *η*^2^_p_ = 0.985, *p* < 0.001, Fleiss’ κ: F_(5, 560)_ = 1793.30, *η*^2^_p_ = 0.941, *p* < 0.001) followed by boosting rounds with substantial secondary effects (ICC: F_(5, 560)_ = 602.76, *η*^2^_p_ = 0.843, *p* < 0.001, Fleiss’ κ: F_(5, 560)_ = 257.11, *η*^2^_p_ = 0.697, *p* < 0.001) (Table S3 - ANOVA 3-way). Significant interaction of learning rate × boosting rounds confirm that joint, co-optimization provide large effects in improvements to stability (ICC: F_(25, 560)_ = 15.15, *η*^2^_p_ = 0.404, *p* < 0.001, Fleiss’ κ: F_(25, 560)_ = 5.86, *η*^2^_p_ = 0.207, *p* < 0.001) (File S3 - ANOVA 3-way).

Pairwise comparisons of the estimated marginal means (Tukey-adjusted) confirmed a consistent trend: lower learning rates produced significantly higher stability scores for both ICC and Fleiss’ κ, with nearly all learning rate combinations being significantly different from one another when compared at the same boosting round level. For example, at 100 boosting rounds, the ICC score with a learning rate of 0.0025 was significantly higher than a rate of 0.042 (Mean Difference = 0.088, SE = 0.0056, t(677) = 15.76, *p* < 0.001). Full pairwise results are provided in File S3.

Model accuracy performance via Pearson’s r varied by trait. GPC showed the greatest increase in Pearson’s r at an intermediate number of boosting rounds (ntrees > 2000) when using learning rate v = 0.0025 (Fig. 4c). The mean Pearson’s r across hyperparameter combinations was 0.42 (SD = 0.091) for TGW and 0.51 (SD = 0.080) for HET (Fig. 4a-d, File S5). A Type III, two-way ANOVA for each trait confirmed that hyperparameter responses differed markedly among traits (File S3 - ANOVA Trait Specific 2-way). For GY, both learning rate (F_(5, 1715)_ = 2.34, *η*^2^_p_ = 0.007, *p* < 0.05) and boosting rounds (F_(5, 1715)_ = 3.78, *η*^2^_p_ = 0.011, *p* < 0.01) had small yet significant main effects, whereas their interaction was not significant (F = 0.63, *η*^2^_p_ = 0.009, *p* = 0.92). In TGW, both hyperparameters showed large, highly significant effects for learning rate (F_(5, 1715)_ = 61.39, *η*^2^_p_ = 0.152, *p* < 0.001) and boosting rounds (F = 98.87, *η*^2^_p_ = 0.224, *p* < 0.001) and the learning rate × boosting rounds interaction also contributed low but significant effect (F = 2.98, *η*^2^_p_ = 0.042, *p* < 0.001). Pearson’s r for GPC was driven primarily by learning rate (F = 58.94, *η*^2^_p_ = 0.147, *p* < 0.001) and boosting rounds showed a low but significant effect (F = 2.36, *η*^2^_p_ = 0.007, *p* < 0.05), and the interaction remained significant with low effect (F = 3.07, *η*^2^_p_ = 0.043, *p* < 0.001). HET displayed the strongest response to hyperparameters with large, significant main effects of learning rate (F = 77.71, *η*^2^_p_ = 0.185, *p* < 0.001) and boosting rounds (F = 113.42, *η*^2^_p_ = 0.249, *p* < 0.001), and a strong interaction effect (F = 38.82, *η*^2^_p_ = 0.361, *p* < 0.001). Full trait-specific ANOVA tables are provided in File S3 - ANOVA Trait-Specific 2-Way.

**Figure 4.**
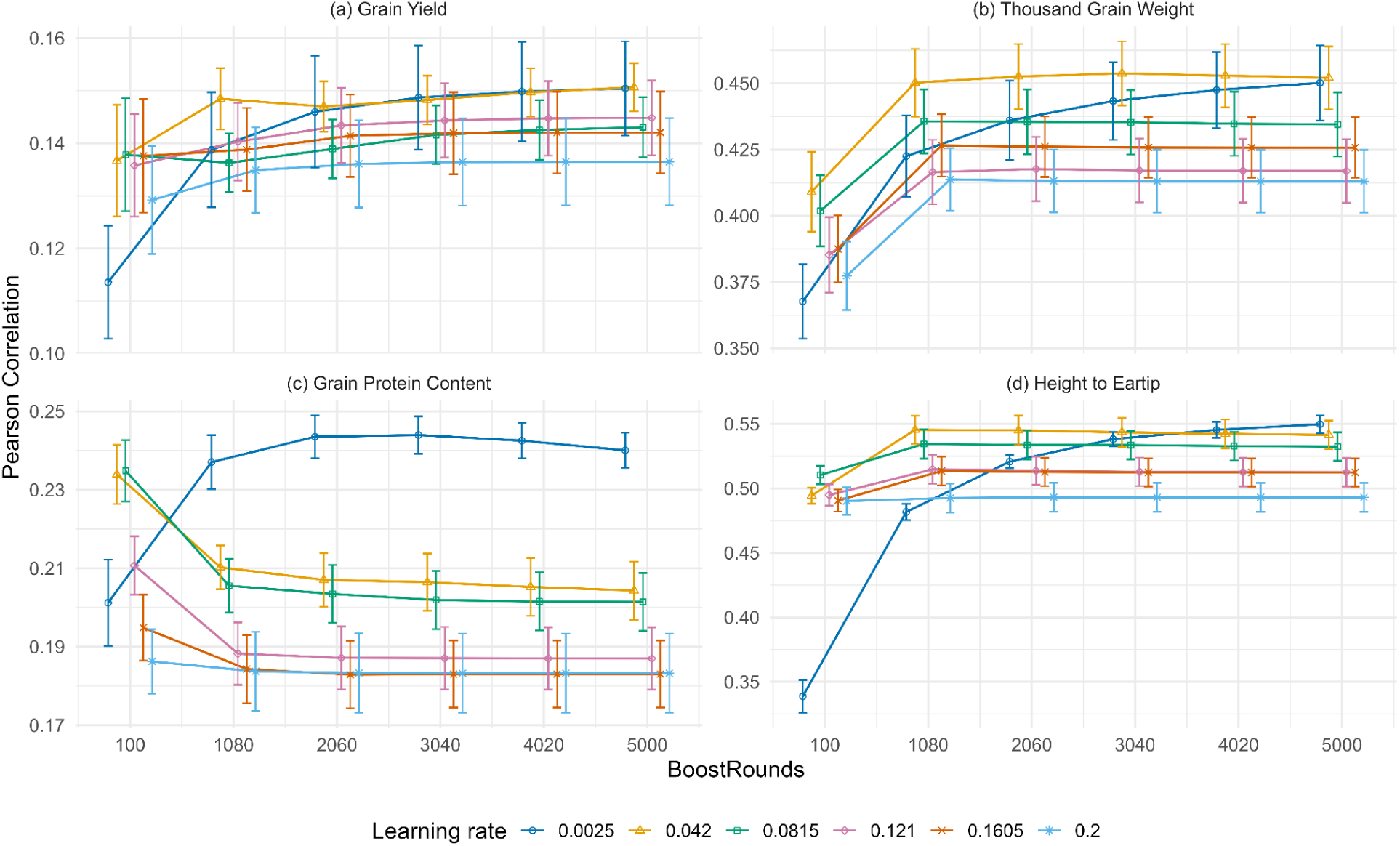
Displays the Pearson’s r coefficient varies with boosting rounds and learning rate across four traits Grain Yield (GY - top left), Thousand Grain Weight (TGW - top right), Grain Protein Content (GPC - bottom left), Height to Ear Tip (HET - bottom right). The x-axis represents boosting rounds and the y-axis indicates the mean Pearson’s r for each model combination, with error bars showing ±1 standard error. Coloured lines connect mean values for each learning rate. Each color corresponds to a distinct Learning Rate; Dark Blue (0.0025), Orange (0.042), Green (0.0815), Purple (0.121), Red-Orange (0.1605), and Light Blue (0.2).

Classification accuracy, measured by Area Under the Curve (AUC), was poor across all hyperparameter combinations, with values below the 0.5 threshold of random chance for all four traits (Fig. 5a-d). Visually, models with higher learning rates (v ≥ 0.042) resulted in higher AUC values, often peaking with 1000 - 2000 boosting rounds. In contrast, the lowest learning rate (v = 0.0025) consistently produced lower AUC values across all boosting rounds. With a learning rate of v = 0.0025, AUC decreased during moderate-to-high numbers of boosting rounds but began to increase from 4,000 boost rounds.

**Figure 5.**
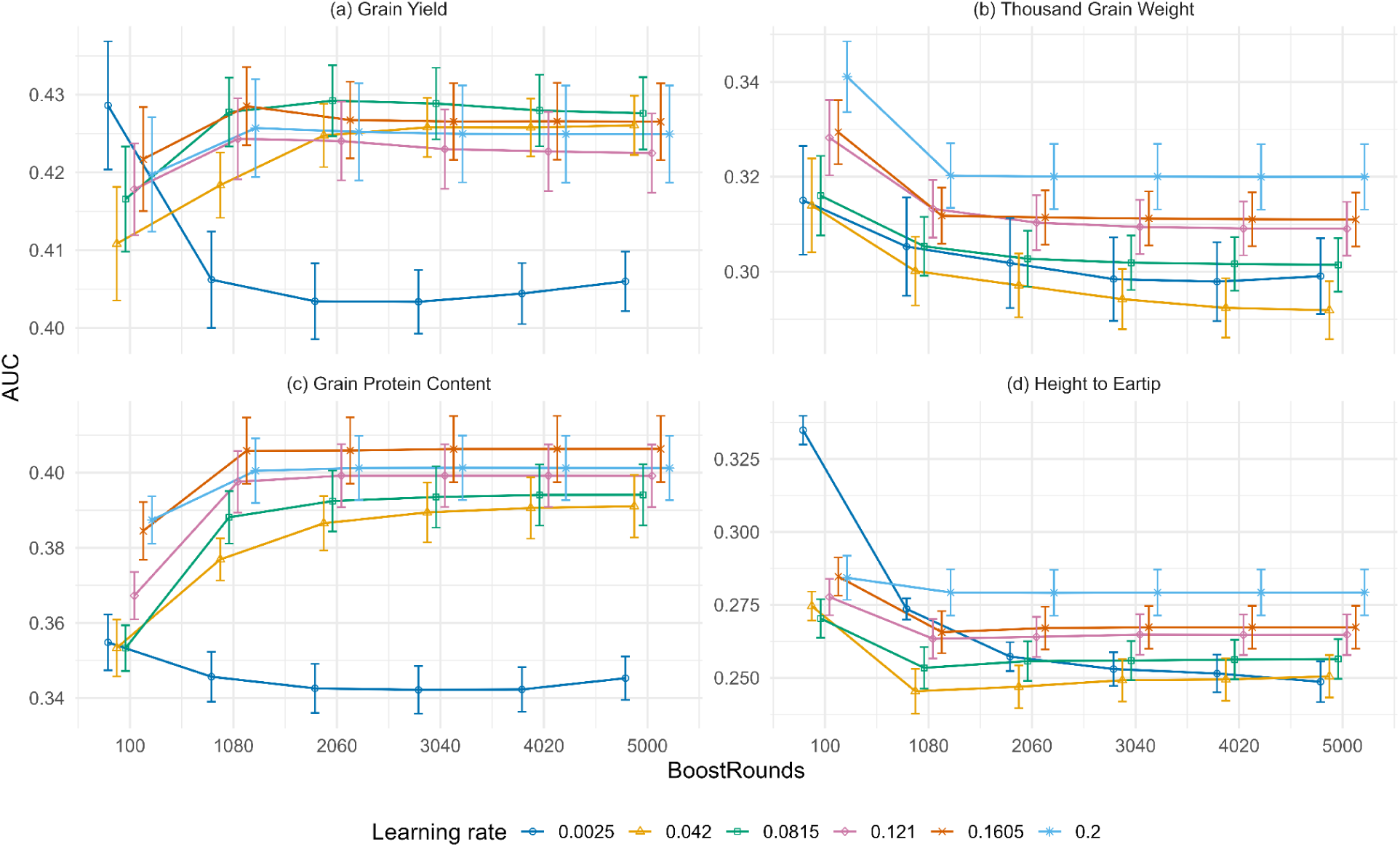
Displays Area Under the Curve (AUC) by boosting rounds across learning rate across four traits Grain Yield (GY - top left), Thousand Grain Weight (TGW - top right), Grain Protein Content (GPC - bottom left), Height to Ear Tip (HET - bottom right). The x-axis represents boosting rounds and the y-axis indicates the mean AUC for each model combination, with error bars showing ±1 standard error. Colored lines connect mean values for each learning rate. Data are presented in separate panels for each trait with free y-axis scales. Each color corresponds to a distinct Learning Rate; Dark Blue (0.0025), Orange (0.042), Green (0.0815), Purple (0.121), Red-Orange (0.1605), and Light Blue (0.2).

ANOVA for each trait confirmed that AUC responses to hyperparameters varied among traits (File S3 - ANOVA Trait-Specific 2-Way). For GY, learning rate showed a significant effect (F_(5, 1715)_ = 16.41, *η*^2^_p_ = 0.046, *p* < 0.001), boosting rounds had no independent influence (F_(5, 1715)_ = 0.53, η²p = 0.002, *p* = 0.755), and the interaction was significant but small (F_(25, 1715)_ = 2.08, *η*^2^_p_ = 0.030, p < 0.01). In TGW, both learning rate (F_(5, 1715)_ = 41.65, *η*^2^_p_ = 0.108, *p* < 0.001) and boosting rounds (F = 25.56, *η*^2^_p_ = 0.070, *p* < 0.001) had moderate effects, whereas the interaction was not significant (F = 0.24, *η*^2^_p_ = 0.004, p > 0.99). For GPC, learning rate had the largest main effect across traits (F_(5, 1715)_ = 111.83, *η*^2^_p_ = 0.246, *p* < 0.001), boosting rounds also contributed (F = 20.31, *η*^2^_p_ = 0.056, *p* < 0.001), and a small but significant interaction was detected (F = 2.66, *η*^2^_p_ = 0.037, *p* < 0.001). HET displayed large, significant effects for learning rate (F_(5, 1715)_ = 40.22, *η*^2^_p_ = 0.105, *p* < 0.001) and boosting rounds (F = 48.09, *η*^2^_p_ = 0.123, *p* < 0.001), alongside a substantial interaction (F_(25, 1715)_ = 9.47, *η*^2^_p_ = 0.121, *p* < 0.001). Full ANOVA results are available in File S3 - ANOVA Trait-Specific 2-Way.

Ranking performance, measured by NDCG@20%, was high across all traits, with values consistently above 0.85 (Fig. 6, File S2, S4). Figure 6 shows that for the lowest learning rate (v = 0.0025), NDCG@20% values increased across all boosting rounds. In contrast, for higher learning rates, performance typically peaked with 1000-3000 boosting rounds before plateauing or declining (Fig. 6a-d).

**Figure 6.**
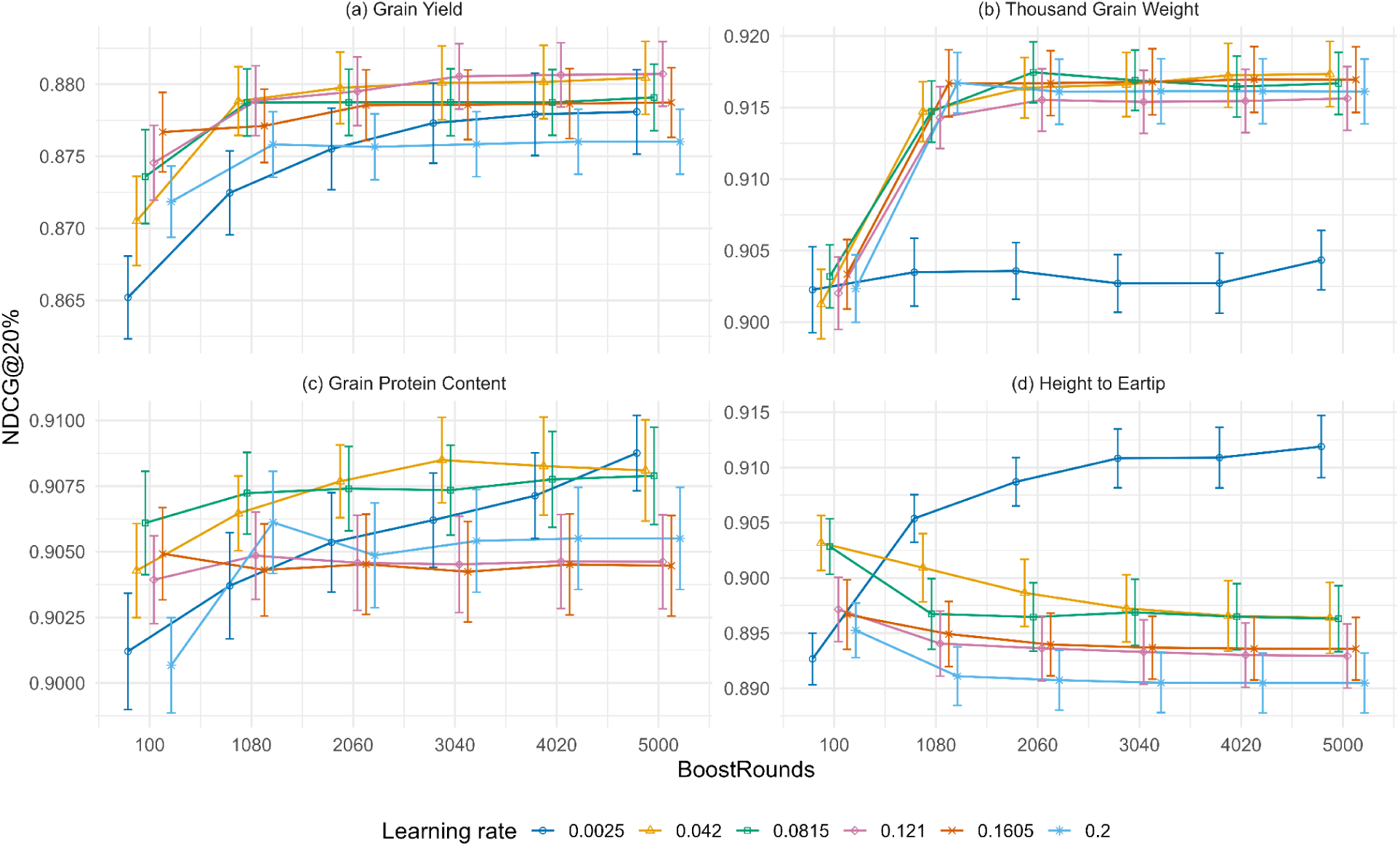
The plot displays Normalized Discounted Cumulative Gain (NDCG) evaluated at the cutoff corresponding to the top 20% position of ranked entries (denoted, NDCG@20%) compared to different combinations of boosting rounds and learning rate across four traits Grain Yield (GY - top left), Thousand Grain Weight (TGW - top right), Grain Protein Content (GPC - bottom left), Height to Ear Tip (HET - bottom right). The x-axis shows boosting rounds and the y-axis indicates the mean NDCG@20% for each model combination, with error bars showing ±1 standard error. Colored lines connect mean values for each learning rate. Data are presented in separate panels for each trait with free y-axis scales. Each color corresponds to a distinct Learning Rate; Dark Blue (0.0025), Orange (0.042), Green (0.0815), Purple (0.121), Red-Orange (0.1605), and Light Blue (0.2).

ANOVA for each trait confirmed that NDCG@20 % responses to hyperparameters varied among traits (File S3 - ANOVA Trait-Specific 2-Way). For GY, learning rate (F_(5, 1715)_ = 6.97, *η*^2^_p_ = 0.020, *p* < 0.001) and boosting rounds (F_(5, 1715)_ = 13.02, *η*^2^_p_ = 0.037, *p* < 0.001) had small but significant effects, while their interaction was not significant (F_(25, 1715)_ = 0.84, *η*^2^_p_ = 0.012, *p* = 0.70). In TGW, both learning rate (F = 50.75, *η*^2^_p_ = 0.129, *p* < 0.001) and boosting rounds (F = 59.49, *η*^2^_p_ = 0.148, *p* < 0.001) showed substantial effects, and a modest interaction was present (F = 2.12, *η*^2^_p_ = 0.030, *p* < 0.001). For GPC, learning rate (F = 6.05, *η*^2^_p_ = 0.017, *p* < 0.001) and boosting rounds (F = 4.14, *η*^2^_p_ = 0.012, *p* < 0.001) had small but significant effects, whereas the interaction was not significant (F = 0.82, *η*^2^_p_ = 0.012, *p* = 0.72). HET differed, showing a strong main effect of learning rate (F = 62.35, *η*^2^_p_ = 0.154, *p* < 0.001), no main effect of boosting rounds (F = 0.35, *η*^2^_p_ = 0.001, *p* = 0.88), and a moderate interaction (F = 5.27, *η*^2^_p_ = 0.071, *p* < 0.001). Full ANOVA results are provided in File S3 - ANOVA Trait-Specific 2-Way.

Prediction stability (ICC) and selection stability (Fleiss Kappa) showed highly consistent and clear trends across all four traits (Fig. 7, 8). For both metrics, stability values systematically increased as the learning rate decreased. The lowest learning rate (ν = 0.0025) consistently resulted in the highest stability, with ICC values approaching 1.0 for all traits at 5,000 boost rounds (M = 0.994, SD = 0.001, N = 20; File S4, S2). For any given learning rate, stability also increased with more boosting rounds, typically plateauing after approximately 2,000 rounds (Fig. 7, 8). Friedman tests confirmed significant differences among the 36 hyperparameter combinations for both stability metrics across all traits. For ICC, χ²(35) = 172.75 - 174.47, *p* < 0.001, with a strong level of agreement among the rankings (Kendall’s *W* = 0.987 - 0.997). For Fleiss’ κ, χ²(35) = 160.48 - 168.10, *p* < 0.001, with very strong agreement (Kendall’s *W* = 0.917 - 0.961). Full trait-specific tables are provided in File S3 - Trait Specific Friedman Stab.

**Figure 7.**
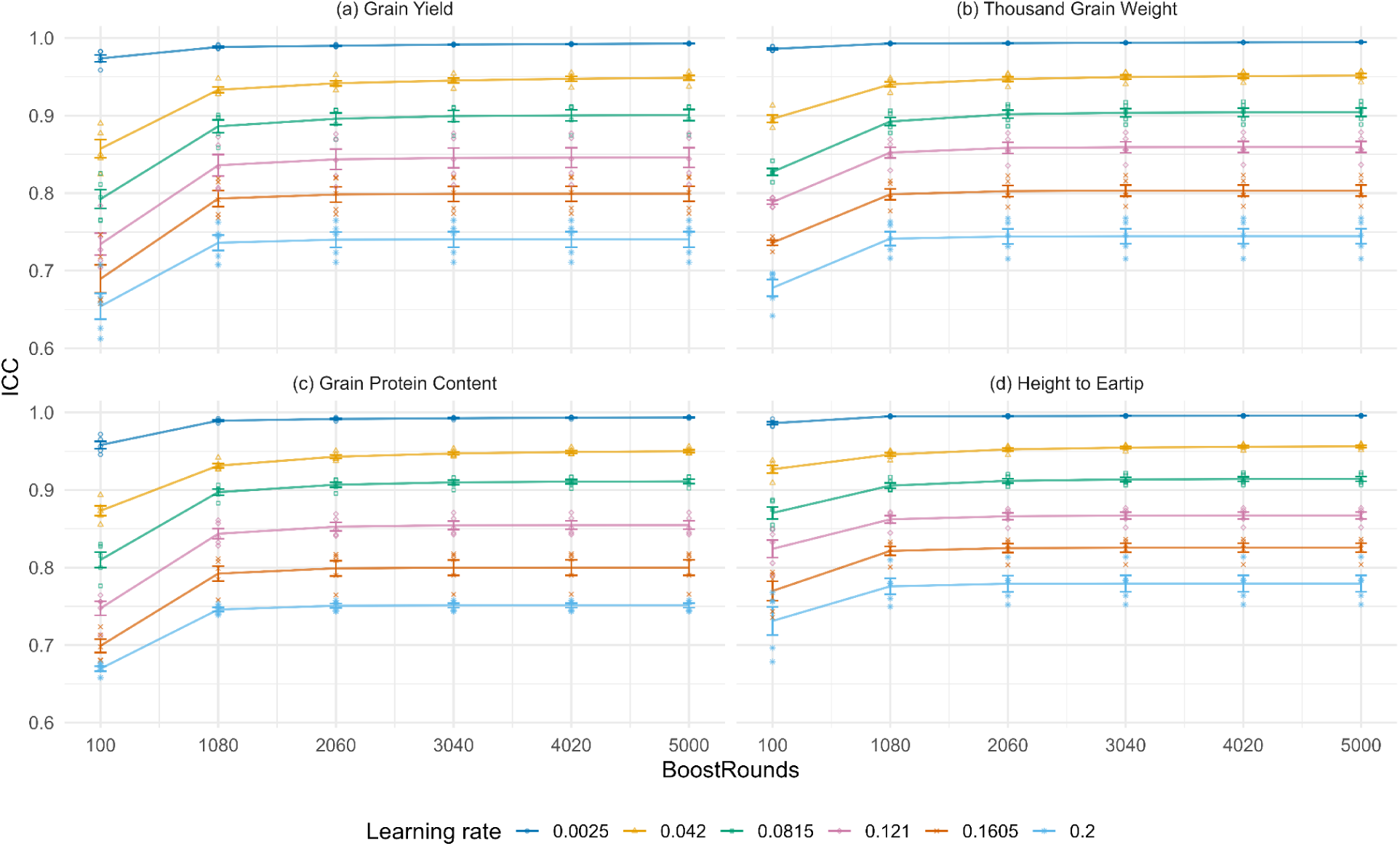
The plot displays the intra class correlation coefficient (ICC) compared to combinations of boost rounds and learning rate across four traits Grain Yield (GY - top left), Thousand Grain Weight (TGW - top right), Grain Protein Content (GPC - bottom left), Height to Ear Tip (HET - bottom right). The x-axis represents boost rounds and the y-axis shows ICC. For each trait and learning rate, mean ICC values are connected by dashed lines with error bars indicating ±1 standard error. Individual ICC observations appear as points. Each color corresponds to a distinct Learning Rate; Dark Blue (0.0025), Orange (0.042), Green (0.0815), Purple (0.121), Red-Orange (0.1605), and Light Blue (0.2).

**Figure 8.**
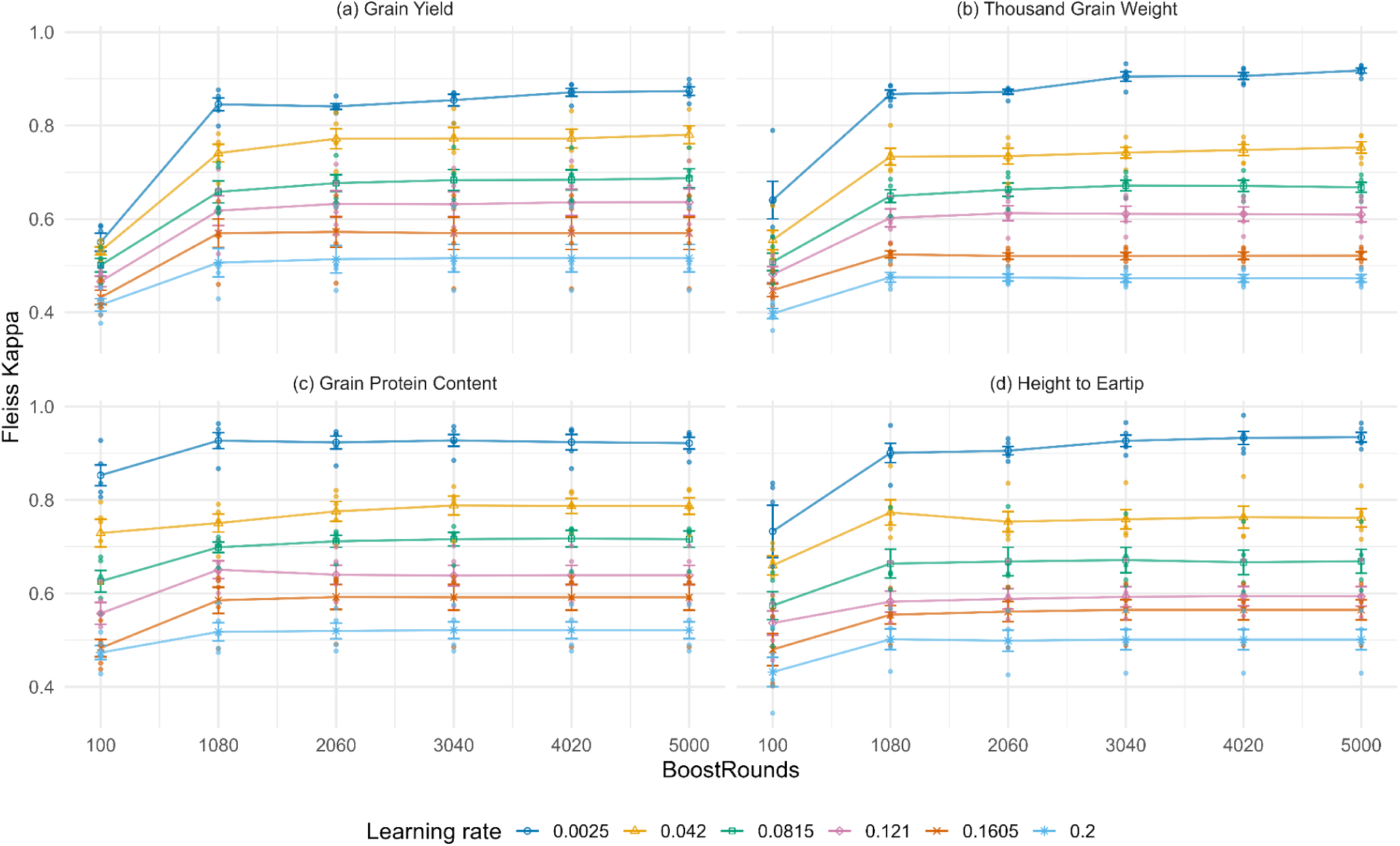
The plot displays the Fleiss’ κ compared to combinations of boost rounds and learning rate across four traits Grain Yield (GY - top left), Thousand Grain Weight (TGW - top right), Grain Protein Content (GPC - bottom left), Height to Ear Tip (HET - bottom right). The x-axis represents boost rounds and the y-axis shows Fleiss’ κ. For each trait and learning rate, mean Fleiss’ κ values are connected by dashed lines with error bars indicating ±1 standard error. Individual Fleiss’ κ observations appear as points. Each color corresponds to a distinct Learning Rate; Dark Blue (0.0025), Orange (0.042), Green (0.0815), Purple (0.121), Red-Orange (0.1605), and Light Blue (0.2).

Model generalizability was assessed by comparing the mean Pearson’s r on the training and test datasets for each learning rate across boosting rounds (Figure 9a-f). A performance gap was observed for all hyperparameter combinations, with training set correlations consistently higher than test set correlations. For learning rates of 0.042 and higher, training performance rapidly approached a correlation of 1.0, creating a large gap relative to the test performance (Fig. 9b-f). In contrast, for the lowest learning rate (v = 0.0025), training performance increased more gradually, resulting in the smallest performance gap between the training and test datasets among all tested learning rates (Fig. 9a).

**Figure 9.**
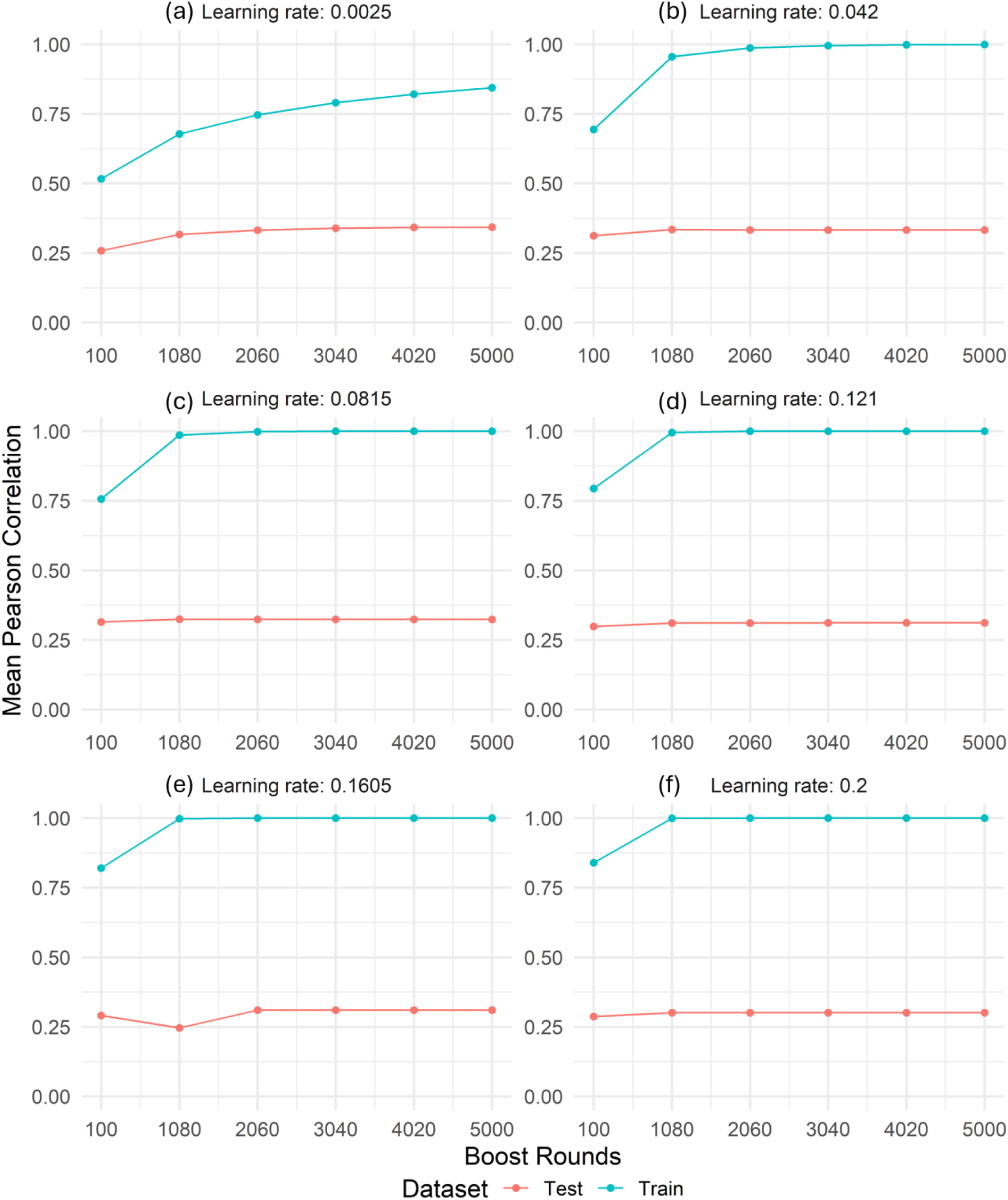
Shows the relationship between correlation coefficient and boosting rounds across different learning rates with traits aggregated. The y-axis shows the mean Pearson’s r coefficient. The x-axis shows the number of boosting rounds. Each facet represents a different learning rate. In each plot, the blue line depicts model performance on the training data, while the orange-red line shows performance on unseen test data, allowing the visualization of overfitting.

## Discussion

The stochastic characteristic of many modern genomic prediction models produces stochastic outputs; evaluating them solely on predictive accuracy ignores this critical instability and undermines their reliability for supporting selection and ranking decisions in breeding programs. Our multi-objective hyperparameter search tuning for model stability (via ICC and Fleiss’ κ) is a highly effective strategy to obtain reproducible predictions and is associated with smaller train - test gaps. A GBM configuration with the lowest learning rate (v = 0.0025) and > 2,000 boosting rounds maximized stability (Fig. 3a-b,7,8) and provided moderate but significant concurrent improvements in Pearson’s r and ranking (NDCG) performance (Fig. 3c, 3e); pairwise contrasts indicate that very low learning rates (0.0025 - 0.042) perform comparably for r and NDCG at higher rounds (Fig. 4,6, File S3).

While model output stability (ICC, Fleiss’ κ) was consistent across traits, performance metrics (Pearson’s r, AUC and NDCG) showed trait dependent responses, as indicated by significant trait × learning rate × boosting rounds interactions for Pearson’s r, AUC and NDCG in the aggregated analysis. This complexity supports the use of a multi-objective approach, as no single optimization strategy can meet all the diverse objectives within breeding programs.

Low learning rates increased stability to excellent levels (e.g, mean ICC ≈ 0.994 at 5,000 boosting rounds across traits), supporting consistent predictions (Fig. 3a, 7). Low learning rates also increased selection stability with Fleiss’ κ frequently exceeding 0.8 (Fig. 3b, 8). To interpret these coefficients, we adopt a unified framework. Following Shrout, who notes conceptual links between chance-corrected agreement statistics such as κ and reliability coefficients like the ICC, we apply a single set of modern benchmarks to both measures (Shrout 1998). Accordingly, we use the thresholds recommended by Koo and Li, defining reliability as good for values between0.75 and 0.90 and excellent for values greater than 0.90 (Koo and Li 2016). Under this framework, both results indicate good to excellent stability, implying highly reliable model predictions across continuous and categorical outputs.

Aggregated analysis identified a practical hyperparameter window: the lowest learning rate (v = 0.0025) maximized stability, while low learning rates up to v = 0.042, when combined with more than 2,000 boost rounds delivered comparable Pearson’s r and NDCG performance (Fig. 3, File S3 - Pairwise_overallmodel_Adj_NDCG and Pairwise_overallmodel_Adj_Pears). This suggests moderately boosted models with low learning rates can achieve similar performance for Pearsons’s r and NDCG with substantially less computation (Friedman 2002) (Fig. S3). Within this near optimal window, the learning rate of 0.0025 with 5,000 boosting rounds achieved the highest combined stability, predictive accuracy, and ranking efficiency (Friedman 2002).

In contrast to other metrics, classification accuracy (AUC) revealed a conflict in optimization objectives: it not only performed worse than random chance across all models and traits but also favored a different hyperparameter configuration, where fewer boosting rounds and higher learning rates yielded better relative discriminatory ability. Since the total AUC metric evaluates performance across all decision thresholds, it can obscure a model’s utility for breeders by giving undue weight to irrelevant portions of the ROC curve (Bradley 1997; Peng et al. 2024). We therefore support that a partial AUC (pAUC), by restricting the analysis to a relevant window of high True Positive and low-to-moderate False Positive Rates (TPR, FPR), should be explored for a more relevant metric for gauging the discriminatory power of models for candidate selection (McClish 1989; Peng et al. 2024). A pAUC can adjust the FPR tolerance to match the decision-making context. For example, in pre-breeding where a common aim is to maximize diversity, breeders may tolerate higher FPR (e.g, > 0.2) to avoid prematurely discarding potentially elite candidates, whereas late stage multi-environment performance trials may have stricter FPR (e.g, FPR < 0.1) due to the cost per entry (Simmonds 1993). Our findings suggest that model optimization requires a strategic choice based on the primary objective, allowable tradeoffs and trait of interest, as a single set of hyperparameters could not deliver optimal performance across all evaluation metrics. The poor absolute values for AUC are likely a consequence of the challenging cross-environment scenario where candidate assignment was based on the top 25% of observed phenotypes within the test environment, creating a ‘moving target’ and, due to this fixed upper threshold, resulting in class imbalance. The unmodeled environmental covariates were likely an important factor in the poor cross-environment prediction. The absence of these covariates inhibited the modeling of G × E interactions, which can degrade the cross-season performance (Jarquín 2017; Tomar et al. 2021; Montesinos-López et al. 2022).

Despite the model’s modest prediction and classification accuracy, its consistently high ranking performance (NDCG@20% > 0.85) indicates practical value for elite-line discovery, where correctly ranking top candidates is more valuable than precise estimation of their breeding values. This high ranking performance, in contrast to the lower Pearson’s r values, is explained by the nature of the NDCG metric, which focuses specifically on rewarding the correct ordering of top-performing entries rather than penalizing errors across the entire distribution (Blondel et al. 2015; Pan et al. 2024). The suitability of this metric for breeding decisions, where correctly ranking and selecting elite lines is a primary goal, has led to its increasing adoption (Gianola et al. 2018; Negussie et al. 2022; Li et al. 2023; Pan et al. 2024).

The ANOVA results confirm that model performance is influenced mostly by main effects of learning rate and boosting rounds but also by their significant interaction, which supports the need for co-optimization. This hierarchy of effects is consistent with previous research where learning rate was a dominant hyperparameter for XGBoost model with a general support for lower learning rates (Westhues et al. 2021). The learning rate provides regularization that prevents overfitting to subsample artifacts by shrinking the contribution of each new boost round (Elith et al. 2008). The non-significant three-way interaction for prediction stability (ICC) is an important result as it indicates that the relationship between learning rate and boosting rounds was consistent across all traits, suggesting that an optimized hyperparameter combination for stability may be broadly applicable across traits.

An important finding was the trait-specific response to hyperparameter tuning. While the overall pattern of the hyperparameters explored (favoring low learning rates and high boost rounds) was consistent, Grain Protein Content (GPC) displayed the clearest and most substantial gain in model accuracy (Pearson’s r) when using the lowest learning rate (v = 0.0025), when combined with sufficient boosting rounds (ntrees > 1000). The findings for GPC show that small incremental error corrections in each boosting round may better capture the cumulative contribution of many small-effect loci, which are more effectively modelled through slower learning rates and high boosting rounds (Azodi et al. 2019; Sandhu et al. 2021; Paina and Gregersen 2023).

While our results demonstrate the benefits of GBM hyperparameter tuning for genomic prediction, we acknowledge the limitations of this study. First, the performance metrics, Pearson’s r and AUC, remained low-moderate in absolute terms across all quantitative traits. This poor performance is supported by a clear train-test gap, suggesting overfitting (Fig. 9). Lower learning rates were associated with smaller train-test gaps (Fig. 9). By shrinking each boost round’s contribution, lower learning rates reduce the influence of subsample artifacts introduced by stochastic subsampling and reduce noise amplification across iterations, improving generalization (Friedman 2001; Friedman 2002). This result aligns with established principles, confirming that learning rates are useful for maintaining generalizability and mitigating overfitting, especially when using high boosting rounds and datasets with large numbers of predictor variables (Friedman 2001; Elith et al. 2008).

Differences between training and test environments were not modeled via environmental covariates or multi-environment-trial (MET) terms. The inclusion of MET data presents an opportunity to improve model performance, as including environmental covariates has been shown to benefit generalizability by better capturing G × E interactions in MET studies (Jarquín 2017; Scott et al. 2021; Westhues et al. 2021; Montesinos-López et al. 2022).

The small n size and low haplotype diversity of the dataset (500 lines) are another area for future expansion. The Multi-parent Advanced Generation Inter Cross (MAGIC) wheat dataset (Scott et al. 2021) offered broad allelic diversity but still could be improved by increasing haplotype variation (van de Wouw et al. 2010; Fradgley et al. 2019; Scott et al. 2021). A larger population with higher haplotype diversity could further improve the reliability and performance of the evaluated models (Norman et al. 2018).

Another direction for exploration is to expand our hyperparameter grid to tune other important GBM hyperparameters including tree depth (interaction.depth) and regularization parameters like minimum node size and colsample rate (Westhues et al. 2021). Additionally, tuning stochastic subsampling values in the range 0.4 - 0.6 have been shown to improve performance, as suggested in the foundational work (Friedman 2002). To determine global optima, we suggest using Bayesian optimization for its adaptability and efficiency, or expanding a manual grid search to cover broader ranges and additional types of hyperparameters for a more thorough model optimization (Shahriari et al. 2016).

Future studies could address the stability of feature contributions or the performance trade-offs when optimizing model structural stability. One study identified a trade-off showing a ∼2% reduction in predictive power for a 30% gain in structural stability (Bertsimas et al. 2025). Their work examined structural stability during model retraining, while our study focused on output stability across iterations. Applying a mixed integer optimization approach, which constrains variation in model structure, such as split variables and leaf values across successive retraining, could show whether the loci driving predictions across separate model training remain consistent, support feature selection efforts and provide evidence of predictive performance tradeoffs when structural stability is improved (Bertsimas et al. 2025).

Computation time increases consistently with higher boosting rounds, and adopting efficient implementations can reduce costs, which is particularly useful in time-sensitive applications (Fig. S3). An effective model strategy to optimize computational costs is to utilize early stopping, allowing the model to find parameters that generalize well and mitigate excessive epochs that can degrade generalizability (Prechelt 2012). The gbm.fit function lacks early stopping; however, alternative gradient boosting packages, XGBoost and LightGBM, contain an early_stopping_rounds argument that can be directly used to set the number of boosting iterations the model can perform without improvement on a specified evaluation metric, after which the training automatically stops (Chen et al. 2014; Ke et al. 2017; lightgbm package - RDocumentation).

This study focussed on one modeling approach (GBM). In future work, benchmarking different algorithms (GBM vs. random forests vs. GBLUP, etc.) using the same multi-objective evaluation under the same cross-year prediction scenario would be useful for model selection, with each algorithm being individually optimized.

Our study shows that model stability can be greatly improved without sacrificing ranking performance and correlation, although classification (AUC) favored different hyperparameters. These findings support the use of multi-objective optimization in genomic prediction models, where stability is evaluated alongside predictive performance metrics. This work outlines a practical framework for tuning stochastic models to deliver reliable and consistent predictions suitable for breeding decisions.

## Data Availability

File S1 contains, Packages and Functions, Formulae of metrics and models, Hyperparameter Grid, Computing Environment.

File S2 contains, evaluation metric descriptive statistics used in data visualization.

File S3 contains, pairwise comparisons of the estimated marginal means and ANOVA tables

File S4 contains the complete evaluation metric results for each model configuration (stability metric per subset, performance metrics per iteration and subset)

File S5 contains the summary statistics for each trait-metric combination (aggregated across configurations)

The R scripts and datasets used for modelling, statistical analysis, and figure generation are available in the GitHub repository (https://github.com/HenryOmix/Multi-objective-Evaluation-and-Optimization-of-Stochastic-Gradient-Boosting-Machine).

The specific version of the code used to produce the results in this manuscript is permanently archived on Zenodo:

HenryOmix. (2025).

HenryOmix/Multi-objective-Evaluation-and-Optimization-of-Stochastic-Gradient-Boosting-Machine: v1.03 (v1.03). Zenodo. https://doi.org/10.5281/zenodo.16785772

## Acknowledgements

We are grateful to Professor Dr. Wolfgang Link for his valuable input and for the discussions within the Department of Crop Sciences, Plant Breeding Methodology Division (University of Göttingen) for fostering a supportive research environment. We thank the University of Göttingen for providing computing resources and IT support used for genotype imputation in 2024. We acknowledge that OpenAI’s ChatGPT assisted in text grammar and code refactoring, which were reviewed and verified by the authors. We especially thank the authors from the Scott et al. 2021 study for their genomic and phenotypic data.

