## Supplementary figures and images for "Multi-objective Evaluation and Optimization of Stochastic Gradient Boosting Machines for Genomic Prediction and Selection in Wheat (*Triticum aestivum*) Breeding"

### Supplementary figure 1_Fig.S1

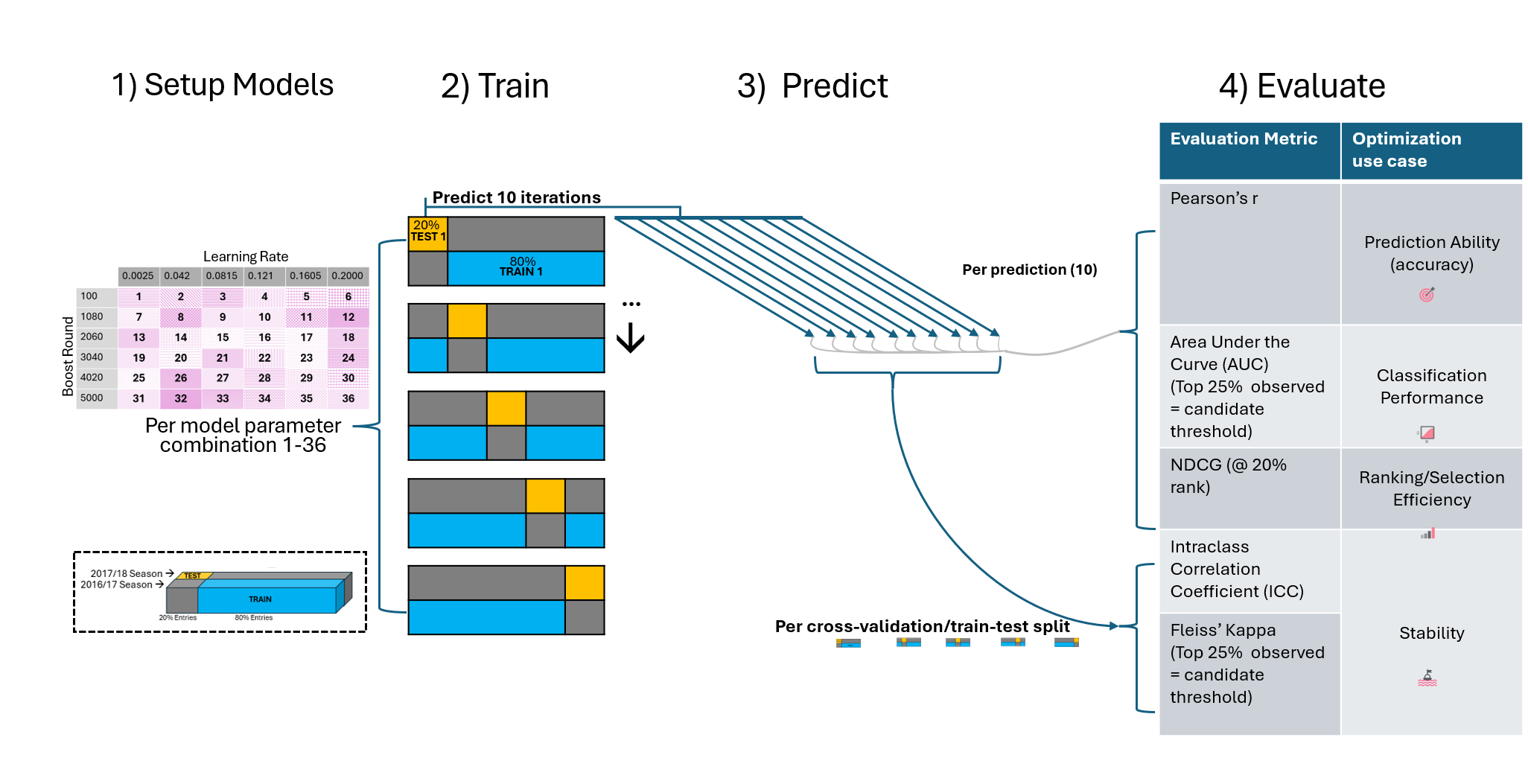

### Supplementary figure 2_Fig.S2

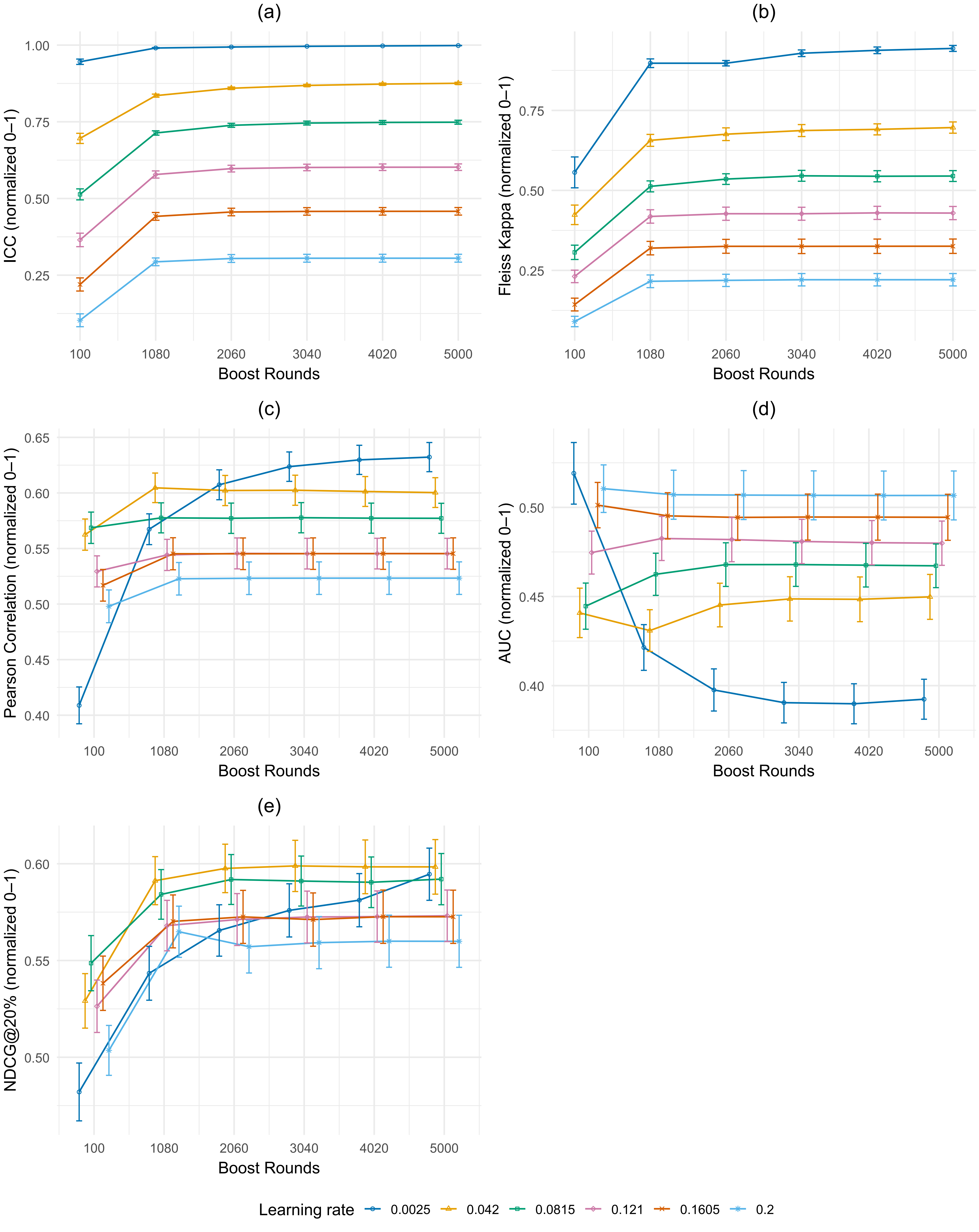

### Supplementary figure 3_Fig.S3

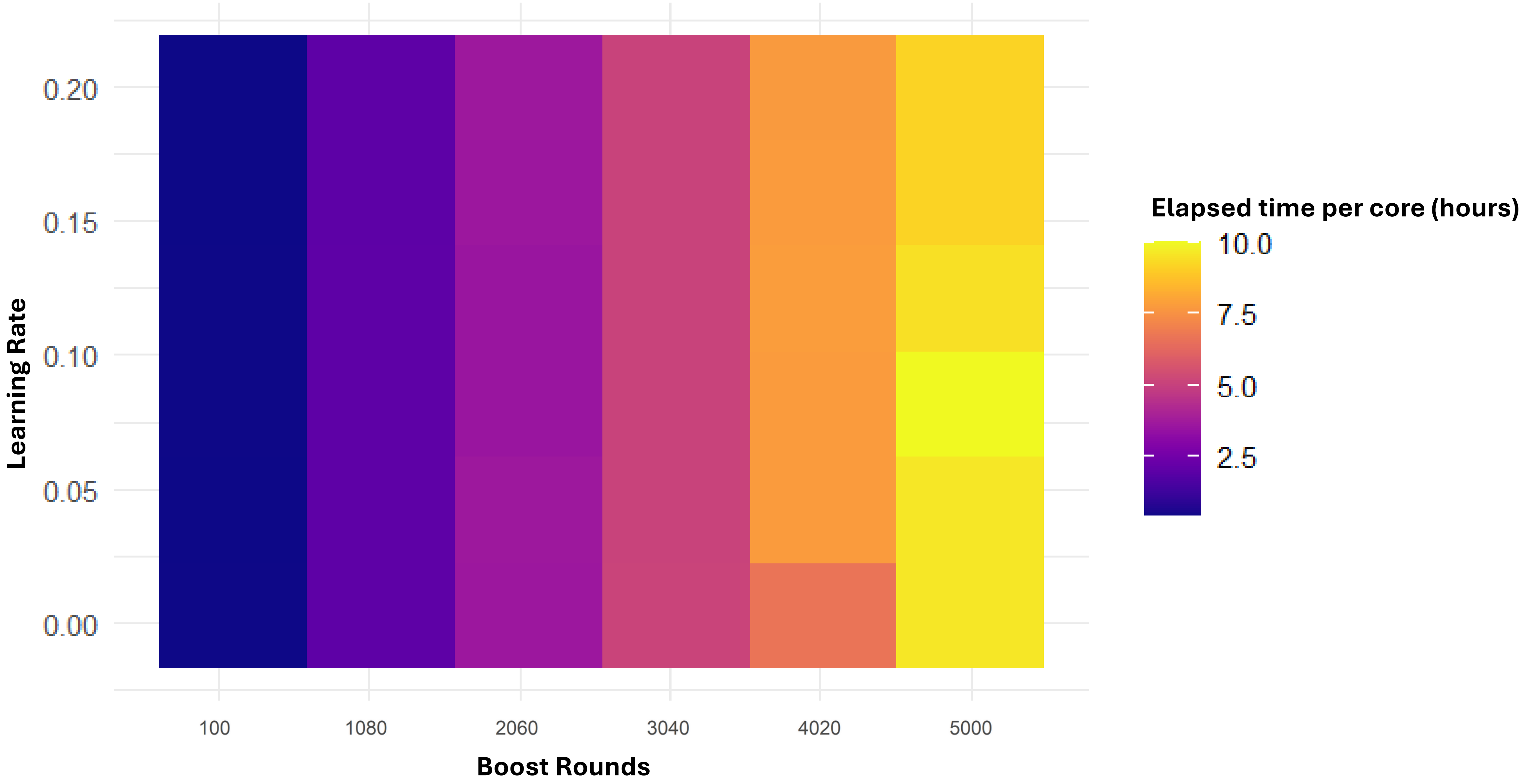

### Supplementary figure 4_Fig.S4

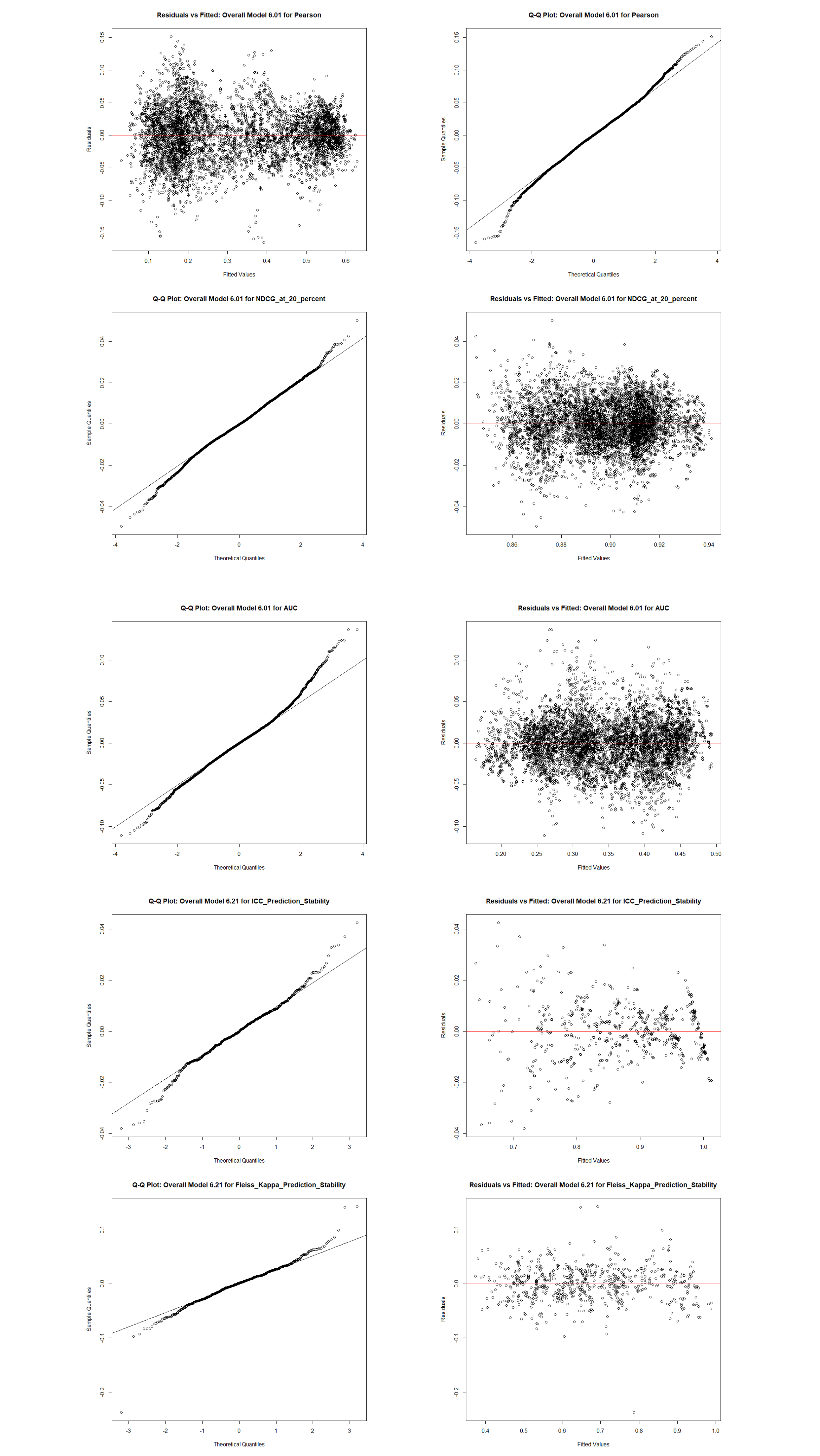

### Supplementary figure 5_Fig.S5

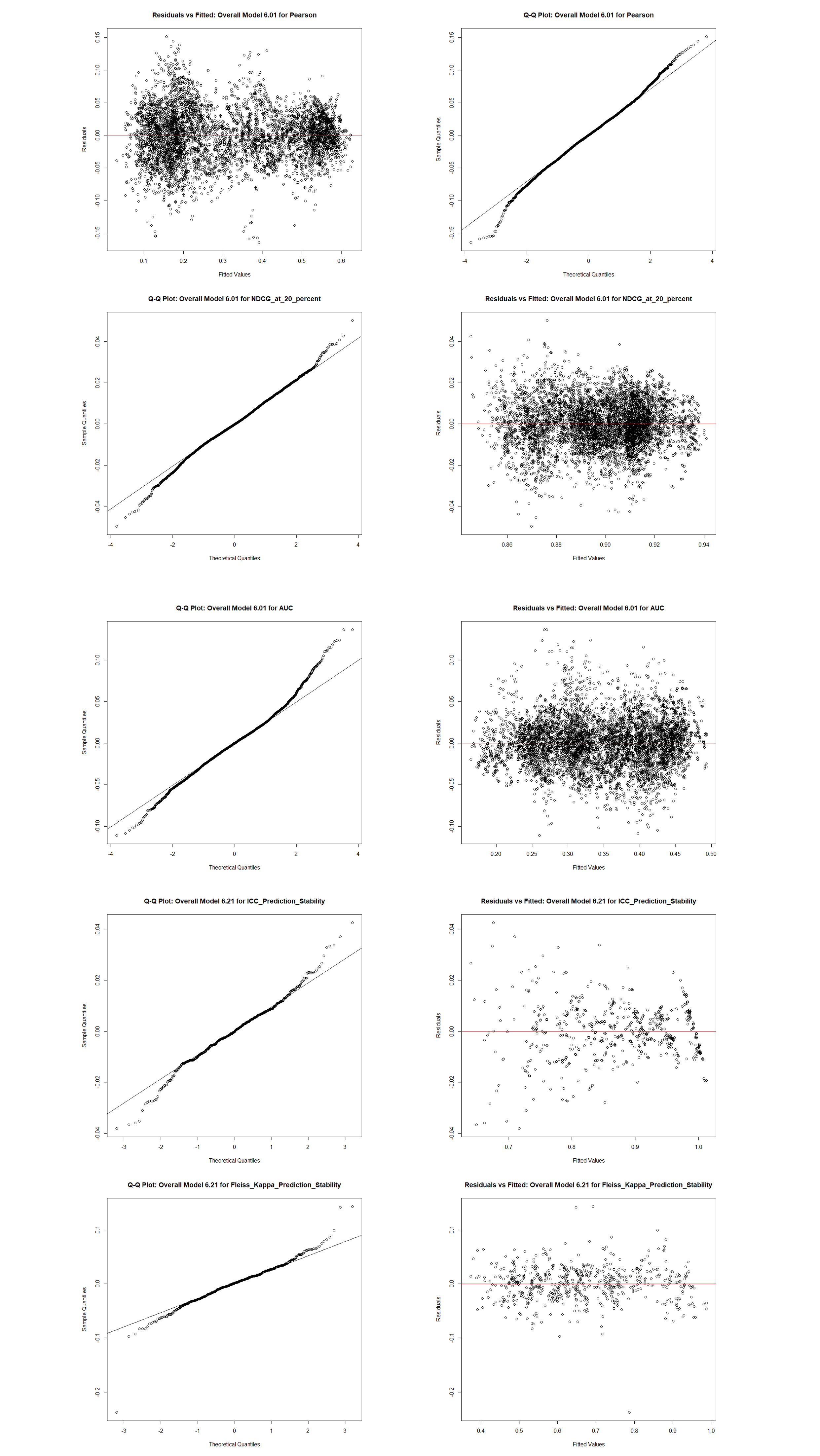
