## Supplementary material for "Multi-objective Evaluation and Optimization of Stochastic Gradient Boosting Machines for Genomic Prediction and Selection in Wheat (*Triticum aestivum*) Breeding": File S2

| | | | | GPC | | | | | | | | GY | | | | | | | | HET | | | | | | | | TGW | | | | | | | |
|---|---|---|---|---|---|---|---|---|---|---|---|---|---|---|---|---|---|---|---|---|---|---|---|---|---|---|---|---|---|---|---|---|---|---|---|
| Metric | Boost Rounds | Learning Rate | N | Mean | SD | SE | Min | Max | IQR | upper outliers | lower outliers | Mean | SD | SE | Min | Max | IQR | upper outliers | lower outliers | Mean | SD | SE | Min | Max | IQR | upper outliers | lower outliers | Mean | SD | SE | Min | Max | IQR | upper outliers | lower outliers |
| AUC | 100 | 0.0025 | 50 | 0.3548 | 0.0524 | 0.0074 | 0.2411 | 0.4551 | 0.0788 | 0 | 0 | 0.4286 | 0.0583 | 0.0082 | 0.3481 | 0.5335 | 0.0973 | 0 | 0 | 0.3348 | 0.0348 | 0.0049 | 0.2846 | 0.4068 | 0.0526 | 0 | 0 | 0.315 | 0.081 | 0.0115 | 0.1751 | 0.437 | 0.1323 | 0 | 0 |
| AUC | 100 | 0.042 | 50 | 0.3533 | 0.0536 | 0.0076 | 0.2519 | 0.4486 | 0.0904 | 0 | 0 | 0.4108 | 0.0517 | 0.0073 | 0.3178 | 0.4924 | 0.0753 | 0 | 0 | 0.2746 | 0.0351 | 0.005 | 0.1816 | 0.3314 | 0.0488 | 0 | 0 | 0.314 | 0.0699 | 0.0099 | 0.1924 | 0.4362 | 0.1151 | 0 | 0 |
| AUC | 100 | 0.0815 | 50 | 0.3533 | 0.043 | 0.0061 | 0.2762 | 0.44 | 0.0614 | 0 | 0 | 0.4166 | 0.0478 | 0.0068 | 0.3303 | 0.5097 | 0.0745 | 0 | 0 | 0.2703 | 0.0467 | 0.0066 | 0.1978 | 0.3692 | 0.0736 | 0 | 0 | 0.316 | 0.0593 | 0.0084 | 0.2 | 0.4389 | 0.0899 | 0 | 0 |
| AUC | 100 | 0.121 | 50 | 0.3673 | 0.0446 | 0.0063 | 0.2811 | 0.46 | 0.0692 | 0 | 0 | 0.4178 | 0.0417 | 0.0059 | 0.3303 | 0.5368 | 0.06 | 1 | 0 | 0.2777 | 0.044 | 0.0062 | 0.1832 | 0.3741 | 0.0551 | 0 | 0 | 0.3282 | 0.0561 | 0.0079 | 0.2081 | 0.4919 | 0.0761 | 1 | 0 |
| AUC | 100 | 0.1605 | 50 | 0.3845 | 0.0543 | 0.0077 | 0.273 | 0.4968 | 0.0782 | 0 | 0 | 0.4217 | 0.0473 | 0.0067 | 0.3081 | 0.5124 | 0.0511 | 0 | 1 | 0.2847 | 0.0463 | 0.0065 | 0.1584 | 0.3724 | 0.0566 | 0 | 1 | 0.3294 | 0.0476 | 0.0067 | 0.2265 | 0.4438 | 0.0604 | 0 | 0 |
| AUC | 100 | 0.2 | 50 | 0.3874 | 0.0446 | 0.0063 | 0.3059 | 0.5038 | 0.0653 | 0 | 0 | 0.4198 | 0.0521 | 0.0074 | 0.3335 | 0.533 | 0.0792 | 0 | 0 | 0.2843 | 0.0536 | 0.0076 | 0.1492 | 0.3903 | 0.0768 | 0 | 0 | 0.3411 | 0.0527 | 0.0075 | 0.2454 | 0.427 | 0.0945 | 0 | 0 |
| AUC | 1080 | 0.0025 | 50 | 0.3457 | 0.0469 | 0.0066 | 0.2741 | 0.4184 | 0.0909 | 0 | 0 | 0.4062 | 0.0438 | 0.0062 | 0.34 | 0.4903 | 0.0738 | 0 | 0 | 0.2736 | 0.0257 | 0.0036 | 0.2378 | 0.3303 | 0.0284 | 2 | 0 | 0.3053 | 0.0732 | 0.0103 | 0.2038 | 0.4086 | 0.1323 | 0 | 0 |
| AUC | 1080 | 0.042 | 50 | 0.3769 | 0.0396 | 0.0056 | 0.3173 | 0.4508 | 0.0715 | 0 | 0 | 0.4184 | 0.0296 | 0.0042 | 0.3519 | 0.4908 | 0.038 | 0 | 0 | 0.2454 | 0.0544 | 0.0077 | 0.1449 | 0.3454 | 0.077 | 0 | 0 | 0.3001 | 0.0512 | 0.0072 | 0.2178 | 0.4227 | 0.0736 | 0 | 0 |
| AUC | 1080 | 0.0815 | 50 | 0.3881 | 0.0495 | 0.007 | 0.3092 | 0.4751 | 0.087 | 0 | 0 | 0.4278 | 0.0313 | 0.0044 | 0.3692 | 0.5032 | 0.0458 | 0 | 0 | 0.2534 | 0.0503 | 0.0071 | 0.1443 | 0.3416 | 0.07 | 0 | 0 | 0.3053 | 0.0438 | 0.0062 | 0.2065 | 0.427 | 0.043 | 2 | 1 |
| AUC | 1080 | 0.121 | 50 | 0.3976 | 0.0579 | 0.0082 | 0.3103 | 0.5027 | 0.1018 | 0 | 0 | 0.4243 | 0.0369 | 0.0052 | 0.3438 | 0.5389 | 0.0415 | 1 | 0 | 0.2634 | 0.0479 | 0.0068 | 0.1643 | 0.3573 | 0.0665 | 0 | 0 | 0.3132 | 0.043 | 0.0061 | 0.2497 | 0.4314 | 0.0449 | 2 | 0 |
| AUC | 1080 | 0.1605 | 50 | 0.4058 | 0.0624 | 0.0088 | 0.2886 | 0.5119 | 0.1107 | 0 | 0 | 0.4285 | 0.0356 | 0.005 | 0.3492 | 0.4914 | 0.0534 | 0 | 0 | 0.2656 | 0.0511 | 0.0072 | 0.1584 | 0.3422 | 0.0739 | 0 | 0 | 0.3118 | 0.0418 | 0.0059 | 0.2232 | 0.4184 | 0.045 | 4 | 0 |
| AUC | 1080 | 0.2 | 50 | 0.4005 | 0.0608 | 0.0086 | 0.2795 | 0.5292 | 0.0935 | 0 | 0 | 0.4257 | 0.0445 | 0.0063 | 0.3324 | 0.5341 | 0.0612 | 0 | 0 | 0.2792 | 0.056 | 0.0079 | 0.167 | 0.4092 | 0.0803 | 0 | 0 | 0.3202 | 0.0481 | 0.0068 | 0.2395 | 0.4341 | 0.0492 | 4 | 0 |
| AUC | 2060 | 0.0025 | 50 | 0.3426 | 0.0462 | 0.0065 | 0.2659 | 0.3989 | 0.0877 | 0 | 0 | 0.4034 | 0.0345 | 0.0049 | 0.3524 | 0.4514 | 0.0657 | 0 | 0 | 0.2573 | 0.035 | 0.005 | 0.2135 | 0.3324 | 0.037 | 2 | 0 | 0.3018 | 0.067 | 0.0095 | 0.2114 | 0.3989 | 0.1235 | 0 | 0 |
| AUC | 2060 | 0.042 | 50 | 0.3865 | 0.0511 | 0.0072 | 0.3189 | 0.4886 | 0.098 | 0 | 0 | 0.4248 | 0.0288 | 0.0041 | 0.3638 | 0.4908 | 0.0434 | 0 | 0 | 0.2469 | 0.0518 | 0.0073 | 0.1492 | 0.3259 | 0.0774 | 0 | 0 | 0.2971 | 0.0475 | 0.0067 | 0.2373 | 0.4222 | 0.0516 | 2 | 0 |
| AUC | 2060 | 0.0815 | 50 | 0.3925 | 0.0573 | 0.0081 | 0.2935 | 0.4946 | 0.098 | 0 | 0 | 0.4292 | 0.0322 | 0.0046 | 0.3686 | 0.5043 | 0.0445 | 0 | 0 | 0.2557 | 0.0479 | 0.0068 | 0.1503 | 0.3389 | 0.0641 | 0 | 0 | 0.3027 | 0.0416 | 0.0059 | 0.2103 | 0.4162 | 0.0408 | 2 | 1 |
| AUC | 2060 | 0.121 | 50 | 0.3992 | 0.0593 | 0.0084 | 0.3076 | 0.5135 | 0.1011 | 0 | 0 | 0.424 | 0.0358 | 0.0051 | 0.3557 | 0.5324 | 0.0439 | 1 | 0 | 0.264 | 0.0487 | 0.0069 | 0.1627 | 0.3589 | 0.0624 | 0 | 0 | 0.3103 | 0.0409 | 0.0058 | 0.2481 | 0.4297 | 0.0435 | 2 | 0 |
| AUC | 2060 | 0.1605 | 50 | 0.4059 | 0.0624 | 0.0088 | 0.2989 | 0.5114 | 0.1153 | 0 | 0 | 0.4267 | 0.0349 | 0.0049 | 0.3508 | 0.4957 | 0.0501 | 0 | 0 | 0.267 | 0.0516 | 0.0073 | 0.1573 | 0.3443 | 0.0776 | 0 | 0 | 0.3114 | 0.0405 | 0.0057 | 0.2254 | 0.4184 | 0.0453 | 3 | 0 |
| AUC | 2060 | 0.2 | 50 | 0.4012 | 0.0606 | 0.0086 | 0.2805 | 0.5297 | 0.0931 | 0 | 0 | 0.4252 | 0.0442 | 0.0063 | 0.3335 | 0.5249 | 0.0586 | 0 | 0 | 0.2791 | 0.0556 | 0.0079 | 0.1638 | 0.4038 | 0.0778 | 0 | 0 | 0.32 | 0.0487 | 0.0069 | 0.24 | 0.4416 | 0.0476 | 4 | 0 |
| AUC | 3040 | 0.0025 | 50 | 0.3422 | 0.0449 | 0.0064 | 0.2622 | 0.4027 | 0.0797 | 0 | 0 | 0.4034 | 0.029 | 0.0041 | 0.3638 | 0.4524 | 0.0566 | 0 | 0 | 0.253 | 0.0408 | 0.0058 | 0.2 | 0.333 | 0.0547 | 0 | 0 | 0.2984 | 0.0622 | 0.0088 | 0.2141 | 0.3881 | 0.1203 | 0 | 0 |
| AUC | 3040 | 0.042 | 50 | 0.3894 | 0.0562 | 0.0079 | 0.3086 | 0.5038 | 0.1035 | 0 | 0 | 0.4258 | 0.0269 | 0.0038 | 0.3616 | 0.4849 | 0.0373 | 0 | 0 | 0.2492 | 0.0514 | 0.0073 | 0.1535 | 0.3243 | 0.0784 | 0 | 0 | 0.2942 | 0.045 | 0.0064 | 0.2422 | 0.4049 | 0.0434 | 6 | 0 |
| AUC | 3040 | 0.0815 | 50 | 0.3935 | 0.0576 | 0.0082 | 0.293 | 0.5016 | 0.0954 | 0 | 0 | 0.4289 | 0.0325 | 0.0046 | 0.3676 | 0.5022 | 0.0422 | 0 | 0 | 0.2559 | 0.0475 | 0.0067 | 0.1524 | 0.3341 | 0.0628 | 0 | 0 | 0.3019 | 0.0405 | 0.0057 | 0.2092 | 0.4141 | 0.0404 | 1 | 1 |
| AUC | 3040 | 0.121 | 50 | 0.3992 | 0.0589 | 0.0083 | 0.307 | 0.5173 | 0.102 | 0 | 0 | 0.423 | 0.0361 | 0.0051 | 0.3578 | 0.5314 | 0.0447 | 1 | 0 | 0.2648 | 0.0492 | 0.007 | 0.1616 | 0.3654 | 0.0615 | 0 | 0 | 0.3094 | 0.0403 | 0.0057 | 0.2492 | 0.4314 | 0.0411 | 2 | 0 |
| AUC | 3040 | 0.1605 | 50 | 0.4063 | 0.0623 | 0.0088 | 0.3016 | 0.5097 | 0.1153 | 0 | 0 | 0.4266 | 0.0349 | 0.0049 | 0.3503 | 0.4935 | 0.0503 | 0 | 0 | 0.2673 | 0.0518 | 0.0073 | 0.1573 | 0.3432 | 0.0809 | 0 | 0 | 0.3112 | 0.0403 | 0.0057 | 0.2259 | 0.4184 | 0.0453 | 3 | 0 |
| AUC | 3040 | 0.2 | 50 | 0.4013 | 0.0606 | 0.0086 | 0.2789 | 0.5303 | 0.0916 | 0 | 0 | 0.425 | 0.0441 | 0.0062 | 0.333 | 0.5243 | 0.0584 | 0 | 0 | 0.2792 | 0.0556 | 0.0079 | 0.1632 | 0.4043 | 0.0782 | 0 | 0 | 0.32 | 0.0488 | 0.0069 | 0.2405 | 0.4416 | 0.0473 | 4 | 0 |
| AUC | 4020 | 0.0025 | 50 | 0.3423 | 0.042 | 0.0059 | 0.2789 | 0.3989 | 0.0732 | 0 | 0 | 0.4044 | 0.0277 | 0.0039 | 0.3649 | 0.4568 | 0.0541 | 0 | 0 | 0.2515 | 0.0457 | 0.0065 | 0.1854 | 0.3314 | 0.0622 | 0 | 0 | 0.2979 | 0.0588 | 0.0083 | 0.2184 | 0.3908 | 0.1131 | 0 | 0 |
| AUC | 4020 | 0.042 | 50 | 0.3906 | 0.0578 | 0.0082 | 0.3022 | 0.5016 | 0.105 | 0 | 0 | 0.4258 | 0.0263 | 0.0037 | 0.3659 | 0.4865 | 0.0373 | 0 | 0 | 0.2494 | 0.0515 | 0.0073 | 0.1546 | 0.3222 | 0.0788 | 0 | 0 | 0.2924 | 0.0443 | 0.0063 | 0.2335 | 0.4059 | 0.0462 | 3 | 0 |
| AUC | 4020 | 0.0815 | 50 | 0.3941 | 0.0576 | 0.0081 | 0.2951 | 0.5038 | 0.0973 | 0 | 0 | 0.428 | 0.0325 | 0.0046 | 0.367 | 0.5016 | 0.0424 | 0 | 0 | 0.2562 | 0.0479 | 0.0068 | 0.1497 | 0.3335 | 0.0664 | 0 | 0 | 0.3016 | 0.0399 | 0.0056 | 0.2097 | 0.4151 | 0.0426 | 1 | 1 |
| AUC | 4020 | 0.121 | 50 | 0.3992 | 0.0588 | 0.0083 | 0.3076 | 0.5184 | 0.1003 | 0 | 0 | 0.4227 | 0.0361 | 0.0051 | 0.3578 | 0.5308 | 0.0442 | 1 | 0 | 0.2647 | 0.0491 | 0.0069 | 0.1622 | 0.3638 | 0.0609 | 0 | 0 | 0.3091 | 0.04 | 0.0057 | 0.2486 | 0.4314 | 0.0409 | 2 | 0 |
| AUC | 4020 | 0.1605 | 50 | 0.4063 | 0.0624 | 0.0088 | 0.3016 | 0.5103 | 0.1157 | 0 | 0 | 0.4266 | 0.0349 | 0.0049 | 0.3503 | 0.4941 | 0.05 | 0 | 0 | 0.2673 | 0.0519 | 0.0073 | 0.1573 | 0.3427 | 0.0819 | 0 | 0 | 0.311 | 0.0401 | 0.0057 | 0.2259 | 0.4178 | 0.0447 | 3 | 0 |
| AUC | 4020 | 0.2 | 50 | 0.4013 | 0.0605 | 0.0086 | 0.2789 | 0.5303 | 0.0914 | 0 | 0 | 0.4249 | 0.0442 | 0.0062 | 0.333 | 0.5254 | 0.0588 | 0 | 0 | 0.2792 | 0.0556 | 0.0079 | 0.1632 | 0.4043 | 0.0784 | 0 | 0 | 0.32 | 0.0487 | 0.0069 | 0.2405 | 0.4411 | 0.0473 | 4 | 0 |
| AUC | 5000 | 0.0025 | 50 | 0.3453 | 0.0409 | 0.0058 | 0.2838 | 0.4011 | 0.0716 | 0 | 0 | 0.406 | 0.0271 | 0.0038 | 0.3681 | 0.4573 | 0.045 | 0 | 0 | 0.2486 | 0.0492 | 0.007 | 0.1789 | 0.3341 | 0.0619 | 0 | 0 | 0.2991 | 0.0565 | 0.008 | 0.2232 | 0.3843 | 0.108 | 0 | 0 |
| AUC | 5000 | 0.042 | 50 | 0.3911 | 0.0589 | 0.0083 | 0.3 | 0.5076 | 0.1039 | 0 | 0 | 0.4261 | 0.0269 | 0.0038 | 0.3659 | 0.4849 | 0.0397 | 0 | 0 | 0.2505 | 0.0513 | 0.0072 | 0.1595 | 0.3243 | 0.0803 | 0 | 0 | 0.2919 | 0.0431 | 0.0061 | 0.2335 | 0.4022 | 0.0424 | 4 | 0 |
| AUC | 5000 | 0.0815 | 50 | 0.3941 | 0.0576 | 0.0081 | 0.2951 | 0.5043 | 0.0972 | 0 | 0 | 0.4276 | 0.0329 | 0.0046 | 0.3649 | 0.5011 | 0.0441 | 0 | 0 | 0.2564 | 0.0479 | 0.0068 | 0.1503 | 0.3324 | 0.0676 | 0 | 0 | 0.3014 | 0.0398 | 0.0056 | 0.2108 | 0.4135 | 0.0424 | 1 | 1 |
| AUC | 5000 | 0.121 | 50 | 0.3992 | 0.0589 | 0.0083 | 0.307 | 0.5184 | 0.1001 | 0 | 0 | 0.4225 | 0.036 | 0.0051 | 0.3578 | 0.5308 | 0.0438 | 1 | 0 | 0.2647 | 0.0493 | 0.007 | 0.1622 | 0.3649 | 0.0614 | 0 | 0 | 0.309 | 0.04 | 0.0057 | 0.2486 | 0.4314 | 0.0408 | 2 | 0 |
| AUC | 5000 | 0.1605 | 50 | 0.4063 | 0.0624 | 0.0088 | 0.3022 | 0.5097 | 0.1157 | 0 | 0 | 0.4265 | 0.0349 | 0.0049 | 0.3503 | 0.4935 | 0.0496 | 0 | 0 | 0.2673 | 0.0519 | 0.0073 | 0.1568 | 0.3427 | 0.0819 | 0 | 0 | 0.311 | 0.0401 | 0.0057 | 0.2259 | 0.4178 | 0.0447 | 3 | 0 |
| AUC | 5000 | 0.2 | 50 | 0.4012 | 0.0605 | 0.0086 | 0.2789 | 0.5303 | 0.0914 | 0 | 0 | 0.4249 | 0.0442 | 0.0062 | 0.333 | 0.5254 | 0.0588 | 0 | 0 | 0.2793 | 0.0556 | 0.0079 | 0.1632 | 0.4043 | 0.0784 | 0 | 0 | 0.32 | 0.0487 | 0.0069 | 0.2405 | 0.4411 | 0.0473 | 4 | 0 |
| Fleiss_Kappa | 100 | 0.0025 | 5 | 0.8528 | 0.0498 | 0.0223 | 0.8062 | 0.9275 | 0.0606 | 0 | 0 | 0.5502 | 0.0431 | 0.0193 | 0.4982 | 0.5862 | 0.0749 | 0 | 0 | 0.7328 | 0.1252 | 0.056 | 0.5492 | 0.8359 | 0.1691 | 0 | 0 | 0.6404 | 0.0894 | 0.04 | 0.5612 | 0.7895 | 0.0559 | 1 | 0 |
| Fleiss_Kappa | 100 | 0.042 | 5 | 0.7291 | 0.0663 | 0.0297 | 0.623 | 0.7955 | 0.0499 | 0 | 1 | 0.5317 | 0.0188 | 0.0084 | 0.5124 | 0.5541 | 0.0321 | 0 | 0 | 0.6599 | 0.0459 | 0.0205 | 0.5945 | 0.7075 | 0.0595 | 0 | 0 | 0.555 | 0.0462 | 0.0207 | 0.5148 | 0.6278 | 0.0464 | 0 | 0 |
| Fleiss_Kappa | 100 | 0.0815 | 5 | 0.6259 | 0.052 | 0.0233 | 0.5564 | 0.6777 | 0.0797 | 0 | 0 | 0.501 | 0.0322 | 0.0144 | 0.4601 | 0.5386 | 0.0404 | 0 | 0 | 0.5738 | 0.067 | 0.0299 | 0.4875 | 0.6444 | 0.1035 | 0 | 0 | 0.5082 | 0.042 | 0.0188 | 0.4613 | 0.5624 | 0.0571 | 0 | 0 |
| Fleiss_Kappa | 100 | 0.121 | 5 | 0.5571 | 0.0524 | 0.0234 | 0.4898 | 0.6266 | 0.0571 | 0 | 0 | 0.4663 | 0.0255 | 0.0114 | 0.4292 | 0.4886 | 0.0333 | 0 | 0 | 0.5365 | 0.0579 | 0.0259 | 0.4565 | 0.5969 | 0.0797 | 0 | 0 | 0.481 | 0.0383 | 0.0171 | 0.4328 | 0.5219 | 0.0583 | 0 | 0 |
| Fleiss_Kappa | 100 | 0.1605 | 5 | 0.4829 | 0.0413 | 0.0185 | 0.4375 | 0.5422 | 0.0452 | 0 | 0 | 0.432 | 0.0343 | 0.0154 | 0.3947 | 0.4708 | 0.0559 | 0 | 0 | 0.4798 | 0.0773 | 0.0346 | 0.4018 | 0.5659 | 0.1439 | 0 | 0 | 0.4475 | 0.0302 | 0.0135 | 0.4149 | 0.4886 | 0.0392 | 0 | 0 |
| Fleiss_Kappa | 100 | 0.2 | 5 | 0.4732 | 0.0335 | 0.015 | 0.428 | 0.5172 | 0.0285 | 0 | 0 | 0.4161 | 0.0298 | 0.0133 | 0.3769 | 0.453 | 0.0321 | 0 | 0 | 0.4316 | 0.0701 | 0.0313 | 0.3436 | 0.5279 | 0.0678 | 0 | 0 | 0.3976 | 0.0241 | 0.0108 | 0.3614 | 0.4221 | 0.0274 | 0 | 0 |
| Fleiss_Kappa | 1080 | 0.0025 | 5 | 0.927 | 0.0384 | 0.0172 | 0.8668 | 0.9631 | 0.0381 | 0 | 0 | 0.8454 | 0.0305 | 0.0136 | 0.799 | 0.8763 | 0.0309 | 0 | 0 | 0.9006 | 0.0462 | 0.0206 | 0.8311 | 0.9596 | 0.0166 | 1 | 1 | 0.8673 | 0.0193 | 0.0086 | 0.8418 | 0.8858 | 0.0309 | 0 | 0 |
| Fleiss_Kappa | 1080 | 0.042 | 5 | 0.7505 | 0.0428 | 0.0191 | 0.6789 | 0.7907 | 0.0214 | 0 | 1 | 0.741 | 0.0421 | 0.0188 | 0.6754 | 0.7824 | 0.0392 | 0 | 0 | 0.7731 | 0.0606 | 0.0271 | 0.7194 | 0.8728 | 0.0464 | 1 | 0 | 0.7334 | 0.04 | 0.0179 | 0.7027 | 0.8002 | 0.0333 | 1 | 0 |
| Fleiss_Kappa | 1080 | 0.0815 | 5 | 0.6987 | 0.0255 | 0.0114 | 0.6551 | 0.7205 | 0.0107 | 0 | 1 | 0.658 | 0.0525 | 0.0235 | 0.6123 | 0.7194 | 0.088 | 0 | 0 | 0.6637 | 0.0687 | 0.0307 | 0.6076 | 0.7836 | 0.0036 | 1 | 1 | 0.649 | 0.0303 | 0.0136 | 0.6064 | 0.6849 | 0.0297 | 0 | 0 |
| Fleiss_Kappa | 1080 | 0.121 | 5 | 0.6506 | 0.0428 | 0.0191 | 0.6004 | 0.7146 | 0.0357 | 0 | 0 | 0.6178 | 0.0713 | 0.0319 | 0.5255 | 0.7063 | 0.0892 | 0 | 0 | 0.5824 | 0.0502 | 0.0224 | 0.5374 | 0.667 | 0.025 | 1 | 0 | 0.6023 | 0.0429 | 0.0192 | 0.5493 | 0.6468 | 0.0678 | 0 | 0 |
| Fleiss_Kappa | 1080 | 0.1605 | 5 | 0.5852 | 0.0629 | 0.0281 | 0.4815 | 0.6337 | 0.0559 | 0 | 1 | 0.5693 | 0.0688 | 0.0308 | 0.4601 | 0.6385 | 0.0535 | 0 | 1 | 0.5543 | 0.044 | 0.0197 | 0.4886 | 0.6111 | 0.0178 | 1 | 1 | 0.5243 | 0.0168 | 0.0075 | 0.5029 | 0.5469 | 0.0166 | 0 | 0 |
| Fleiss_Kappa | 1080 | 0.2 | 5 | 0.5177 | 0.0434 | 0.0194 | 0.4732 | 0.579 | 0.0583 | 0 | 0 | 0.5065 | 0.0685 | 0.0306 | 0.4292 | 0.6171 | 0.019 | 1 | 1 | 0.5017 | 0.0497 | 0.0222 | 0.4328 | 0.5671 | 0.0404 | 0 | 0 | 0.4753 | 0.0233 | 0.0104 | 0.4494 | 0.5101 | 0.0202 | 1 | 0 |
| Fleiss_Kappa | 2060 | 0.0025 | 5 | 0.9229 | 0.0307 | 0.0137 | 0.8728 | 0.9465 | 0.0262 | 0 | 1 | 0.8409 | 0.0139 | 0.0062 | 0.8264 | 0.8632 | 0.0095 | 1 | 0 | 0.9053 | 0.0202 | 0.009 | 0.8823 | 0.931 | 0.0274 | 0 | 0 | 0.8723 | 0.0112 | 0.005 | 0.8525 | 0.8799 | 0.0024 | 0 | 1 |
| Fleiss_Kappa | 2060 | 0.042 | 5 | 0.7755 | 0.0476 | 0.0213 | 0.7015 | 0.8204 | 0.0499 | 0 | 0 | 0.7717 | 0.0481 | 0.0215 | 0.7015 | 0.8276 | 0.0416 | 0 | 0 | 0.7536 | 0.0481 | 0.0215 | 0.7158 | 0.8359 | 0.0285 | 1 | 0 | 0.7346 | 0.0378 | 0.0169 | 0.6754 | 0.7741 | 0.0321 | 0 | 1 |
| Fleiss_Kappa | 2060 | 0.0815 | 5 | 0.7115 | 0.0285 | 0.0128 | 0.6623 | 0.7324 | 0.0155 | 0 | 1 | 0.6768 | 0.0402 | 0.018 | 0.6349 | 0.736 | 0.044 | 0 | 0 | 0.6682 | 0.0683 | 0.0305 | 0.6123 | 0.7859 | 0.0262 | 1 | 0 | 0.6627 | 0.032 | 0.0143 | 0.6206 | 0.6991 | 0.0428 | 0 | 0 |
| Fleiss_Kappa | 2060 | 0.121 | 5 | 0.6399 | 0.0453 | 0.0203 | 0.5814 | 0.6991 | 0.0488 | 0 | 0 | 0.6323 | 0.063 | 0.0282 | 0.5659 | 0.717 | 0.0892 | 0 | 0 | 0.5881 | 0.0488 | 0.0218 | 0.5445 | 0.6694 | 0.0285 | 1 | 0 | 0.6123 | 0.0357 | 0.016 | 0.5588 | 0.6516 | 0.0333 | 0 | 0 |
| Fleiss_Kappa | 2060 | 0.1605 | 5 | 0.5921 | 0.0592 | 0.0265 | 0.491 | 0.6314 | 0.0392 | 0 | 1 | 0.5724 | 0.0732 | 0.0328 | 0.4625 | 0.6504 | 0.0761 | 0 | 0 | 0.561 | 0.048 | 0.0215 | 0.4839 | 0.6135 | 0.0226 | 0 | 1 | 0.5205 | 0.0154 | 0.0069 | 0.4994 | 0.5374 | 0.0202 | 0 | 0 |
| Fleiss_Kappa | 2060 | 0.2 | 5 | 0.5196 | 0.0374 | 0.0167 | 0.4768 | 0.5683 | 0.0523 | 0 | 0 | 0.5139 | 0.0664 | 0.0297 | 0.447 | 0.6254 | 0.0095 | 1 | 1 | 0.4986 | 0.0505 | 0.0226 | 0.4256 | 0.5624 | 0.0392 | 0 | 0 | 0.4746 | 0.0169 | 0.0076 | 0.4601 | 0.497 | 0.025 | 0 | 0 |
| Fleiss_Kappa | 3040 | 0.0025 | 5 | 0.9275 | 0.0281 | 0.0126 | 0.8846 | 0.9572 | 0.0238 | 0 | 1 | 0.8544 | 0.0279 | 0.0125 | 0.805 | 0.8704 | 0.0083 | 0 | 1 | 0.9265 | 0.0277 | 0.0124 | 0.8954 | 0.9655 | 0.0297 | 0 | 0 | 0.9049 | 0.0225 | 0.0101 | 0.8716 | 0.9322 | 0.0178 | 0 | 0 |
| Fleiss_Kappa | 3040 | 0.042 | 5 | 0.7881 | 0.0451 | 0.0202 | 0.711 | 0.8288 | 0.0143 | 0 | 1 | 0.7724 | 0.0526 | 0.0235 | 0.7015 | 0.8359 | 0.063 | 0 | 0 | 0.7586 | 0.0461 | 0.0206 | 0.7241 | 0.8371 | 0.0321 | 1 | 0 | 0.7419 | 0.026 | 0.0116 | 0.7051 | 0.7752 | 0.0214 | 0 | 0 |
| Fleiss_Kappa | 3040 | 0.0815 | 5 | 0.716 | 0.0324 | 0.0145 | 0.6599 | 0.7431 | 0.0071 | 1 | 1 | 0.6832 | 0.0502 | 0.0225 | 0.6278 | 0.7538 | 0.0547 | 0 | 0 | 0.6713 | 0.0608 | 0.0272 | 0.6171 | 0.7705 | 0.0511 | 1 | 0 | 0.6715 | 0.0255 | 0.0114 | 0.6444 | 0.7003 | 0.044 | 0 | 0 |
| Fleiss_Kappa | 3040 | 0.121 | 5 | 0.638 | 0.049 | 0.0219 | 0.5731 | 0.7015 | 0.0488 | 0 | 0 | 0.6316 | 0.0592 | 0.0265 | 0.5671 | 0.7086 | 0.088 | 0 | 0 | 0.5926 | 0.0485 | 0.0217 | 0.5457 | 0.6718 | 0.0321 | 1 | 0 | 0.6109 | 0.0367 | 0.0164 | 0.5612 | 0.6504 | 0.0488 | 0 | 0 |
| Fleiss_Kappa | 3040 | 0.1605 | 5 | 0.5916 | 0.0618 | 0.0276 | 0.4851 | 0.6349 | 0.0333 | 0 | 1 | 0.5695 | 0.0773 | 0.0346 | 0.4506 | 0.6504 | 0.0737 | 0 | 0 | 0.5648 | 0.0483 | 0.0216 | 0.4886 | 0.6206 | 0.0214 | 1 | 1 | 0.5205 | 0.0178 | 0.008 | 0.497 | 0.5398 | 0.0262 | 0 | 0 |
| Fleiss_Kappa | 3040 | 0.2 | 5 | 0.5212 | 0.04 | 0.0179 | 0.4768 | 0.5755 | 0.0535 | 0 | 0 | 0.516 | 0.066 | 0.0295 | 0.447 | 0.6254 | 0.0166 | 1 | 1 | 0.501 | 0.0487 | 0.0218 | 0.4292 | 0.5624 | 0.0309 | 0 | 1 | 0.473 | 0.0188 | 0.0084 | 0.4542 | 0.497 | 0.0285 | 0 | 0 |
| Fleiss_Kappa | 4020 | 0.0025 | 5 | 0.9237 | 0.0371 | 0.0166 | 0.8668 | 0.9512 | 0.0428 | 0 | 0 | 0.8711 | 0.0189 | 0.0085 | 0.8418 | 0.8882 | 0.0214 | 0 | 0 | 0.9327 | 0.0316 | 0.0141 | 0.9001 | 0.981 | 0.0309 | 0 | 0 | 0.9061 | 0.0165 | 0.0074 | 0.887 | 0.9227 | 0.0297 | 0 | 0 |
| Fleiss_Kappa | 4020 | 0.042 | 5 | 0.7871 | 0.0357 | 0.016 | 0.7289 | 0.8181 | 0.0274 | 0 | 1 | 0.7722 | 0.0449 | 0.0201 | 0.7122 | 0.8311 | 0.0404 | 0 | 0 | 0.7631 | 0.0523 | 0.0234 | 0.7217 | 0.8502 | 0.0416 | 1 | 0 | 0.7477 | 0.0264 | 0.0118 | 0.7194 | 0.7752 | 0.044 | 0 | 0 |
| Fleiss_Kappa | 4020 | 0.0815 | 5 | 0.7172 | 0.0396 | 0.0177 | 0.6468 | 0.7384 | 0.0107 | 0 | 1 | 0.6837 | 0.0477 | 0.0213 | 0.6361 | 0.7526 | 0.0547 | 0 | 0 | 0.6666 | 0.0588 | 0.0263 | 0.6028 | 0.7538 | 0.0595 | 0 | 0 | 0.6708 | 0.0271 | 0.0121 | 0.6373 | 0.7039 | 0.0345 | 0 | 0 |
| Fleiss_Kappa | 4020 | 0.121 | 5 | 0.639 | 0.0468 | 0.0209 | 0.5802 | 0.7015 | 0.0476 | 0 | 0 | 0.6356 | 0.0632 | 0.0283 | 0.5719 | 0.7241 | 0.088 | 0 | 0 | 0.594 | 0.0464 | 0.0207 | 0.5517 | 0.6718 | 0.0262 | 1 | 0 | 0.6104 | 0.0342 | 0.0153 | 0.5624 | 0.6468 | 0.044 | 0 | 0 |
| Fleiss_Kappa | 4020 | 0.1605 | 5 | 0.5916 | 0.0618 | 0.0276 | 0.4851 | 0.6349 | 0.0333 | 0 | 1 | 0.5695 | 0.0773 | 0.0346 | 0.4506 | 0.6504 | 0.0737 | 0 | 0 | 0.5648 | 0.048 | 0.0214 | 0.4886 | 0.6206 | 0.0155 | 1 | 1 | 0.5212 | 0.0178 | 0.0079 | 0.497 | 0.5386 | 0.0262 | 0 | 0 |
| Fleiss_Kappa | 4020 | 0.2 | 5 | 0.5212 | 0.04 | 0.0179 | 0.4768 | 0.5755 | 0.0535 | 0 | 0 | 0.516 | 0.066 | 0.0295 | 0.447 | 0.6254 | 0.0166 | 1 | 1 | 0.501 | 0.0487 | 0.0218 | 0.4292 | 0.5624 | 0.0309 | 0 | 1 | 0.473 | 0.0186 | 0.0083 | 0.4542 | 0.4946 | 0.0309 | 0 | 0 |
| Fleiss_Kappa | 5000 | 0.0025 | 5 | 0.9215 | 0.0279 | 0.0125 | 0.8811 | 0.9441 | 0.0369 | 0 | 0 | 0.8737 | 0.0206 | 0.0092 | 0.8466 | 0.8989 | 0.025 | 0 | 0 | 0.9341 | 0.0233 | 0.0104 | 0.9084 | 0.9643 | 0.0309 | 0 | 0 | 0.9177 | 0.0121 | 0.0054 | 0.9001 | 0.9286 | 0.0166 | 0 | 0 |
| Fleiss_Kappa | 5000 | 0.042 | 5 | 0.7869 | 0.0395 | 0.0177 | 0.7253 | 0.8228 | 0.0392 | 0 | 0 | 0.7802 | 0.0427 | 0.0191 | 0.7241 | 0.8347 | 0.0428 | 0 | 0 | 0.7617 | 0.0436 | 0.0195 | 0.7158 | 0.8299 | 0.0369 | 1 | 0 | 0.7531 | 0.0274 | 0.0123 | 0.7182 | 0.7788 | 0.0464 | 0 | 0 |
| Fleiss_Kappa | 5000 | 0.0815 | 5 | 0.716 | 0.0388 | 0.0174 | 0.6468 | 0.7384 | 0.0036 | 0 | 1 | 0.6872 | 0.0454 | 0.0203 | 0.6409 | 0.7526 | 0.0547 | 0 | 0 | 0.6687 | 0.0568 | 0.0254 | 0.6064 | 0.7538 | 0.0535 | 0 | 0 | 0.6677 | 0.0239 | 0.0107 | 0.6373 | 0.6944 | 0.0345 | 0 | 0 |
| Fleiss_Kappa | 5000 | 0.121 | 5 | 0.639 | 0.0468 | 0.0209 | 0.5802 | 0.7015 | 0.0476 | 0 | 0 | 0.6359 | 0.0633 | 0.0283 | 0.5671 | 0.7241 | 0.0821 | 0 | 0 | 0.5935 | 0.0469 | 0.021 | 0.5493 | 0.6718 | 0.0262 | 1 | 0 | 0.6092 | 0.0347 | 0.0155 | 0.5612 | 0.6468 | 0.044 | 0 | 0 |
| Fleiss_Kappa | 5000 | 0.1605 | 5 | 0.5916 | 0.0618 | 0.0276 | 0.4851 | 0.6349 | 0.0333 | 0 | 1 | 0.5695 | 0.0773 | 0.0346 | 0.4506 | 0.6504 | 0.0737 | 0 | 0 | 0.5648 | 0.048 | 0.0214 | 0.4886 | 0.6206 | 0.0155 | 1 | 1 | 0.5215 | 0.0181 | 0.0081 | 0.497 | 0.5398 | 0.0262 | 0 | 0 |
| Fleiss_Kappa | 5000 | 0.2 | 5 | 0.5212 | 0.04 | 0.0179 | 0.4768 | 0.5755 | 0.0535 | 0 | 0 | 0.516 | 0.066 | 0.0295 | 0.447 | 0.6254 | 0.0166 | 1 | 1 | 0.501 | 0.0487 | 0.0218 | 0.4292 | 0.5624 | 0.0309 | 0 | 1 | 0.473 | 0.0186 | 0.0083 | 0.4542 | 0.4946 | 0.0309 | 0 | 0 |
| ICC | 100 | 0.0025 | 5 | 0.958 | 0.0106 | 0.0047 | 0.9457 | 0.9719 | 0.0143 | 0 | 0 | 0.9737 | 0.0099 | 0.0044 | 0.9587 | 0.9829 | 0.012 | 0 | 0 | 0.9862 | 0.004 | 0.0018 | 0.9817 | 0.9917 | 0.0048 | 0 | 0 | 0.9861 | 0.0028 | 0.0013 | 0.9837 | 0.9895 | 0.0047 | 0 | 0 |
| ICC | 100 | 0.042 | 5 | 0.8732 | 0.0141 | 0.0063 | 0.8554 | 0.8936 | 0.0107 | 0 | 0 | 0.8573 | 0.0263 | 0.0117 | 0.8243 | 0.89 | 0.0321 | 0 | 0 | 0.9268 | 0.0112 | 0.005 | 0.9092 | 0.9381 | 0.01 | 0 | 0 | 0.896 | 0.0107 | 0.0048 | 0.8841 | 0.9131 | 0.005 | 1 | 1 |
| ICC | 100 | 0.0815 | 5 | 0.8099 | 0.0223 | 0.01 | 0.7763 | 0.8302 | 0.0275 | 0 | 0 | 0.7924 | 0.0271 | 0.0121 | 0.7648 | 0.8256 | 0.0458 | 0 | 0 | 0.8704 | 0.0173 | 0.0078 | 0.8495 | 0.8872 | 0.0306 | 0 | 0 | 0.8274 | 0.01 | 0.0045 | 0.8141 | 0.8418 | 0.0049 | 1 | 1 |
| ICC | 100 | 0.121 | 5 | 0.7474 | 0.0203 | 0.0091 | 0.7127 | 0.7644 | 0.0082 | 0 | 1 | 0.7343 | 0.0315 | 0.0141 | 0.7039 | 0.7839 | 0.0305 | 0 | 0 | 0.8241 | 0.0253 | 0.0113 | 0.7902 | 0.8495 | 0.0371 | 0 | 0 | 0.7885 | 0.006 | 0.0027 | 0.7821 | 0.7943 | 0.0116 | 0 | 0 |
| ICC | 100 | 0.1605 | 5 | 0.699 | 0.0193 | 0.0086 | 0.6798 | 0.7235 | 0.0321 | 0 | 0 | 0.6894 | 0.0405 | 0.0181 | 0.6576 | 0.7465 | 0.0562 | 0 | 0 | 0.7698 | 0.028 | 0.0125 | 0.7357 | 0.7942 | 0.0475 | 0 | 0 | 0.736 | 0.0074 | 0.0033 | 0.7244 | 0.744 | 0.0051 | 0 | 1 |
| ICC | 100 | 0.2 | 5 | 0.6695 | 0.0072 | 0.0032 | 0.658 | 0.6768 | 0.0057 | 0 | 1 | 0.6541 | 0.0373 | 0.0167 | 0.6122 | 0.7074 | 0.0412 | 0 | 0 | 0.7311 | 0.0406 | 0.0182 | 0.6785 | 0.7678 | 0.0603 | 0 | 0 | 0.6779 | 0.024 | 0.0107 | 0.6417 | 0.6968 | 0.0307 | 0 | 0 |
| ICC | 1080 | 0.0025 | 5 | 0.9894 | 0.0022 | 0.001 | 0.9862 | 0.9923 | 0.0011 | 1 | 1 | 0.9885 | 0.0022 | 0.001 | 0.9861 | 0.9916 | 0.0027 | 0 | 0 | 0.9951 | 0.0006 | 0.0003 | 0.9941 | 0.9957 | 0.0008 | 0 | 0 | 0.993 | 0.0006 | 0.0003 | 0.9922 | 0.9938 | 0.0008 | 0 | 0 |
| ICC | 1080 | 0.042 | 5 | 0.9314 | 0.0064 | 0.0028 | 0.9264 | 0.9418 | 0.0062 | 0 | 0 | 0.9333 | 0.0083 | 0.0037 | 0.9277 | 0.9478 | 0.0037 | 1 | 0 | 0.9458 | 0.0052 | 0.0023 | 0.9382 | 0.9507 | 0.0066 | 0 | 0 | 0.9403 | 0.0072 | 0.0032 | 0.9294 | 0.9482 | 0.0061 | 0 | 1 |
| ICC | 1080 | 0.0815 | 5 | 0.8972 | 0.0089 | 0.004 | 0.8831 | 0.9065 | 0.0062 | 0 | 1 | 0.8862 | 0.0183 | 0.0082 | 0.8585 | 0.9007 | 0.0217 | 0 | 0 | 0.9057 | 0.0076 | 0.0034 | 0.8999 | 0.9166 | 0.0107 | 0 | 0 | 0.8924 | 0.0118 | 0.0053 | 0.8777 | 0.9063 | 0.0169 | 0 | 0 |
| ICC | 1080 | 0.121 | 5 | 0.8436 | 0.0146 | 0.0065 | 0.8283 | 0.8608 | 0.0246 | 0 | 0 | 0.836 | 0.031 | 0.0139 | 0.8048 | 0.8726 | 0.0551 | 0 | 0 | 0.8622 | 0.0108 | 0.0048 | 0.8448 | 0.8715 | 0.0106 | 0 | 0 | 0.8523 | 0.0154 | 0.0069 | 0.8294 | 0.8694 | 0.0151 | 0 | 0 |
| ICC | 1080 | 0.1605 | 5 | 0.7922 | 0.0216 | 0.0097 | 0.7582 | 0.8113 | 0.023 | 0 | 0 | 0.7931 | 0.0233 | 0.0104 | 0.7681 | 0.8188 | 0.042 | 0 | 0 | 0.8214 | 0.0128 | 0.0057 | 0.8007 | 0.8331 | 0.011 | 0 | 1 | 0.7985 | 0.0158 | 0.007 | 0.777 | 0.8157 | 0.0189 | 0 | 0 |

| | | | | | | | | | | | | | | | | | | | | | | | | | | | | | | | | | | | |
|---|---|---|---|---|---|---|---|---|---|---|---|---|---|---|---|---|---|---|---|---|---|---|---|---|---|---|---|---|---|---|---|---|---|---|---|
| ICC | 1080 | 0.2 | 5 | 0.7459 | 0.006 | 0.0027 | 0.739 | 0.7529 | 0.0095 | 0 | 0 | 0.736 | 0.0223 | 0.01 | 0.7077 | 0.7626 | 0.0283 | 0 | 0 | 0.7758 | 0.023 | 0.0103 | 0.7496 | 0.8099 | 0.0199 | 1 | 0 | 0.7413 | 0.02 | 0.009 | 0.7161 | 0.763 | 0.0314 | 0 | 0 |
| ICC | 2060 | 0.0025 | 5 | 0.9917 | 0.0017 | 0.0008 | 0.9889 | 0.9936 | 0.0006 | 1 | 1 | 0.9901 | 0.0013 | 0.0006 | 0.9889 | 0.9918 | 0.002 | 0 | 0 | 0.9953 | 0.0008 | 0.0004 | 0.9943 | 0.996 | 0.0016 | 0 | 0 | 0.9934 | 0.0003 | 0.0002 | 0.993 | 0.9939 | 0.0004 | 0 | 0 |
| ICC | 2060 | 0.042 | 5 | 0.943 | 0.0052 | 0.0023 | 0.9376 | 0.9505 | 0.0061 | 0 | 0 | 0.9417 | 0.0072 | 0.0032 | 0.9326 | 0.9525 | 0.0043 | 1 | 1 | 0.9524 | 0.0045 | 0.002 | 0.9452 | 0.9566 | 0.0043 | 0 | 0 | 0.9471 | 0.0069 | 0.0031 | 0.9368 | 0.9542 | 0.0075 | 0 | 0 |
| ICC | 2060 | 0.0815 | 5 | 0.9068 | 0.0073 | 0.0033 | 0.8956 | 0.9135 | 0.0087 | 0 | 0 | 0.8959 | 0.0169 | 0.0075 | 0.8692 | 0.9075 | 0.018 | 0 | 0 | 0.9118 | 0.0066 | 0.0029 | 0.9044 | 0.9203 | 0.0082 | 0 | 0 | 0.9019 | 0.0117 | 0.0052 | 0.8864 | 0.9146 | 0.0169 | 0 | 0 |
| ICC | 2060 | 0.121 | 5 | 0.8527 | 0.0124 | 0.0056 | 0.8409 | 0.8692 | 0.0198 | 0 | 0 | 0.8436 | 0.0294 | 0.0132 | 0.8079 | 0.8762 | 0.0472 | 0 | 0 | 0.8662 | 0.0099 | 0.0044 | 0.8509 | 0.8756 | 0.0106 | 0 | 0 | 0.8584 | 0.0161 | 0.0072 | 0.8355 | 0.8774 | 0.0157 | 0 | 0 |
| ICC | 2060 | 0.1605 | 5 | 0.799 | 0.022 | 0.0099 | 0.7645 | 0.8171 | 0.0247 | 0 | 0 | 0.7982 | 0.0221 | 0.0099 | 0.773 | 0.8204 | 0.0406 | 0 | 0 | 0.8251 | 0.0129 | 0.0058 | 0.8033 | 0.8358 | 0.009 | 0 | 1 | 0.8027 | 0.0161 | 0.0072 | 0.7822 | 0.8224 | 0.019 | 0 | 0 |
| ICC | 2060 | 0.2 | 5 | 0.7508 | 0.0064 | 0.0029 | 0.7432 | 0.7575 | 0.0099 | 0 | 0 | 0.74 | 0.0223 | 0.01 | 0.7108 | 0.7648 | 0.0306 | 0 | 0 | 0.7791 | 0.0234 | 0.0105 | 0.752 | 0.8135 | 0.0199 | 1 | 0 | 0.7441 | 0.0215 | 0.0096 | 0.7153 | 0.7675 | 0.0306 | 0 | 0 |
| ICC | 3040 | 0.0025 | 5 | 0.9926 | 0.0013 | 0.0006 | 0.9904 | 0.9936 | 0.0007 | 0 | 1 | 0.9916 | 0.0007 | 0.0003 | 0.9908 | 0.9925 | 0.0008 | 0 | 0 | 0.9956 | 0.0005 | 0.0002 | 0.9949 | 0.9961 | 0.0007 | 0 | 0 | 0.9941 | 0.0002 | 0.0001 | 0.9937 | 0.9943 | 0.0002 | 0 | 0 |
| ICC | 3040 | 0.042 | 5 | 0.9471 | 0.0043 | 0.0019 | 0.9431 | 0.9535 | 0.0051 | 0 | 0 | 0.9453 | 0.007 | 0.0031 | 0.9346 | 0.9543 | 0.002 | 1 | 1 | 0.9548 | 0.0034 | 0.0015 | 0.9495 | 0.9577 | 0.0041 | 0 | 0 | 0.9498 | 0.0062 | 0.0028 | 0.9405 | 0.9556 | 0.0076 | 0 | 0 |
| ICC | 3040 | 0.0815 | 5 | 0.9099 | 0.0066 | 0.0029 | 0.9002 | 0.9159 | 0.0087 | 0 | 0 | 0.8995 | 0.0161 | 0.0072 | 0.8738 | 0.9105 | 0.0173 | 0 | 0 | 0.9137 | 0.0066 | 0.0029 | 0.9064 | 0.9219 | 0.0093 | 0 | 0 | 0.9037 | 0.0119 | 0.0053 | 0.8884 | 0.9173 | 0.0172 | 0 | 0 |
| ICC | 3040 | 0.121 | 5 | 0.8545 | 0.0121 | 0.0054 | 0.8427 | 0.8708 | 0.0184 | 0 | 0 | 0.8454 | 0.0286 | 0.0128 | 0.8104 | 0.8771 | 0.0452 | 0 | 0 | 0.867 | 0.01 | 0.0045 | 0.8516 | 0.8767 | 0.0107 | 0 | 0 | 0.8592 | 0.016 | 0.0072 | 0.8363 | 0.8782 | 0.0147 | 0 | 0 |
| ICC | 3040 | 0.1605 | 5 | 0.7998 | 0.0222 | 0.0099 | 0.7653 | 0.8178 | 0.0256 | 0 | 0 | 0.799 | 0.0218 | 0.0097 | 0.7737 | 0.8207 | 0.0394 | 0 | 0 | 0.8256 | 0.013 | 0.0058 | 0.8038 | 0.8365 | 0.0093 | 0 | 1 | 0.8033 | 0.0161 | 0.0072 | 0.7829 | 0.8231 | 0.019 | 0 | 0 |
| ICC | 3040 | 0.2 | 5 | 0.7511 | 0.0065 | 0.0029 | 0.7436 | 0.7578 | 0.0102 | 0 | 0 | 0.7403 | 0.0224 | 0.01 | 0.7109 | 0.7649 | 0.0307 | 0 | 0 | 0.7794 | 0.0235 | 0.0105 | 0.7522 | 0.8138 | 0.0199 | 1 | 0 | 0.7444 | 0.0216 | 0.0097 | 0.7153 | 0.7677 | 0.0304 | 0 | 0 |
| ICC | 4020 | 0.0025 | 5 | 0.9932 | 0.0012 | 0.0005 | 0.9912 | 0.9941 | 0.0007 | 0 | 1 | 0.9923 | 0.0007 | 0.0003 | 0.9915 | 0.9931 | 0.0012 | 0 | 0 | 0.9957 | 0.0005 | 0.0002 | 0.9948 | 0.9961 | 0.0005 | 0 | 0 | 0.9944 | 0.0001 | 0.0001 | 0.9942 | 0.9946 | 0.0001 | 0 | 0 |
| ICC | 4020 | 0.042 | 5 | 0.9491 | 0.0041 | 0.0018 | 0.9461 | 0.9553 | 0.0048 | 0 | 0 | 0.9475 | 0.0071 | 0.0032 | 0.936 | 0.9553 | 0.0027 | 1 | 1 | 0.9558 | 0.0034 | 0.0015 | 0.9507 | 0.9591 | 0.0037 | 0 | 0 | 0.9508 | 0.0059 | 0.0027 | 0.9422 | 0.9563 | 0.008 | 0 | 0 |
| ICC | 4020 | 0.0815 | 5 | 0.9108 | 0.0063 | 0.0028 | 0.9018 | 0.9167 | 0.0087 | 0 | 0 | 0.9003 | 0.0162 | 0.0072 | 0.8744 | 0.9114 | 0.0168 | 0 | 0 | 0.9142 | 0.0067 | 0.003 | 0.9066 | 0.9225 | 0.0097 | 0 | 0 | 0.9042 | 0.0122 | 0.0055 | 0.8885 | 0.9182 | 0.0176 | 0 | 0 |
| ICC | 4020 | 0.121 | 5 | 0.8548 | 0.0121 | 0.0054 | 0.8427 | 0.871 | 0.0182 | 0 | 0 | 0.8459 | 0.0284 | 0.0127 | 0.8109 | 0.8772 | 0.0448 | 0 | 0 | 0.8671 | 0.0101 | 0.0045 | 0.8517 | 0.8768 | 0.0111 | 0 | 0 | 0.8595 | 0.016 | 0.0072 | 0.8365 | 0.8785 | 0.0145 | 0 | 0 |
| ICC | 4020 | 0.1605 | 5 | 0.8 | 0.0222 | 0.0099 | 0.7655 | 0.8178 | 0.0256 | 0 | 0 | 0.7992 | 0.0217 | 0.0097 | 0.7739 | 0.8208 | 0.0393 | 0 | 0 | 0.8256 | 0.013 | 0.0058 | 0.8038 | 0.8366 | 0.0091 | 0 | 1 | 0.8033 | 0.0161 | 0.0072 | 0.7831 | 0.8232 | 0.019 | 0 | 0 |
| ICC | 4020 | 0.2 | 5 | 0.7512 | 0.0065 | 0.0029 | 0.7436 | 0.7578 | 0.0102 | 0 | 0 | 0.7403 | 0.0224 | 0.01 | 0.7109 | 0.7649 | 0.0307 | 0 | 0 | 0.7794 | 0.0234 | 0.0105 | 0.7522 | 0.8138 | 0.0198 | 1 | 0 | 0.7444 | 0.0216 | 0.0097 | 0.7154 | 0.7677 | 0.0304 | 0 | 0 |
| ICC | 5000 | 0.0025 | 5 | 0.9936 | 0.0009 | 0.0004 | 0.9922 | 0.9945 | 0.0012 | 0 | 0 | 0.993 | 0.0006 | 0.0003 | 0.9922 | 0.9937 | 0.0005 | 0 | 0 | 0.9958 | 0.0005 | 0.0002 | 0.995 | 0.9963 | 0.0004 | 0 | 1 | 0.9948 | 0.0001 | 0 | 0.9947 | 0.9949 | 0.0001 | 0 | 0 |
| ICC | 5000 | 0.042 | 5 | 0.9501 | 0.004 | 0.0018 | 0.9469 | 0.9562 | 0.0048 | 0 | 0 | 0.9488 | 0.007 | 0.0031 | 0.9375 | 0.9564 | 0.002 | 1 | 1 | 0.9563 | 0.0033 | 0.0015 | 0.9514 | 0.9597 | 0.0037 | 0 | 0 | 0.9517 | 0.0059 | 0.0026 | 0.943 | 0.957 | 0.008 | 0 | 0 |
| ICC | 5000 | 0.0815 | 5 | 0.9111 | 0.0063 | 0.0028 | 0.9023 | 0.9171 | 0.0088 | 0 | 0 | 0.9007 | 0.0161 | 0.0072 | 0.8747 | 0.9116 | 0.0164 | 0 | 0 | 0.9143 | 0.0068 | 0.003 | 0.9066 | 0.9227 | 0.01 | 0 | 0 | 0.9044 | 0.0123 | 0.0055 | 0.8886 | 0.9187 | 0.0174 | 0 | 0 |
| ICC | 5000 | 0.121 | 5 | 0.8548 | 0.0121 | 0.0054 | 0.8428 | 0.8711 | 0.0182 | 0 | 0 | 0.846 | 0.0284 | 0.0127 | 0.8111 | 0.8773 | 0.0448 | 0 | 0 | 0.8672 | 0.0101 | 0.0045 | 0.8518 | 0.8768 | 0.0111 | 0 | 0 | 0.8595 | 0.016 | 0.0072 | 0.8366 | 0.8786 | 0.0145 | 0 | 0 |
| ICC | 5000 | 0.1605 | 5 | 0.8 | 0.0222 | 0.0099 | 0.7655 | 0.8178 | 0.0256 | 0 | 0 | 0.7992 | 0.0217 | 0.0097 | 0.7739 | 0.8208 | 0.0393 | 0 | 0 | 0.8256 | 0.013 | 0.0058 | 0.8038 | 0.8366 | 0.0092 | 0 | 1 | 0.8033 | 0.0161 | 0.0072 | 0.7831 | 0.8232 | 0.019 | 0 | 0 |
| ICC | 5000 | 0.2 | 5 | 0.7512 | 0.0065 | 0.0029 | 0.7436 | 0.7578 | 0.0102 | 0 | 0 | 0.7403 | 0.0223 | 0.01 | 0.7109 | 0.7649 | 0.0307 | 0 | 0 | 0.7794 | 0.0234 | 0.0105 | 0.7522 | 0.8138 | 0.0198 | 1 | 0 | 0.7444 | 0.0216 | 0.0097 | 0.7154 | 0.7677 | 0.0304 | 0 | 0 |
| NDCG@20% | 100 | 0.0025 | 50 | 0.9012 | 0.0157 | 0.0022 | 0.8805 | 0.936 | 0.011 | 9 | 0 | 0.8652 | 0.0203 | 0.0029 | 0.8208 | 0.9097 | 0.0315 | 0 | 0 | 0.8927 | 0.0165 | 0.0023 | 0.8644 | 0.9263 | 0.0234 | 0 | 0 | 0.9023 | 0.0212 | 0.003 | 0.8713 | 0.9459 | 0.0322 | 0 | 0 |
| NDCG@20% | 100 | 0.042 | 50 | 0.9043 | 0.0127 | 0.0018 | 0.8829 | 0.9311 | 0.0152 | 0 | 0 | 0.8705 | 0.0219 | 0.0031 | 0.8203 | 0.9107 | 0.0398 | 0 | 0 | 0.9032 | 0.0176 | 0.0025 | 0.8587 | 0.9324 | 0.0233 | 0 | 0 | 0.9013 | 0.0172 | 0.0024 | 0.8591 | 0.9434 | 0.0233 | 0 | 0 |
| NDCG@20% | 100 | 0.0815 | 50 | 0.9061 | 0.0139 | 0.002 | 0.8698 | 0.9329 | 0.0188 | 0 | 0 | 0.8736 | 0.0231 | 0.0033 | 0.8263 | 0.9145 | 0.0348 | 0 | 0 | 0.9028 | 0.0178 | 0.0025 | 0.8596 | 0.9306 | 0.0292 | 0 | 0 | 0.9032 | 0.0156 | 0.0022 | 0.8779 | 0.938 | 0.0234 | 0 | 0 |
| NDCG@20% | 100 | 0.121 | 50 | 0.9039 | 0.0118 | 0.0017 | 0.8815 | 0.9335 | 0.0187 | 0 | 0 | 0.8746 | 0.0183 | 0.0026 | 0.8263 | 0.9169 | 0.0189 | 1 | 1 | 0.8971 | 0.0206 | 0.0029 | 0.8549 | 0.9373 | 0.0332 | 0 | 0 | 0.902 | 0.0179 | 0.0025 | 0.8622 | 0.9328 | 0.0295 | 0 | 0 |
| NDCG@20% | 100 | 0.1605 | 50 | 0.9049 | 0.0124 | 0.0018 | 0.8746 | 0.9248 | 0.0174 | 0 | 0 | 0.8767 | 0.0195 | 0.0028 | 0.8382 | 0.9158 | 0.028 | 0 | 0 | 0.8967 | 0.0223 | 0.0031 | 0.8505 | 0.9441 | 0.0371 | 0 | 0 | 0.9034 | 0.0172 | 0.0024 | 0.8624 | 0.9319 | 0.0277 | 0 | 0 |
| NDCG@20% | 100 | 0.2 | 50 | 0.9007 | 0.0128 | 0.0018 | 0.8784 | 0.9303 | 0.0209 | 0 | 0 | 0.8719 | 0.0175 | 0.0025 | 0.8355 | 0.912 | 0.0207 | 0 | 0 | 0.8953 | 0.0175 | 0.0025 | 0.8532 | 0.9329 | 0.027 | 0 | 0 | 0.9023 | 0.0167 | 0.0024 | 0.8673 | 0.9323 | 0.0252 | 0 | 0 |
| NDCG@20% | 1080 | 0.0025 | 50 | 0.9037 | 0.0143 | 0.002 | 0.8832 | 0.9322 | 0.0134 | 8 | 0 | 0.8725 | 0.0206 | 0.0029 | 0.8383 | 0.9069 | 0.0345 | 0 | 0 | 0.9054 | 0.0152 | 0.0022 | 0.8775 | 0.9355 | 0.0222 | 0 | 0 | 0.9035 | 0.0168 | 0.0024 | 0.8815 | 0.9387 | 0.0268 | 0 | 0 |
| NDCG@20% | 1080 | 0.042 | 50 | 0.9065 | 0.0101 | 0.0014 | 0.8809 | 0.9254 | 0.0132 | 0 | 0 | 0.8788 | 0.0169 | 0.0024 | 0.8468 | 0.9116 | 0.0272 | 0 | 0 | 0.9009 | 0.0219 | 0.0031 | 0.8579 | 0.9373 | 0.0379 | 0 | 0 | 0.9147 | 0.0148 | 0.0021 | 0.89 | 0.9455 | 0.0214 | 0 | 0 |
| NDCG@20% | 1080 | 0.0815 | 50 | 0.9072 | 0.011 | 0.0016 | 0.8834 | 0.9307 | 0.0153 | 0 | 0 | 0.8787 | 0.0164 | 0.0023 | 0.8458 | 0.909 | 0.0232 | 0 | 0 | 0.8967 | 0.0226 | 0.0032 | 0.8529 | 0.9337 | 0.0383 | 0 | 0 | 0.9147 | 0.0151 | 0.0021 | 0.887 | 0.9416 | 0.026 | 0 | 0 |
| NDCG@20% | 1080 | 0.121 | 50 | 0.9049 | 0.0118 | 0.0017 | 0.8777 | 0.9273 | 0.0132 | 0 | 1 | 0.8789 | 0.0172 | 0.0024 | 0.8386 | 0.9023 | 0.0249 | 0 | 0 | 0.8941 | 0.0209 | 0.0029 | 0.8454 | 0.9346 | 0.0294 | 0 | 0 | 0.9143 | 0.0153 | 0.0022 | 0.8828 | 0.9421 | 0.0203 | 0 | 0 |
| NDCG@20% | 1080 | 0.1605 | 50 | 0.9043 | 0.0124 | 0.0018 | 0.8698 | 0.9252 | 0.014 | 0 | 2 | 0.8771 | 0.018 | 0.0025 | 0.8376 | 0.9092 | 0.027 | 0 | 0 | 0.8949 | 0.0209 | 0.0029 | 0.8385 | 0.9315 | 0.0276 | 0 | 1 | 0.9167 | 0.0165 | 0.0023 | 0.8829 | 0.941 | 0.0266 | 0 | 0 |
| NDCG@20% | 1080 | 0.2 | 50 | 0.9061 | 0.0138 | 0.0019 | 0.8768 | 0.931 | 0.0179 | 0 | 0 | 0.8758 | 0.0161 | 0.0023 | 0.8451 | 0.9134 | 0.0249 | 0 | 0 | 0.8911 | 0.0187 | 0.0026 | 0.8558 | 0.9381 | 0.0246 | 0 | 0 | 0.9167 | 0.015 | 0.0021 | 0.8842 | 0.9469 | 0.0192 | 0 | 0 |
| NDCG@20% | 2060 | 0.0025 | 50 | 0.9054 | 0.0134 | 0.0019 | 0.8868 | 0.9327 | 0.0183 | 0 | 0 | 0.8755 | 0.02 | 0.0028 | 0.8419 | 0.9137 | 0.0354 | 0 | 0 | 0.9087 | 0.0155 | 0.0022 | 0.8778 | 0.9422 | 0.0191 | 0 | 0 | 0.9036 | 0.014 | 0.002 | 0.8839 | 0.9308 | 0.0177 | 0 | 0 |
| NDCG@20% | 2060 | 0.042 | 50 | 0.9077 | 0.0098 | 0.0014 | 0.8828 | 0.9256 | 0.0126 | 0 | 0 | 0.8798 | 0.0176 | 0.0025 | 0.8387 | 0.9099 | 0.0275 | 0 | 0 | 0.8986 | 0.0215 | 0.003 | 0.8586 | 0.939 | 0.0339 | 0 | 0 | 0.9164 | 0.015 | 0.0021 | 0.8905 | 0.9467 | 0.0253 | 0 | 0 |
| NDCG@20% | 2060 | 0.0815 | 50 | 0.9074 | 0.0114 | 0.0016 | 0.8798 | 0.9272 | 0.016 | 0 | 0 | 0.8787 | 0.0163 | 0.0023 | 0.8432 | 0.9089 | 0.0247 | 0 | 0 | 0.8965 | 0.0219 | 0.0031 | 0.8512 | 0.9303 | 0.0389 | 0 | 0 | 0.9175 | 0.015 | 0.0021 | 0.8881 | 0.9437 | 0.0259 | 0 | 0 |
| NDCG@20% | 2060 | 0.121 | 50 | 0.9046 | 0.0128 | 0.0018 | 0.8759 | 0.9272 | 0.017 | 0 | 0 | 0.8795 | 0.0169 | 0.0024 | 0.8398 | 0.9044 | 0.0238 | 0 | 0 | 0.8936 | 0.0208 | 0.0029 | 0.8466 | 0.9351 | 0.033 | 0 | 0 | 0.9155 | 0.0154 | 0.0022 | 0.8829 | 0.9436 | 0.0205 | 0 | 0 |
| NDCG@20% | 2060 | 0.1605 | 50 | 0.9045 | 0.0135 | 0.0019 | 0.872 | 0.9309 | 0.0143 | 0 | 3 | 0.8786 | 0.0174 | 0.0025 | 0.845 | 0.9136 | 0.0277 | 0 | 0 | 0.894 | 0.0201 | 0.0028 | 0.8393 | 0.9308 | 0.0289 | 0 | 0 | 0.9167 | 0.016 | 0.0023 | 0.8877 | 0.9426 | 0.0264 | 0 | 0 |
| NDCG@20% | 2060 | 0.2 | 50 | 0.9049 | 0.0141 | 0.002 | 0.8764 | 0.9311 | 0.0184 | 0 | 0 | 0.8757 | 0.0162 | 0.0023 | 0.844 | 0.9139 | 0.0267 | 0 | 0 | 0.8907 | 0.0192 | 0.0027 | 0.8551 | 0.938 | 0.0281 | 0 | 0 | 0.9161 | 0.0161 | 0.0023 | 0.8841 | 0.9485 | 0.0198 | 0 | 0 |
| NDCG@20% | 3040 | 0.0025 | 50 | 0.9062 | 0.0127 | 0.0018 | 0.8849 | 0.9304 | 0.0161 | 0 | 0 | 0.8773 | 0.0197 | 0.0028 | 0.8487 | 0.9106 | 0.0355 | 0 | 0 | 0.9108 | 0.0188 | 0.0027 | 0.873 | 0.9374 | 0.0356 | 0 | 0 | 0.9027 | 0.0142 | 0.002 | 0.8813 | 0.9309 | 0.0197 | 0 | 0 |
| NDCG@20% | 3040 | 0.042 | 50 | 0.9085 | 0.0115 | 0.0016 | 0.8792 | 0.9325 | 0.0113 | 1 | 3 | 0.8801 | 0.0181 | 0.0026 | 0.8463 | 0.9077 | 0.0324 | 0 | 0 | 0.8972 | 0.0215 | 0.003 | 0.8572 | 0.9396 | 0.0342 | 0 | 0 | 0.9166 | 0.0159 | 0.0022 | 0.8901 | 0.9446 | 0.0252 | 0 | 0 |
| NDCG@20% | 3040 | 0.0815 | 50 | 0.9073 | 0.0121 | 0.0017 | 0.88 | 0.9282 | 0.0164 | 0 | 0 | 0.8787 | 0.0165 | 0.0023 | 0.8445 | 0.9084 | 0.0269 | 0 | 0 | 0.8969 | 0.0212 | 0.003 | 0.8525 | 0.9328 | 0.0349 | 0 | 0 | 0.9169 | 0.0149 | 0.0021 | 0.8848 | 0.9422 | 0.0218 | 0 | 0 |
| NDCG@20% | 3040 | 0.121 | 50 | 0.9045 | 0.013 | 0.0018 | 0.8759 | 0.9272 | 0.0169 | 0 | 0 | 0.8806 | 0.0161 | 0.0023 | 0.8397 | 0.9045 | 0.0228 | 0 | 0 | 0.8933 | 0.0206 | 0.0029 | 0.8462 | 0.9352 | 0.0322 | 0 | 0 | 0.9154 | 0.0156 | 0.0022 | 0.8829 | 0.946 | 0.0193 | 0 | 0 |
| NDCG@20% | 3040 | 0.1605 | 50 | 0.9042 | 0.0135 | 0.0019 | 0.8733 | 0.9309 | 0.0149 | 0 | 2 | 0.8786 | 0.0171 | 0.0024 | 0.8454 | 0.9122 | 0.0275 | 0 | 0 | 0.8937 | 0.0201 | 0.0028 | 0.8394 | 0.9308 | 0.0294 | 0 | 0 | 0.9168 | 0.0163 | 0.0023 | 0.8877 | 0.9435 | 0.0255 | 0 | 0 |
| NDCG@20% | 3040 | 0.2 | 50 | 0.9054 | 0.0138 | 0.002 | 0.8761 | 0.9311 | 0.0176 | 0 | 0 | 0.8758 | 0.016 | 0.0023 | 0.844 | 0.9139 | 0.0261 | 0 | 0 | 0.8905 | 0.0191 | 0.0027 | 0.8551 | 0.9382 | 0.0278 | 0 | 0 | 0.9161 | 0.0161 | 0.0023 | 0.8841 | 0.9485 | 0.0201 | 0 | 0 |
| NDCG@20% | 4020 | 0.0025 | 50 | 0.9071 | 0.0115 | 0.0016 | 0.8902 | 0.9265 | 0.0169 | 0 | 0 | 0.8779 | 0.0202 | 0.0029 | 0.8511 | 0.9163 | 0.0334 | 0 | 0 | 0.9109 | 0.0194 | 0.0027 | 0.8799 | 0.9421 | 0.0376 | 0 | 0 | 0.9027 | 0.0148 | 0.0021 | 0.8845 | 0.9319 | 0.023 | 0 | 0 |
| NDCG@20% | 4020 | 0.042 | 50 | 0.9083 | 0.0132 | 0.0019 | 0.8778 | 0.9329 | 0.0141 | 0 | 2 | 0.8802 | 0.018 | 0.0025 | 0.8503 | 0.9084 | 0.0338 | 0 | 0 | 0.8966 | 0.0225 | 0.0032 | 0.8604 | 0.9456 | 0.0375 | 0 | 0 | 0.9172 | 0.0158 | 0.0022 | 0.89 | 0.9451 | 0.0235 | 0 | 0 |
| NDCG@20% | 4020 | 0.0815 | 50 | 0.9078 | 0.0129 | 0.0018 | 0.8799 | 0.9303 | 0.0175 | 0 | 0 | 0.8787 | 0.0161 | 0.0023 | 0.844 | 0.9087 | 0.0254 | 0 | 0 | 0.8965 | 0.0212 | 0.003 | 0.8521 | 0.9329 | 0.033 | 0 | 0 | 0.9165 | 0.0151 | 0.0021 | 0.8849 | 0.9421 | 0.0226 | 0 | 0 |
| NDCG@20% | 4020 | 0.121 | 50 | 0.9046 | 0.0126 | 0.0018 | 0.8759 | 0.9249 | 0.0166 | 0 | 0 | 0.8807 | 0.0158 | 0.0022 | 0.8397 | 0.9044 | 0.0241 | 0 | 0 | 0.893 | 0.0206 | 0.0029 | 0.8462 | 0.9355 | 0.0322 | 0 | 0 | 0.9155 | 0.0156 | 0.0022 | 0.8831 | 0.9457 | 0.0199 | 0 | 0 |
| NDCG@20% | 4020 | 0.1605 | 50 | 0.9045 | 0.0136 | 0.0019 | 0.8733 | 0.9309 | 0.0147 | 0 | 2 | 0.8787 | 0.0172 | 0.0024 | 0.8454 | 0.9122 | 0.0278 | 0 | 0 | 0.8936 | 0.0201 | 0.0028 | 0.8394 | 0.9308 | 0.0294 | 0 | 0 | 0.917 | 0.0163 | 0.0023 | 0.8877 | 0.9435 | 0.0252 | 0 | 0 |
| NDCG@20% | 4020 | 0.2 | 50 | 0.9055 | 0.0138 | 0.0019 | 0.8761 | 0.9311 | 0.0171 | 0 | 0 | 0.876 | 0.0159 | 0.0023 | 0.844 | 0.9139 | 0.0261 | 0 | 0 | 0.8905 | 0.0191 | 0.0027 | 0.8551 | 0.9382 | 0.0278 | 0 | 0 | 0.9161 | 0.0161 | 0.0023 | 0.8841 | 0.9485 | 0.0201 | 0 | 0 |
| NDCG@20% | 5000 | 0.0025 | 50 | 0.9088 | 0.0102 | 0.0014 | 0.889 | 0.9253 | 0.0152 | 0 | 0 | 0.8781 | 0.0207 | 0.0029 | 0.852 | 0.9159 | 0.0346 | 0 | 0 | 0.9119 | 0.02 | 0.0028 | 0.878 | 0.9407 | 0.035 | 0 | 0 | 0.9043 | 0.0147 | 0.0021 | 0.8852 | 0.9349 | 0.0237 | 0 | 0 |
| NDCG@20% | 5000 | 0.042 | 50 | 0.9081 | 0.0136 | 0.0019 | 0.8779 | 0.9348 | 0.0151 | 0 | 2 | 0.8805 | 0.0179 | 0.0025 | 0.8532 | 0.909 | 0.0323 | 0 | 0 | 0.8964 | 0.0227 | 0.0032 | 0.8509 | 0.9447 | 0.0354 | 0 | 0 | 0.9173 | 0.0161 | 0.0023 | 0.8898 | 0.9459 | 0.0241 | 0 | 0 |
| NDCG@20% | 5000 | 0.0815 | 50 | 0.9079 | 0.0131 | 0.0019 | 0.8799 | 0.9305 | 0.018 | 0 | 0 | 0.8791 | 0.0163 | 0.0023 | 0.844 | 0.9087 | 0.0254 | 0 | 0 | 0.8963 | 0.0212 | 0.003 | 0.8521 | 0.9326 | 0.033 | 0 | 0 | 0.9167 | 0.0153 | 0.0022 | 0.8845 | 0.9423 | 0.0228 | 0 | 0 |
| NDCG@20% | 5000 | 0.121 | 50 | 0.9046 | 0.0127 | 0.0018 | 0.8759 | 0.9249 | 0.0167 | 0 | 0 | 0.8807 | 0.0159 | 0.0022 | 0.8397 | 0.9045 | 0.0241 | 0 | 0 | 0.8929 | 0.0205 | 0.0029 | 0.8462 | 0.9355 | 0.0322 | 0 | 0 | 0.9156 | 0.0157 | 0.0022 | 0.8831 | 0.9456 | 0.0205 | 0 | 0 |
| NDCG@20% | 5000 | 0.1605 | 50 | 0.9045 | 0.0135 | 0.0019 | 0.8733 | 0.9309 | 0.0147 | 0 | 2 | 0.8787 | 0.0171 | 0.0024 | 0.8454 | 0.9122 | 0.0272 | 0 | 0 | 0.8936 | 0.0201 | 0.0028 | 0.8394 | 0.9308 | 0.0294 | 0 | 0 | 0.9169 | 0.0163 | 0.0023 | 0.8877 | 0.9435 | 0.0252 | 0 | 0 |
| NDCG@20% | 5000 | 0.2 | 50 | 0.9055 | 0.0138 | 0.0019 | 0.8761 | 0.9311 | 0.0171 | 0 | 0 | 0.876 | 0.0159 | 0.0023 | 0.844 | 0.9139 | 0.0261 | 0 | 0 | 0.8905 | 0.0191 | 0.0027 | 0.8551 | 0.9382 | 0.0278 | 0 | 0 | 0.9161 | 0.016 | 0.0023 | 0.8841 | 0.9476 | 0.0201 | 0 | 0 |
| Pearson | 100 | 0.0025 | 50 | 0.2012 | 0.0777 | 0.011 | 0.0903 | 0.3198 | 0.1362 | 0 | 0 | 0.1136 | 0.076 | 0.0107 | -0.0146 | 0.2463 | 0.0937 | 0 | 0 | 0.3386 | 0.0903 | 0.0128 | 0.206 | 0.4959 | 0.153 | 0 | 0 | 0.3677 | 0.0996 | 0.0141 | 0.226 | 0.5432 | 0.1257 | 0 | 0 |
| Pearson | 100 | 0.042 | 50 | 0.2339 | 0.0533 | 0.0075 | 0.1194 | 0.3278 | 0.0841 | 0 | 0 | 0.1367 | 0.075 | 0.0106 | -0.0249 | 0.2843 | 0.0938 | 0 | 0 | 0.4943 | 0.0439 | 0.0062 | 0.3801 | 0.5626 | 0.0562 | 0 | 2 | 0.4091 | 0.1064 | 0.015 | 0.23 | 0.6063 | 0.1697 | 0 | 0 |
| Pearson | 100 | 0.0815 | 50 | 0.2348 | 0.0552 | 0.0078 | 0.1097 | 0.33 | 0.0673 | 0 | 0 | 0.1378 | 0.0759 | 0.0107 | -0.0348 | 0.2854 | 0.1111 | 0 | 0 | 0.5103 | 0.0505 | 0.0071 | 0.3658 | 0.5793 | 0.0469 | 0 | 6 | 0.4019 | 0.0947 | 0.0134 | 0.2081 | 0.5915 | 0.1428 | 0 | 0 |
| Pearson | 100 | 0.121 | 50 | 0.2107 | 0.0525 | 0.0074 | 0.0906 | 0.3094 | 0.0611 | 0 | 3 | 0.1358 | 0.0689 | 0.0097 | -0.039 | 0.3045 | 0.0801 | 1 | 1 | 0.4948 | 0.0588 | 0.0083 | 0.3439 | 0.5943 | 0.0776 | 0 | 0 | 0.3853 | 0.1009 | 0.0143 | 0.1979 | 0.5609 | 0.142 | 0 | 0 |
| Pearson | 100 | 0.1605 | 50 | 0.1949 | 0.0596 | 0.0084 | 0.0591 | 0.3156 | 0.0729 | 0 | 0 | 0.1376 | 0.0765 | 0.0108 | -0.0194 | 0.3092 | 0.0992 | 0 | 0 | 0.4905 | 0.061 | 0.0086 | 0.3033 | 0.614 | 0.0815 | 0 | 1 | 0.3875 | 0.0898 | 0.0127 | 0.1923 | 0.567 | 0.1371 | 0 | 0 |
| Pearson | 100 | 0.2 | 50 | 0.1863 | 0.0581 | 0.0082 | 0.0607 | 0.2981 | 0.0748 | 0 | 0 | 0.1292 | 0.0727 | 0.0103 | -0.0031 | 0.3188 | 0.0997 | 0 | 0 | 0.4902 | 0.0755 | 0.0107 | 0.3304 | 0.6423 | 0.1072 | 0 | 0 | 0.3774 | 0.0913 | 0.0129 | 0.1873 | 0.5579 | 0.1463 | 0 | 0 |
| Pearson | 1080 | 0.0025 | 50 | 0.237 | 0.0485 | 0.0069 | 0.1491 | 0.3246 | 0.0818 | 0 | 0 | 0.1388 | 0.0774 | 0.0109 | 0.0364 | 0.2816 | 0.0998 | 0 | 0 | 0.4816 | 0.0445 | 0.0063 | 0.4013 | 0.5524 | 0.0691 | 0 | 0 | 0.4225 | 0.1087 | 0.0154 | 0.2624 | 0.5914 | 0.1352 | 0 | 0 |
| Pearson | 1080 | 0.042 | 50 | 0.2102 | 0.0394 | 0.0056 | 0.1452 | 0.3058 | 0.0553 | 0 | 0 | 0.1485 | 0.0412 | 0.0058 | 0.0423 | 0.235 | 0.044 | 1 | 2 | 0.5454 | 0.077 | 0.0109 | 0.3706 | 0.6276 | 0.0439 | 0 | 10 | 0.4502 | 0.0904 | 0.0128 | 0.2707 | 0.6084 | 0.1374 | 0 | 0 |
| Pearson | 1080 | 0.0815 | 50 | 0.2055 | 0.0485 | 0.0069 | 0.1122 | 0.3191 | 0.0595 | 1 | 0 | 0.1363 | 0.0395 | 0.0056 | 0.0621 | 0.2083 | 0.054 | 0 | 0 | 0.5344 | 0.0808 | 0.0114 | 0.3351 | 0.6431 | 0.0706 | 0 | 8 | 0.4356 | 0.0852 | 0.0121 | 0.2684 | 0.5874 | 0.1343 | 0 | 0 |
| Pearson | 1080 | 0.121 | 50 | 0.1882 | 0.0562 | 0.0079 | 0.052 | 0.2931 | 0.0919 | 0 | 0 | 0.1403 | 0.052 | 0.0074 | 0.0291 | 0.2414 | 0.0675 | 0 | 0 | 0.5148 | 0.0791 | 0.0112 | 0.3017 | 0.6146 | 0.0751 | 0 | 5 | 0.4165 | 0.0861 | 0.0122 | 0.2491 | 0.5488 | 0.1417 | 0 | 0 |
| Pearson | 1080 | 0.1605 | 50 | 0.1843 | 0.0614 | 0.0087 | 0.0774 | 0.3003 | 0.1064 | 0 | 0 | 0.1388 | 0.056 | 0.0079 | 0.0266 | 0.2567 | 0.0846 | 0 | 0 | 0.5134 | 0.0786 | 0.0111 | 0.3179 | 0.6548 | 0.0731 | 0 | 5 | 0.4266 | 0.0831 | 0.0118 | 0.2414 | 0.5926 | 0.1304 | 0 | 0 |
| Pearson | 1080 | 0.2 | 50 | 0.1837 | 0.0714 | 0.0101 | 0.0522 | 0.3617 | 0.0968 | 0 | 0 | 0.1349 | 0.0577 | 0.0082 | -0.0199 | 0.2569 | 0.0596 | 0 | 2 | 0.4925 | 0.0797 | 0.0113 | 0.2674 | 0.5939 | 0.0947 | 0 | 2 | 0.4137 | 0.0838 | 0.0119 | 0.2223 | 0.5738 | 0.129 | 0 | 0 |
| Pearson | 2060 | 0.0025 | 50 | 0.2435 | 0.0386 | 0.0055 | 0.1683 | 0.3062 | 0.0654 | 0 | 0 | 0.146 | 0.0752 | 0.0106 | 0.0454 | 0.2872 | 0.0901 | 0 | 0 | 0.5207 | 0.036 | 0.0051 | 0.4422 | 0.5654 | 0.0478 | 0 | 0 | 0.436 | 0.1061 | 0.015 | 0.2783 | 0.596 | 0.132 | 0 | 0 |
| Pearson | 2060 | 0.042 | 50 | 0.207 | 0.0484 | 0.0068 | 0.1193 | 0.3002 | 0.0748 | 0 | 0 | 0.147 | 0.0338 | 0.0048 | 0.0649 | 0.2012 | 0.0444 | 0 | 0 | 0.5449 | 0.0811 | 0.0115 | 0.3498 | 0.6239 | 0.0355 | 0 | 10 | 0.4526 | 0.0869 | 0.0123 | 0.2823 | 0.5849 | 0.1417 | 0 | 0 |
| Pearson | 2060 | 0.0815 | 50 | 0.2035 | 0.0521 | 0.0074 | 0.1083 | 0.3149 | 0.0702 | 0 | 0 | 0.1389 | 0.0396 | 0.0056 | 0.0568 | 0.2096 | 0.0569 | 0 | 0 | 0.5336 | 0.0795 | 0.0112 | 0.3295 | 0.6383 | 0.0767 | 0 | 5 | 0.4355 | 0.0863 | 0.0122 | 0.2784 | 0.5794 | 0.1317 | 0 | 0 |
| Pearson | 2060 | 0.121 | 50 | 0.1872 | 0.0568 | 0.008 | 0.0527 | 0.2914 | 0.0873 | 0 | 0 | 0.1434 | 0.0504 | 0.0071 | 0.0271 | 0.2327 | 0.0665 | 0 | 0 | 0.5136 | 0.0782 | 0.0111 | 0.3029 | 0.6176 | 0.0703 | 0 | 5 | 0.4177 | 0.0857 | 0.0121 | 0.2515 | 0.5518 | 0.1352 | 0 | 0 |
| Pearson | 2060 | 0.1605 | 50 | 0.1829 | 0.0609 | 0.0086 | 0.0752 | 0.2819 | 0.1051 | 0 | 0 | 0.1414 | 0.0552 | 0.0078 | 0.0374 | 0.261 | 0.0856 | 0 | 0 | 0.5127 | 0.0772 | 0.0109 | 0.3202 | 0.6512 | 0.074 | 0 | 5 | 0.4261 | 0.081 | 0.0115 | 0.2462 | 0.5841 | 0.1248 | 0 | 0 |
| Pearson | 2060 | 0.2 | 50 | 0.1833 | 0.0713 | 0.0101 | 0.0499 | 0.362 | 0.0978 | 0 | 0 | 0.1361 | 0.0587 | 0.0083 | -0.0256 | 0.262 | 0.0628 | 0 | 1 | 0.493 | 0.0794 | 0.0112 | 0.2668 | 0.5949 | 0.0914 | 0 | 2 | 0.4132 | 0.0838 | 0.0119 | 0.2196 | 0.5754 | 0.1224 | 0 | 0 |
| Pearson | 3040 | 0.0025 | 50 | 0.2439 | 0.0335 | 0.0047 | 0.1717 | 0.3017 | 0.0511 | 0 | 0 | 0.1487 | 0.07 | 0.0099 | 0.0543 | 0.2737 | 0.0826 | 0 | 0 | 0.5382 | 0.0395 | 0.0056 | 0.4465 | 0.5806 | 0.0321 | 0 | 10 | 0.4433 | 0.1038 | 0.0147 | 0.2875 | 0.5954 | 0.1353 | 0 | 0 |
| Pearson | 3040 | 0.042 | 50 | 0.2065 | 0.0514 | 0.0073 | 0.1127 | 0.3043 | 0.0777 | 0 | 0 | 0.1482 | 0.0329 | 0.0047 | 0.0645 | 0.222 | 0.0527 | 0 | 0 | 0.5434 | 0.0793 | 0.0112 | 0.3539 | 0.6318 | 0.0388 | 0 | 10 | 0.4538 | 0.0859 | 0.0121 | 0.2953 | 0.5869 | 0.1555 | 0 | 0 |
| Pearson | 3040 | 0.0815 | 50 | 0.2019 | 0.0525 | 0.0074 | 0.1066 | 0.3177 | 0.0728 | 0 | 0 | 0.1416 | 0.0394 | 0.0056 | 0.0556 | 0.212 | 0.0505 | 0 | 0 | 0.5335 | 0.0782 | 0.0111 | 0.3342 | 0.6328 | 0.0707 | 0 | 7 | 0.4353 | 0.0859 | 0.0121 | 0.2789 | 0.5785 | 0.1332 | 0 | 0 |
| Pearson | 3040 | 0.121 | 50 | 0.1871 | 0.0563 | 0.008 | 0.0581 | 0.2924 | 0.0835 | 0 | 0 | 0.1443 | 0.05 | 0.0071 | 0.0293 | 0.2342 | 0.0675 | 0 | 0 | 0.5128 | 0.0778 | 0.011 | 0.3027 | 0.6207 | 0.0694 | 0 | 5 | 0.4171 | 0.0851 | 0.012 | 0.2515 | 0.5494 | 0.1364 | 0 | 0 |
| Pearson | 3040 | 0.1605 | 50 | 0.183 | 0.0604 | 0.0085 | 0.0768 | 0.2822 | 0.1041 | 0 | 0 | 0.1419 | 0.0551 | 0.0078 | 0.0368 | 0.262 | 0.0846 | 0 | 0 | 0.5124 | 0.0772 | 0.0109 | 0.3209 | 0.6511 | 0.0741 | 0 | 4 | 0.4258 | 0.0809 | 0.0114 | 0.2464 | 0.5827 | 0.1253 | 0 | 0 |
| Pearson | 3040 | 0.2 | 50 | 0.1832 | 0.0713 | 0.0101 | 0.0497 | 0.3616 | 0.0968 | 0 | 0 | 0.1364 | 0.0587 | 0.0083 | -0.0249 | 0.2622 | 0.0634 | 0 | 1 | 0.493 | 0.0793 | 0.0112 | 0.2669 | 0.5954 | 0.0908 | 0 | 2 | 0.413 | 0.0838 | 0.0119 | 0.2196 | 0.5755 | 0.1228 | 0 | 0 |
| Pearson | 4020 | 0.0025 | 50 | 0.2425 | 0.0317 | 0.0045 | 0.1826 | 0.2853 | 0.051 | 0 | 0 | 0.1498 | 0.0667 | 0.0094 | 0.0553 | 0.2696 | 0.0811 | 0 | 0 | 0.5453 | 0.044 | 0.0062 | 0.4479 | 0.5889 | 0.0173 | 0 | 10 | 0.4475 | 0.1014 | 0.0143 | 0.2938 | 0.5974 | 0.1419 | 0 | 0 |

| | | | | | | | | | | | | | | | | | | | | | | | | | | | | | | | | | | | |
|---|---|---|---|---|---|---|---|---|---|---|---|---|---|---|---|---|---|---|---|---|---|---|---|---|---|---|---|---|---|---|---|---|---|---|---|
| Pearson | 4020 | 0.042 | 50 | 0.2052 | 0.0519 | 0.0073 | 0.1076 | 0.3011 | 0.0814 | 0 | 0 | 0.1497 | 0.0324 | 0.0046 | 0.0755 | 0.2341 | 0.0466 | 0 | 0 | 0.5422 | 0.0791 | 0.0112 | 0.3505 | 0.628 | 0.0364 | 0 | 10 | 0.4529 | 0.0844 | 0.0119 | 0.2951 | 0.5838 | 0.1513 | 0 | 0 |
| Pearson | 4020 | 0.0815 | 50 | 0.2015 | 0.0522 | 0.0074 | 0.1083 | 0.3168 | 0.0713 | 0 | 0 | 0.1425 | 0.0401 | 0.0057 | 0.0556 | 0.2168 | 0.0503 | 0 | 0 | 0.5327 | 0.0781 | 0.0111 | 0.331 | 0.6317 | 0.0701 | 0 | 7 | 0.4348 | 0.0856 | 0.0121 | 0.2786 | 0.5768 | 0.1332 | 0 | 0 |
| Pearson | 4020 | 0.121 | 50 | 0.187 | 0.0562 | 0.008 | 0.0587 | 0.2914 | 0.0837 | 0 | 0 | 0.1447 | 0.0501 | 0.0071 | 0.0293 | 0.2351 | 0.0685 | 0 | 0 | 0.5126 | 0.0777 | 0.011 | 0.3032 | 0.6209 | 0.0681 | 0 | 5 | 0.417 | 0.085 | 0.012 | 0.2516 | 0.5493 | 0.1355 | 0 | 0 |
| Pearson | 4020 | 0.1605 | 50 | 0.183 | 0.0604 | 0.0085 | 0.077 | 0.2821 | 0.1039 | 0 | 0 | 0.142 | 0.0551 | 0.0078 | 0.0371 | 0.2623 | 0.0845 | 0 | 0 | 0.5123 | 0.0771 | 0.0109 | 0.321 | 0.6511 | 0.074 | 0 | 4 | 0.4258 | 0.0809 | 0.0114 | 0.2465 | 0.5824 | 0.125 | 0 | 0 |
| Pearson | 4020 | 0.2 | 50 | 0.1832 | 0.0712 | 0.0101 | 0.0497 | 0.3615 | 0.0967 | 0 | 0 | 0.1365 | 0.0587 | 0.0083 | -0.0249 | 0.2623 | 0.0634 | 0 | 1 | 0.493 | 0.0793 | 0.0112 | 0.2668 | 0.5954 | 0.0907 | 0 | 2 | 0.413 | 0.0838 | 0.0118 | 0.2196 | 0.5754 | 0.1227 | 0 | 0 |
| Pearson | 5000 | 0.0025 | 50 | 0.24 | 0.0319 | 0.0045 | 0.1802 | 0.281 | 0.0546 | 0 | 0 | 0.1504 | 0.0634 | 0.009 | 0.0579 | 0.261 | 0.072 | 0 | 0 | 0.5497 | 0.0489 | 0.0069 | 0.4463 | 0.5996 | 0.0161 | 0 | 10 | 0.4502 | 0.1002 | 0.0142 | 0.2966 | 0.5963 | 0.1407 | 0 | 0 |
| Pearson | 5000 | 0.042 | 50 | 0.2043 | 0.0522 | 0.0074 | 0.1033 | 0.2988 | 0.0856 | 0 | 0 | 0.1506 | 0.0324 | 0.0046 | 0.0744 | 0.235 | 0.0466 | 0 | 0 | 0.5414 | 0.0783 | 0.0111 | 0.3536 | 0.6255 | 0.036 | 0 | 10 | 0.4521 | 0.0839 | 0.0119 | 0.2984 | 0.5803 | 0.1498 | 0 | 0 |
| Pearson | 5000 | 0.0815 | 50 | 0.2014 | 0.0521 | 0.0074 | 0.1084 | 0.317 | 0.0699 | 0 | 0 | 0.143 | 0.0403 | 0.0057 | 0.0554 | 0.218 | 0.0512 | 0 | 0 | 0.5324 | 0.0779 | 0.011 | 0.3313 | 0.6301 | 0.0689 | 0 | 8 | 0.4345 | 0.0855 | 0.0121 | 0.2791 | 0.5761 | 0.1329 | 0 | 0 |
| Pearson | 5000 | 0.121 | 50 | 0.187 | 0.0562 | 0.0079 | 0.0592 | 0.2913 | 0.0839 | 0 | 0 | 0.1448 | 0.0502 | 0.0071 | 0.0294 | 0.2357 | 0.0682 | 0 | 0 | 0.5125 | 0.0777 | 0.011 | 0.3033 | 0.6209 | 0.068 | 0 | 5 | 0.417 | 0.085 | 0.012 | 0.2516 | 0.549 | 0.1355 | 0 | 0 |
| Pearson | 5000 | 0.1605 | 50 | 0.183 | 0.0603 | 0.0085 | 0.0771 | 0.2821 | 0.1038 | 0 | 0 | 0.1421 | 0.0551 | 0.0078 | 0.0371 | 0.2623 | 0.0845 | 0 | 0 | 0.5123 | 0.0771 | 0.0109 | 0.321 | 0.6511 | 0.074 | 0 | 4 | 0.4258 | 0.0809 | 0.0114 | 0.2465 | 0.5824 | 0.125 | 0 | 0 |
| Pearson | 5000 | 0.2 | 50 | 0.1832 | 0.0712 | 0.0101 | 0.0497 | 0.3615 | 0.0966 | 0 | 0 | 0.1365 | 0.0587 | 0.0083 | -0.0249 | 0.2623 | 0.0634 | 0 | 1 | 0.493 | 0.0793 | 0.0112 | 0.2668 | 0.5954 | 0.0906 | 0 | 2 | 0.413 | 0.0838 | 0.0118 | 0.2196 | 0.5754 | 0.1227 | 0 | 0 |
| R_squared | 100 | 0.0025 | 50 | -22.2938 | 3.9305 | 0.5559 | -29.7385 | -19.0835 | 2.9028 | 0 | 10 | -0.0685 | 0.0534 | 0.0076 | -0.1361 | -0.0001 | 0.1 | 0 | 0 | -0.1282 | 0.0701 | 0.0099 | -0.2622 | -0.0608 | 0.0506 | 0 | 10 | -0.0398 | 0.051 | 0.0072 | -0.1005 | 0.0468 | 0.0722 | 0 | 0 |
| R_squared | 100 | 0.042 | 50 | -22.1912 | 3.8879 | 0.5498 | -29.7607 | -18.7808 | 2.884 | 0 | 10 | -0.0609 | 0.0619 | 0.0087 | -0.1695 | 0.0293 | 0.1142 | 0 | 0 | 0.0577 | 0.0534 | 0.0076 | -0.0485 | 0.1477 | 0.0545 | 0 | 1 | 0.0616 | 0.0853 | 0.0121 | -0.0485 | 0.2523 | 0.0567 | 10 | 0 |
| R_squared | 100 | 0.0815 | 50 | -22.1698 | 3.8969 | 0.5511 | -29.8261 | -18.7416 | 2.9049 | 0 | 10 | -0.0774 | 0.0653 | 0.0092 | -0.2384 | 0.0358 | 0.1133 | 0 | 0 | 0.1069 | 0.0515 | 0.0073 | -0.0247 | 0.195 | 0.064 | 0 | 2 | 0.0681 | 0.1021 | 0.0144 | -0.0869 | 0.2976 | 0.1043 | 4 | 0 |
| R_squared | 100 | 0.121 | 50 | -22.21 | 3.9067 | 0.5525 | -29.9845 | -18.5413 | 2.9519 | 0 | 10 | -0.0996 | 0.0657 | 0.0093 | -0.2471 | 0.0211 | 0.0917 | 0 | 0 | 0.1172 | 0.0615 | 0.0087 | -0.0463 | 0.2381 | 0.0961 | 0 | 0 | 0.0472 | 0.1205 | 0.017 | -0.1499 | 0.2952 | 0.17 | 0 | 0 |
| R_squared | 100 | 0.1605 | 50 | -22.2811 | 3.9893 | 0.5642 | -30.4897 | -18.3976 | 2.9388 | 0 | 10 | -0.1263 | 0.0744 | 0.0105 | -0.339 | 0.0136 | 0.0915 | 0 | 1 | 0.1186 | 0.068 | 0.0096 | -0.0608 | 0.2372 | 0.0819 | 0 | 2 | 0.0356 | 0.124 | 0.0175 | -0.178 | 0.2968 | 0.1767 | 0 | 0 |
| R_squared | 100 | 0.2 | 50 | -22.3503 | 3.968 | 0.5612 | -30.3328 | -18.5943 | 3.0354 | 0 | 10 | -0.1658 | 0.0772 | 0.0109 | -0.3495 | 0.0218 | 0.1131 | 0 | 0 | 0.1233 | 0.0792 | 0.0112 | -0.0865 | 0.2685 | 0.0995 | 0 | 2 | 0.0085 | 0.1295 | 0.0183 | -0.2208 | 0.2862 | 0.1759 | 0 | 0 |
| R_squared | 1080 | 0.0025 | 50 | -22.2011 | 3.8781 | 0.5484 | -29.597 | -19.022 | 2.7792 | 0 | 10 | -0.0567 | 0.0591 | 0.0084 | -0.1358 | 0.018 | 0.1066 | 0 | 0 | 0.019 | 0.0556 | 0.0079 | -0.0831 | 0.0994 | 0.051 | 0 | 2 | 0.0529 | 0.0758 | 0.0107 | -0.0173 | 0.2049 | 0.0118 | 10 | 10 |
| R_squared | 1080 | 0.042 | 50 | -22.3182 | 3.9138 | 0.5535 | -29.8163 | -18.8459 | 2.9583 | 0 | 10 | -0.1535 | 0.0536 | 0.0076 | -0.2626 | -0.0643 | 0.0782 | 0 | 0 | 0.1843 | 0.0702 | 0.0099 | 0.0461 | 0.2915 | 0.1085 | 0 | 0 | 0.0853 | 0.1413 | 0.02 | -0.1281 | 0.3482 | 0.2046 | 0 | 0 |
| R_squared | 1080 | 0.0815 | 50 | -22.4572 | 3.9764 | 0.5623 | -30.3376 | -18.8835 | 3.0542 | 0 | 10 | -0.2227 | 0.0527 | 0.0075 | -0.3405 | -0.1152 | 0.065 | 0 | 0 | 0.1707 | 0.0841 | 0.0119 | -0.0109 | 0.2872 | 0.1126 | 0 | 0 | 0.0278 | 0.1489 | 0.0211 | -0.1987 | 0.3071 | 0.2525 | 0 | 0 |
| R_squared | 1080 | 0.121 | 50 | -22.5472 | 4.0013 | 0.5659 | -30.6509 | -18.7263 | 3.162 | 0 | 10 | -0.2571 | 0.0705 | 0.01 | -0.3937 | -0.1203 | 0.1148 | 0 | 0 | 0.1366 | 0.09 | 0.0127 | -0.0865 | 0.2648 | 0.1331 | 0 | 0 | -0.0204 | 0.1475 | 0.0209 | -0.2729 | 0.2232 | 0.2659 | 0 | 0 |
| R_squared | 1080 | 0.1605 | 50 | -22.6293 | 4.0378 | 0.571 | -31.0848 | -18.9255 | 3.0719 | 0 | 10 | -0.2941 | 0.0817 | 0.0116 | -0.4697 | -0.1102 | 0.0859 | 2 | 0 | 0.126 | 0.0888 | 0.0126 | -0.0475 | 0.326 | 0.1172 | 0 | 0 | -0.0319 | 0.16 | 0.0226 | -0.3635 | 0.2907 | 0.2512 | 0 | 0 |
| R_squared | 1080 | 0.2 | 50 | -22.6321 | 4.0597 | 0.5741 | -31.306 | -18.6856 | 3.3146 | 0 | 10 | -0.3209 | 0.075 | 0.0106 | -0.4854 | -0.1757 | 0.088 | 0 | 0 | 0.0911 | 0.0919 | 0.013 | -0.1148 | 0.2381 | 0.1442 | 0 | 0 | -0.0632 | 0.1543 | 0.0218 | -0.4177 | 0.2137 | 0.2636 | 0 | 0 |
| R_squared | 2060 | 0.0025 | 50 | -22.1904 | 3.8676 | 0.547 | -29.5878 | -18.9755 | 2.7263 | 0 | 10 | -0.0562 | 0.0602 | 0.0085 | -0.1397 | 0.0169 | 0.1204 | 0 | 0 | 0.0744 | 0.0483 | 0.0068 | -0.0112 | 0.152 | 0.0346 | 6 | 9 | 0.078 | 0.0881 | 0.0125 | -0.0101 | 0.2488 | 0.0558 | 10 | 0 |
| R_squared | 2060 | 0.042 | 50 | -22.4284 | 3.9218 | 0.5546 | -30.0072 | -19.0211 | 2.8875 | 0 | 10 | -0.1973 | 0.0444 | 0.0063 | -0.2965 | -0.1196 | 0.0685 | 0 | 0 | 0.1835 | 0.0767 | 0.0109 | 0.0072 | 0.2838 | 0.1037 | 0 | 0 | 0.0631 | 0.146 | 0.0206 | -0.1703 | 0.3075 | 0.2378 | 0 | 0 |
| R_squared | 2060 | 0.0815 | 50 | -22.5288 | 4.0286 | 0.5697 | -30.6907 | -18.9642 | 3.0631 | 0 | 10 | -0.2518 | 0.0518 | 0.0073 | -0.3664 | -0.1402 | 0.0667 | 0 | 0 | 0.1645 | 0.0838 | 0.0118 | -0.0318 | 0.2783 | 0.1089 | 0 | 0 | 0.0081 | 0.1543 | 0.0218 | -0.223 | 0.2917 | 0.26 | 0 | 0 |
| R_squared | 2060 | 0.121 | 50 | -22.5933 | 4.0305 | 0.57 | -30.69 | -18.7552 | 3.0924 | 0 | 10 | -0.2707 | 0.0662 | 0.0094 | -0.3941 | -0.1388 | 0.1032 | 0 | 0 | 0.1304 | 0.0873 | 0.0123 | -0.0962 | 0.2507 | 0.133 | 0 | 0 | -0.0303 | 0.1478 | 0.0209 | -0.2821 | 0.2089 | 0.2653 | 0 | 0 |
| R_squared | 2060 | 0.1605 | 50 | -22.6545 | 4.0615 | 0.5744 | -31.1747 | -19.0362 | 3.1406 | 0 | 10 | -0.3017 | 0.0791 | 0.0112 | -0.469 | -0.1224 | 0.0849 | 1 | 0 | 0.1226 | 0.087 | 0.0123 | -0.0514 | 0.3175 | 0.1128 | 0 | 0 | -0.039 | 0.1565 | 0.0221 | -0.3676 | 0.268 | 0.2531 | 0 | 0 |
| R_squared | 2060 | 0.2 | 50 | -22.6446 | 4.0761 | 0.5764 | -31.3869 | -18.6906 | 3.2892 | 0 | 10 | -0.3246 | 0.0751 | 0.0106 | -0.4938 | -0.1829 | 0.0847 | 0 | 1 | 0.0898 | 0.091 | 0.0129 | -0.1157 | 0.2376 | 0.1412 | 0 | 0 | -0.0679 | 0.1542 | 0.0218 | -0.434 | 0.2114 | 0.2554 | 0 | 0 |
| R_squared | 3040 | 0.0025 | 50 | -22.1775 | 3.8684 | 0.5471 | -29.6056 | -18.9803 | 2.714 | 0 | 10 | -0.0595 | 0.0603 | 0.0085 | -0.1444 | 0.0127 | 0.1213 | 0 | 0 | 0.1068 | 0.0442 | 0.0063 | 0.0313 | 0.1799 | 0.0356 | 0 | 2 | 0.0914 | 0.0977 | 0.0138 | -0.0155 | 0.274 | 0.089 | 10 | 0 |
| R_squared | 3040 | 0.042 | 50 | -22.4724 | 3.9576 | 0.5597 | -30.1448 | -19.0708 | 2.8207 | 0 | 10 | -0.2159 | 0.0421 | 0.0059 | -0.3056 | -0.1186 | 0.0595 | 0 | 0 | 0.1789 | 0.0755 | 0.0107 | 0.0057 | 0.2729 | 0.1008 | 0 | 0 | 0.0542 | 0.1485 | 0.021 | -0.1853 | 0.3034 | 0.2523 | 0 | 0 |
| R_squared | 3040 | 0.0815 | 50 | -22.5489 | 4.0538 | 0.5733 | -30.8093 | -19.0268 | 2.9968 | 0 | 10 | -0.2579 | 0.0509 | 0.0072 | -0.378 | -0.1439 | 0.0619 | 0 | 0 | 0.1621 | 0.0826 | 0.0117 | -0.0305 | 0.2754 | 0.1119 | 0 | 0 | 0.003 | 0.1539 | 0.0218 | -0.2288 | 0.2842 | 0.2629 | 0 | 0 |
| R_squared | 3040 | 0.121 | 50 | -22.6023 | 4.0363 | 0.5708 | -30.7123 | -18.7904 | 3.0629 | 0 | 10 | -0.2729 | 0.0658 | 0.0093 | -0.3992 | -0.141 | 0.1036 | 0 | 0 | 0.1284 | 0.0865 | 0.0122 | -0.0985 | 0.2465 | 0.1315 | 0 | 0 | -0.033 | 0.1469 | 0.0208 | -0.2815 | 0.2003 | 0.2696 | 0 | 0 |
| R_squared | 3040 | 0.1605 | 50 | -22.6608 | 4.0669 | 0.5751 | -31.1942 | -19.0591 | 3.1415 | 0 | 10 | -0.3024 | 0.0786 | 0.0111 | -0.4722 | -0.1224 | 0.084 | 1 | 1 | 0.1219 | 0.0869 | 0.0123 | -0.0506 | 0.3176 | 0.1129 | 0 | 0 | -0.0402 | 0.1563 | 0.0221 | -0.3677 | 0.2648 | 0.2502 | 0 | 0 |
| R_squared | 3040 | 0.2 | 50 | -22.647 | 4.0791 | 0.5769 | -31.3969 | -18.697 | 3.2891 | 0 | 10 | -0.3246 | 0.0751 | 0.0106 | -0.4953 | -0.1836 | 0.0839 | 0 | 1 | 0.0896 | 0.0908 | 0.0128 | -0.115 | 0.2383 | 0.1413 | 0 | 0 | -0.0684 | 0.1542 | 0.0218 | -0.4352 | 0.2116 | 0.2547 | 0 | 0 |
| R_squared | 4020 | 0.0025 | 50 | -22.1816 | 3.8718 | 0.5476 | -29.6129 | -18.9605 | 2.7509 | 0 | 10 | -0.0646 | 0.0608 | 0.0086 | -0.1521 | 0.0065 | 0.125 | 0 | 0 | 0.1271 | 0.0426 | 0.006 | 0.0593 | 0.193 | 0.0521 | 0 | 0 | 0.0995 | 0.1049 | 0.0148 | -0.0139 | 0.2926 | 0.1152 | 0 | 0 |
| R_squared | 4020 | 0.042 | 50 | -22.5084 | 3.9895 | 0.5642 | -30.2253 | -19.0812 | 2.8393 | 0 | 10 | -0.2246 | 0.0412 | 0.0058 | -0.3106 | -0.1236 | 0.0518 | 0 | 0 | 0.1755 | 0.0753 | 0.0107 | -0.0027 | 0.2737 | 0.0987 | 0 | 0 | 0.0477 | 0.1475 | 0.0209 | -0.1942 | 0.2918 | 0.2474 | 0 | 0 |
| R_squared | 4020 | 0.0815 | 50 | -22.5595 | 4.0608 | 0.5743 | -30.8464 | -19.0279 | 2.9789 | 0 | 10 | -0.2597 | 0.0511 | 0.0072 | -0.3792 | -0.1461 | 0.0644 | 0 | 0 | 0.1604 | 0.0824 | 0.0116 | -0.0353 | 0.2733 | 0.1102 | 0 | 0 | 0.0004 | 0.1535 | 0.0217 | -0.2316 | 0.2786 | 0.2619 | 0 | 0 |
| R_squared | 4020 | 0.121 | 50 | -22.6055 | 4.0389 | 0.5712 | -30.727 | -18.7989 | 3.0597 | 0 | 10 | -0.2731 | 0.0659 | 0.0093 | -0.401 | -0.1411 | 0.103 | 0 | 0 | 0.128 | 0.0863 | 0.0122 | -0.0984 | 0.2466 | 0.1307 | 0 | 0 | -0.0336 | 0.1467 | 0.0207 | -0.2817 | 0.1987 | 0.2684 | 0 | 0 |
| R_squared | 4020 | 0.1605 | 50 | -22.6615 | 4.0676 | 0.5752 | -31.1988 | -19.0636 | 3.1439 | 0 | 10 | -0.3024 | 0.0786 | 0.0111 | -0.4722 | -0.1224 | 0.0843 | 1 | 1 | 0.1218 | 0.0869 | 0.0123 | -0.0508 | 0.3177 | 0.1127 | 0 | 0 | -0.0404 | 0.1562 | 0.0221 | -0.368 | 0.2642 | 0.2497 | 0 | 0 |
| R_squared | 4020 | 0.2 | 50 | -22.6471 | 4.0792 | 0.5769 | -31.3974 | -18.6974 | 3.2891 | 0 | 10 | -0.3246 | 0.0751 | 0.0106 | -0.4952 | -0.1835 | 0.0839 | 0 | 1 | 0.0895 | 0.0908 | 0.0128 | -0.1149 | 0.2384 | 0.1412 | 0 | 0 | -0.0684 | 0.1542 | 0.0218 | -0.4352 | 0.2116 | 0.2547 | 0 | 0 |
| R_squared | 5000 | 0.0025 | 50 | -22.1844 | 3.8822 | 0.549 | -29.6443 | -18.9489 | 2.7626 | 0 | 10 | -0.0706 | 0.0605 | 0.0086 | -0.1553 | 0.002 | 0.1216 | 0 | 0 | 0.1424 | 0.0422 | 0.006 | 0.0826 | 0.2066 | 0.0679 | 0 | 0 | 0.1044 | 0.1115 | 0.0158 | -0.018 | 0.3037 | 0.1341 | 0 | 0 |
| R_squared | 5000 | 0.042 | 50 | -22.5294 | 4.0121 | 0.5674 | -30.31 | -19.1174 | 2.7906 | 0 | 10 | -0.2292 | 0.0415 | 0.0059 | -0.3147 | -0.1269 | 0.0523 | 0 | 0 | 0.1737 | 0.0742 | 0.0105 | -0.0011 | 0.2695 | 0.0942 | 0 | 0 | 0.0436 | 0.1471 | 0.0208 | -0.1978 | 0.2826 | 0.2432 | 0 | 0 |
| R_squared | 5000 | 0.0815 | 50 | -22.563 | 4.0661 | 0.575 | -30.8683 | -19.0287 | 2.9644 | 0 | 10 | -0.26 | 0.0511 | 0.0072 | -0.38 | -0.1454 | 0.0652 | 0 | 0 | 0.1597 | 0.082 | 0.0116 | -0.0356 | 0.2729 | 0.1094 | 0 | 0 | -0.0006 | 0.1535 | 0.0217 | -0.233 | 0.2767 | 0.262 | 0 | 0 |
| R_squared | 5000 | 0.121 | 50 | -22.6064 | 4.0396 | 0.5713 | -30.7298 | -18.8015 | 3.0601 | 0 | 10 | -0.2732 | 0.0659 | 0.0093 | -0.4011 | -0.1418 | 0.1033 | 0 | 0 | 0.1278 | 0.0863 | 0.0122 | -0.0983 | 0.2466 | 0.1306 | 0 | 0 | -0.0338 | 0.1467 | 0.0208 | -0.282 | 0.1985 | 0.2684 | 0 | 0 |
| R_squared | 5000 | 0.1605 | 50 | -22.6616 | 4.0678 | 0.5753 | -31.1998 | -19.0638 | 3.1443 | 0 | 10 | -0.3024 | 0.0785 | 0.0111 | -0.4721 | -0.1223 | 0.0843 | 1 | 1 | 0.1218 | 0.0868 | 0.0123 | -0.0508 | 0.3176 | 0.1127 | 0 | 0 | -0.0404 | 0.1562 | 0.0221 | -0.3681 | 0.2641 | 0.2496 | 0 | 0 |
| R_squared | 5000 | 0.2 | 50 | -22.6471 | 4.0792 | 0.5769 | -31.3976 | -18.6975 | 3.289 | 0 | 10 | -0.3246 | 0.0751 | 0.0106 | -0.4952 | -0.1835 | 0.0839 | 0 | 1 | 0.0895 | 0.0908 | 0.0128 | -0.1149 | 0.2384 | 0.1412 | 0 | 0 | -0.0684 | 0.1542 | 0.0218 | -0.4352 | 0.2116 | 0.2547 | 0 | 0 |
